# Sexual dimorphism in the complete connectome of the *Drosophila* male central nervous system

**DOI:** 10.1101/2025.10.09.680999

**Authors:** Stuart Berg, Isabella R Beckett, Marta Costa, Philipp Schlegel, Michał Januszewski, Elizabeth C Marin, Aljoscha Nern, Stephan Preibisch, Wei Qiu, Shin-ya Takemura, Alexandra MC Fragniere, Andrew S Champion, Diane-Yayra Adjavon, Michael Cook, Marina Gkantia, Kenneth J Hayworth, Gary B Huang, William T Katz, Florian Kämpf, Zhiyuan Lu, Christopher Ordish, Tyler Paterson, Tomke Stürner, Eric T Trautman, Catherine R Whittle, Laura E Burnett, Judith Hoeller, Feng Li, Frank Loesche, Billy J Morris, Tobias Pietzsch, Markus W Pleijzier, Valeria Silva, Yijie Yin, Iris Ali, Griffin Badalamente, Alexander Shakeel Bates, Rory J Beresford, John Bogovic, Paul Brooks, Sebastian Cachero, Brandon S Canino, Bhumpanya Chaisrisawatsuk, Jody Clements, Arthur Crowe, Inês de Haan Vicente, Georgia Dempsey, Erika Donà, Márcia dos Santos, Marisa Dreher, Christopher R Dunne, Katharina Eichler, Samantha Finley-May, Miriam A Flynn, Imran Hameed, Gary Patrick Hopkins, Philip M Hubbard, Ladann Kiassat, Julie Kovalyak, Shirley A Lauchie, Meghan Leonard, Alanna Lohff, Kit D Longden, Charli A Maldonado, Ilina Moitra, Sung Soo Moon, Caroline Mooney, Eva J Munnelly, Nneoma Okeoma, Donald J Olbris, Anika Pai, Birava Patel, Emily M Phillips, Stephen M Plaza, Alana Richards, Jennifer Rivas Salinas, Ruairí JV Roberts, Edward M Rogers, Ashley L Scott, Louis A Scuderi, Pavithraa Seenivasan, Laia Serratosa Capdevila, Claire Smith, Rob Svirskas, Satoko Takemura, Ibrahim Tastekin, Alexander Thomson, Lowell Umayam, John J Walsh, Holly Whittome, C Shan Xu, Emily A Yakal, Tansy Yang, Arthur Zhao, Reed George, Viren Jain, Vivek Jayaraman, Wyatt Korff, Geoffrey W Meissner, Sandro Romani, Jan Funke, Christopher Knecht, Stephan Saalfeld, Louis K Scheffer, Scott Waddell, Gwyneth M Card, Carlos Ribeiro, Michael B Reiser, Harald F Hess, Gerald M Rubin, Gregory SXE Jefferis

**Affiliations:** Janelia Research Campus, Howard Hughes Medical Institute, Ashburn, VA, USA; Janelia FlyEM Project Team; Neurobiology Division, MRC Laboratory of Molecular Biology, Cambridge, UK; Department of Zoology, University of Cambridge, UK; Google Research, Zürich, Switzerland; Department of Physiology, Development and Neuroscience, University of Cambridge, UK; Centre for Neural Circuits and Behaviour, University of Oxford, UK; Institute of Neuroscience, Consiglio Nazionale delle Ricerche (CNR), Vedano al Lambro (MB), Italy; Champalimaud Foundation, Lisbon, Portugal; Zuckerman Institute, Columbia University, New York, NY, USA; Google Research, Mountain View, CA, USA; Department of Neurobiology, Harvard Medical School, Boston, MA, USA; Aelysia Ltd., Bristol, UK

## Abstract

Sex differences in behaviour exist across all animals, typically under strong genetic regulation. In *Drosophila*, *fruitless*/*doublesex* transcription factors can identify dimorphic neurons but their organisation into functional circuits remains unclear.

We present the connectome of the entire *Drosophila* male central nervous system. This contains 166,691 neurons spanning the brain and nerve cord, fully proofread and annotated including *fruitless*/*doublesex* expression and 11,691 types. We provide the first comprehensive comparison between male and female brain connectomes to synaptic resolution, finding 7,205 isomorphic, 114 dimorphic, 262 male-specific and 69 female-specific types.

This resource enables analysis of full sensory-to-motor circuits underlying complex behaviours and the impact of dimorphic elements. Sex-specific/dimorphic neurons are concentrated in higher brain centres while the sensory and motor periphery are largely isomorphic. Within higher centres, male-specific connections are organised into hotspots defined by male-specific neurons or arbours. Numerous circuit switches reroute sensory information to form antagonistic circuits controlling opposing behaviours.

## Introduction

To obtain a detailed understanding of how brains transform sensory information and memories into motor actions, we must define the components of neural circuits, understand how they fit together, and investigate the functional consequences of manipulating specific circuit elements. Sexual dimorphisms provide a natural experimental framework for causal inference: when the same sensory stimulus evokes different behaviours in males and females of a species, dissecting the underlying neural circuits offers insights into how the brain produces behaviour. For example, virgin male and female mice respond differently to the presence of pups: while females parent, males are infanticidal. These antagonistic behaviours are controlled by a sexually dimorphic population of inhibitory neurons in the medial amygdala^1^. While such studies in rodents^2–8^ have identified brain regions and broad cell populations implicated in dimorphic behaviours, mapping the entire underlying circuit architecture remains intractable in such a numerically complex system.

The adult fruit fly, *Drosophila melanogaster,* is a powerful model for studying sexual dimorphism at the levels of genes, circuits, and behaviour. Dimorphic behaviours such as courtship and aggression depend on complex sensorimotor integration. Although genetically specified by the transcription factors *fruitless* and *doublesex*^9–11^, they can be modified by experience^12^. Powerful molecular genetic toolkits have enabled access to specific cell types within broad *fruitless* and *doublesex*-expressing neuronal populations, making it possible to dissect their contributions to numerous sexually dimorphic behaviours^13–17^.

Until now, studies of sexual dimorphism in *Drosophila* have focused on the impact of individual cell types on physiology and behaviour^18–20^. For example, male-specific P1 neurons, a diverse population which express *doublesex* and in some cases also *fruitless*, promote courtship song^19,21–23^ and aggression^24^. P1 neurons appear to be a key node controlling sex-specific behaviour with strong parallels to mammals^25^, but the circuit architecture that allows them to promote mutually exclusive behaviours remains elusive. In a few other cases it has been possible to pinpoint circuit nodes that redirect sensory information to opposing downstream circuitry^26,27^ but this remains the exception.

Whole-brain connectomics provides a systematic approach to address this circuit gap by placing individual neurons into complete synaptic resolution circuits. Connectomics has already proven useful for investigating neural circuit mechanisms of a range of complex behaviours in *Drosophila*^28^ even including cognitive processes that are individual to an organism or dynamic such as memory or the interaction between visual attention and internal state^29^. Connectomes of the fly brain and nerve cord have recently become available^30–35^, enabling dissection of neural circuits with single-neuron resolution.

Building on this foundation, we present the first finished connectome of an entire male *Drosophila* central nervous system. This provides two critical advances (1) a fully proofread and annotated brain and nerve cord connectome with an intact neck connective; and (2) the first example of a male brain. Leveraging existing female connectomes, this resource enables the first brainwide, synaptic-resolution comparison across sexes of an adult animal with complex anatomy and behaviour. We identified 331 sex-specific and 114 sexually dimorphic cell types, comprising 4.8% of the male and 2.4% of the female central brain. We found a high correspondence between dimorphism status and *fruitless*/*doublesex* expression, but these are not completely overlapping. Sex-specific and dimorphic neurons are concentrated in higher order brain centres while the sensory and motor periphery is largely isomorphic. Although only a small fraction of all neurons, dimorphism propagates through the nervous system via dimorphic connectivity, suggesting that even modest changes in circuitry can exert brain-wide influence. Comparative connectomics of circuits for multiple sensory and behavioural modules uncovers general principles of the neural architecture for sex-shared, sex-specific, and flexible behaviours. This resource enables exploration of neural circuits spanning the entire CNS and comparison of these circuits across sexes to elucidate the circuit-level impact of sexual dimorphism.

## Results

### A complete connectome of a whole male CNS

We present the fully proofread connectome of the central nervous system (CNS) of an adult *Drosophila melanogaster* male, extending the recently reported right optic lobe connectome^36^. Using 7 enhanced focused ion beam scanning electron microscopy (eFIB-SEM) systems^37,38^ over 13 months, we generated an image volume of 160 teravoxels at 8×8×8 nm isotropic resolution (0.082 mm^3^ total volume) encompassing the central brain, optic lobes, and ventral nerve cord (VNC, insect spinal cord analogue) (Fig 1a). The volume was aligned and neurons were automatically segmented with flood filling networks^39^. 46 million presynapses connected to 312 million PSDs were automatically detected with an average precision/recall of 0.82/0.81, respectively (Fig S1i, <u>Methods</u>). We extracted additional features including nuclei and neurotransmitter predictions^40^.

**Figure 1.**
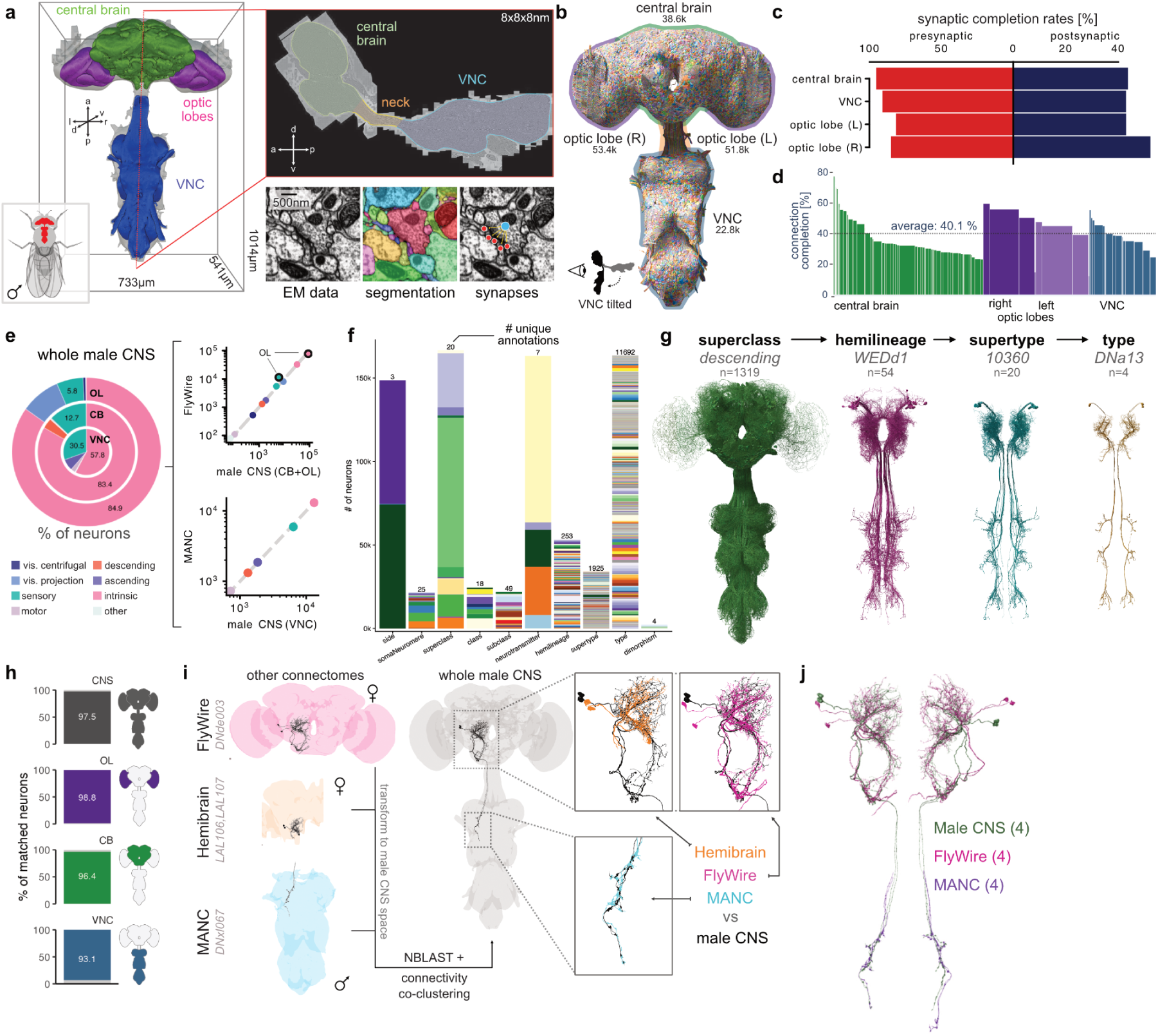
A densely annotated and cross-matched male CNS connectome. **a** Outlines of the aligned male CNS volume, coloured by CNS region (left). Sagittal slice through the EM image data (top right). Insets show (from left): image data, neuron segmentation, and synapse detection. **b** 3D rendering of all neurons in the connectome. Frontal view; VNC tilted down for visualisation. Numbers represent neuron counts per brain region, including sensory neurons. **c** Synaptic completion rates for pre- and postsynapses per brain region. **d** Percentage of connections where both pre- and postsynaptic partners are proofread broken down by neuropil region. Bar width corresponds to size of neuropil. **e** Percentages of neurons for main superclasses across brain regions (left). Comparison of neuron counts to existing connectomes: female brain (FAFB/FlyWire, top right) and male nerve cord (MANC, bottom right). **f** Number of neurons per annotation field. The number of unique annotations for each field is shown at the top of the bar. `side` combines fields `somaSide` and `rootSide`; `hemilineage` combines fields `itoleeHl` and `trumanHl`. **g** Example of the hierarchical annotation of neurons, from superclass through hemilineage, supertype and type. The annotation and the number of neurons in that category is shown in grey. **h** Percentage of neurons matched to an existing dataset in the whole male CNS and for each CNS region. **i** Neurons were matched to existing connectomes using a combination of spatial transforms + NBLAST and connectivity co-clustering. Example shown here is DNde003, a descending neuron type, with a 1:1 match in FlyWire and MANC, and matched to two types in hemibrain. **j** Full view of DNde003 in male CNS, FlyWire and MANC. The number of neurons in each dataset is shown in brackets. EM: electron microscopy; OL: optic lobes; CB: central brain; MANC: male adult nerve cord connectome; VNC: ventral nerve cord.

The initial neuron segmentation was proofread with an estimated effort of 44 person-years (Fig S9, <u>Methods</u>). In total, we identified, proofread and annotated 166,691 neurons (including sensory axons) across the entire volume (Fig 1b). While absolute certainty is difficult to achieve in datasets as large as this, we are confident that these numbers represent essentially all traceable neurons. First, we proofread all fragments from the initial segmentation with > 100 synaptic connections. Second, 98.9% of the 141,780 detected neuron-associated nuclei are part of a proofread neuron; of the remaining 1.1% we estimate that >99% were proofread by the first strategy. Third, the resulting neuron counts match those reported for existing brain and VNC connectomes (Fig 1e). A small number of cells could not be reconstructed due to sample artefacts on the edge of the volume (e.g. some R1-6 photoreceptors neurons in the laminae) or segmentation issues (some sensory and motor neurons) (see <u>Methods</u> for details).

Connectome completeness is typically assessed by the fraction of pre- and postsynapses associated with proofread neurons; with 94% pre- and 42% postsynaptic completion rates in neuropils, this dataset is similar to or exceeds previously published connectomes^30–34^ (Fig 1c, Fig S9g). Going forwards we suggest that the fraction of synaptic connections for which both pre- <u>and</u> postsynaptic sites belong to a proofread neuron is more useful. For the male CNS connectome, that fraction is 40.1% (Fig 1d). The neuron segmentation and synaptic connections jointly define a connectome graph containing 25.6M edges between 166,391 neurons.

To maximise utility for the wider biological community, we provide comprehensive annotations detailing the organization and structure of the entire *Drosophila* central nervous system. These are informed by and cross-referenced with recent partial connectome datasets. The coarsest annotation, “`superclass`”, describes the direction of information flow, anatomical location and broad function of neurons. Orthogonal annotations include developmental origin (`hemilineage`), spatial location (`somaSide`, `somaNeuromere`, `entry/exitNerve`), putative function (`class` and `subclass`) and names used in the literature (`synonyms`) (Fig 1f and Fig S1c). At the most granular level, we defined 11,691 unique cell types based on morphology and connectivity of individual neurons across the CNS (Fig 1g).

To enable across-dataset comparisons, we used NBLAST^41^ morphology scores and connectivity similarity to systematically match and annotate cell types across existing connectomes (see <u>Methods</u> for details) (Fig 1h-j). 97.5% of neurons across the CNS have a cell type match to either the FAFB/FlyWire^30,31,42^, hemibrain^32^ and/or male adult nerve cord (MANC)^32–34^ datasets (Fig 1h and Fig S1e). Of the remaining 2.5%, some could not be matched due to technical limitations of previous partial datasets (Fig S1f, <u>Methods</u>); male-specific neurons are presented <u>later</u>.

We matched cell types across connectomes with high fidelity, but nomenclature is not yet consistent. We therefore recorded parallel type annotations for other datasets (`mancType`, `flywireType`, `hemibrainType`) so that the corpus of male CNS annotations now serve as a connectomic rosetta stone. We occasionally revised existing cell types in FAFB/FlyWire, hemibrain and/or MANC when whole-CNS information enabled more accurate cell typing (Fig S1g and see <u>Methods</u> for downloadable resources). Reassuringly, we see that cell type definitions are converging: we revised about 4% of FlyWire neurons in this work (Fig S1h) whereas in earlier work matching the hemibrain to FlyWire we had to revise over 44% of cell types^31^. The male CNS therefore represents an accurate and durable consensus cell type atlas.

Taken together, this high quality and extensively annotated connectome should be the principal reference for researchers exploring the anatomy, connectivity, and development across the *Drosophila* CNS, and for unraveling the structure-function relationships linking circuits to behaviour. A <u>landing page</u> brings together all resources (images, segmentation, neuron reconstructions, connectome graph, neurotransmitter predictions, annotations, etc.), with interactive tools to explore cell types, connectivity and male-female dimorphism, and raw data for download (see also <u>Methods</u>).

### Information flow from sensory to motor

Using a limited repertoire of sensors (sensory neurons) and effectors (motor neurons), flies flexibly perform complex behaviours such as navigation, aggression, and courtship^23,43–47^. We comprehensively mapped sensory and motor neurons across the brain and nerve cord (Fig 2a and Fig S2a). Our hierarchical annotations include specialised information such as the receptor subtype for sensory neurons and the exit nerve and muscle innervation of motor neurons (Fig 2b). Combining these annotations with the whole-CNS connectome, we can investigate full sensory-to-motor circuits.

**Figure 2.**
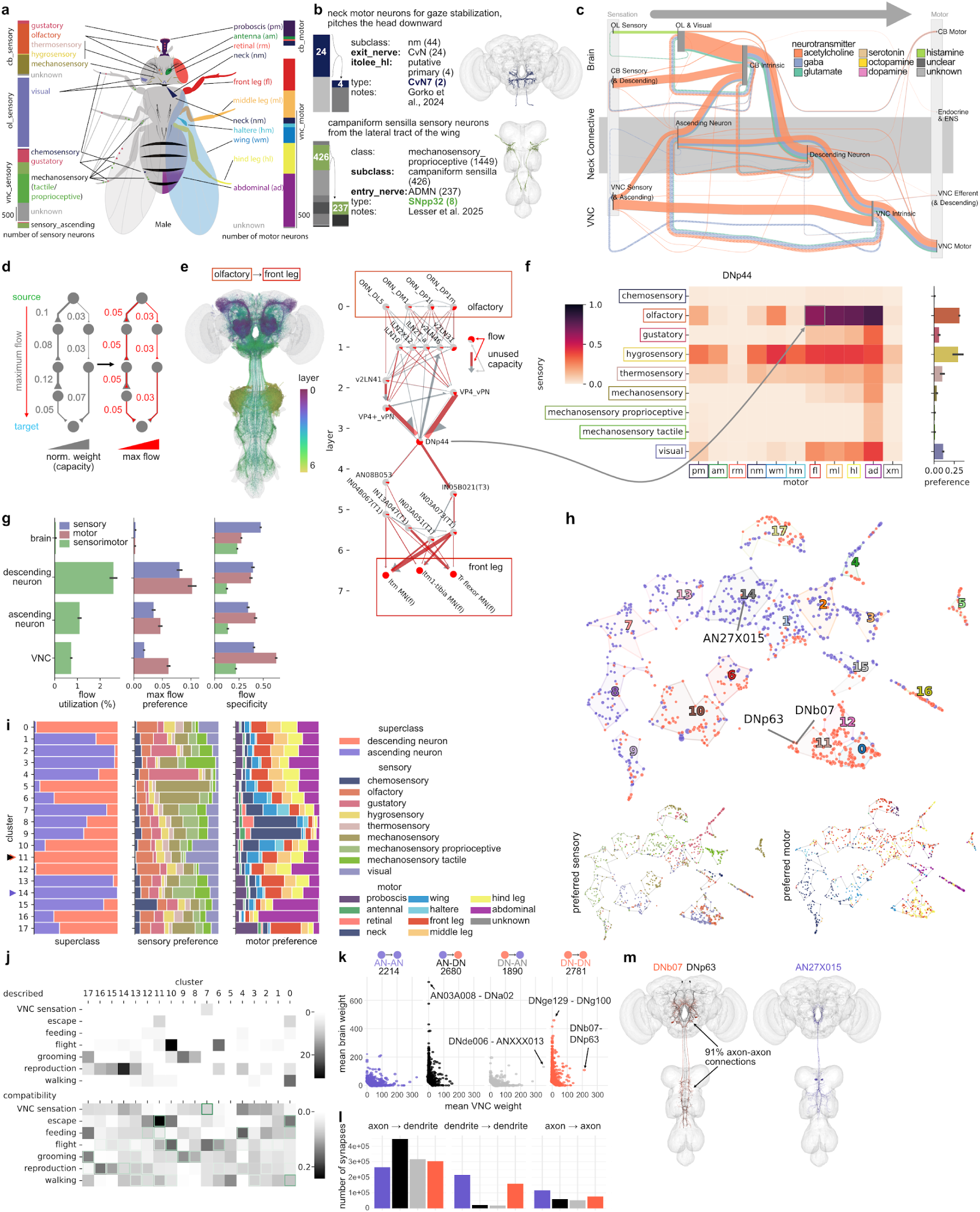
Information flow from sensory input to motor output organizes circuits spanning brain and nerve cord. **a** Schematic of sensory groups (by `class`) and motor group (by `subclass`). Bars show number of neurons per group, lines the peripheral origin or target. Asterisk mark examples in (b). **b** Sample detailed annotation for sensory and motor neurons. Top: morphology of neck motor neuron CvN7. Stacked bar for exit nerve and hemilineage (`itoleeHl`). Bottom: wing campaniform sensilla sensory type SNpp32 in the VNC. Stacked bar for subclass and entry nerve. Grey shades indicate annotation values besides those of the sample neurons. **c** Directed flow diagram of connections from sensory to motor (and endocrine) through the entire CNS. Colours indicate neurotransmitter identity, dashed lines show feedback connections. Edge width is proportional to the number of synapses between groups, node width is proportional to synaptic connections within each group. Edges with fewer than 20k synapses omitted. **d** Schematic of maxflow analysis. Left: normalized edge weights establish potential information flow capacities between nodes. Right: maxflow assignment of flow value across each edge: flow is constrained by the minimum capacity along a path. **e** Example of maxflow routes from olfactory sensory inputs to front leg motor outputs. Left: neurons (flow value > 0.25) colored according to their pseudo-layer ordering. Right: strong flow partners (by type) upstream and downstream of example neurons in type DNp44. Arrow to (f) indicates where the flow for this sensory-motor pairing fills in DNp44’s overall sensory-to-motor flow matrix. **f** Example of mean maxflow through neurons in type DNp44 (left) and sensory modality preference averaged from flow (right). **g** Flow for neck connective neurons. Proportion of flow utilization of different superclasses (left), max flow preference (middle), and flow specificity for one sensory group, motor group or a single sensory to motor group (right). **h** UMAP embedding of DN and AN types by sensorimotor flow, coloured by superclass and sized by presynaptic sites, outlined by numbered clusters (top). Types shown in (m) are annotated. Bottom left: coloured by preferred sensory modality, sized by maximum sensory preference value. Bottom right: as above but for motor preference. **i** Composition of superclass, preferred sensory preference and motor preference for clusters in (h). Arrows indicate clusters containing types in (m). **j** Behavioral categorization of clusters. Top: histogram of cluster membership of neck connective types described in literature by behavior category. Bottom: compatibility of clusters with behavior categories as predicted based on maxflow analysis. Frequency of descriptions from top panel are overlaid in green. Receiver operating characteristic area under curve 0.84 when compatibility is treated as a multilabel classifier of described behavioral category. **k** Connection pairs of neck connective types by superclass with mean weight above 20. Plotted by strength of connection in the brain and VNC. Arrows highlight examples of strong connections. Numbers above show type-type connections for each motif. **l** Axon-dendrite split of DNs and ANs shows the type of connections made by pairs between these superclasses. Only the three most prominent types of connections are shown: axon to dendrite, dendrite to dendrite, axon to axon. **m** Example of a strong DN to DN connection that is 91% axo-axonic. DNb07 and DNp63 have similar axonal arbours in the VNC, but form distinct dendritic arbours in the brain. The glutamatergic DNb07 synapses onto the cholinergic DNp63 in both the brain and VNC, an inhibitory motif. 30% of their downstream partners are shared, yet the only common target in the brain is AN27X015. AN27X015 projects back from the VNC axonal domain of DNb07 and DNp63 to their brain dendrites, but is preferential for mechanosensory (tactile & proprioceptive) modalities and proboscis and abdominal motor domains.

Fig 2c gives a bird’s eye view of the connectome by considering only connections between superclasses. Information primarily traverses the CNS in a feedforward manner: sensory information from the head and body enters via brain and VNC nerves, respectively, and exits mostly via VNC motor neurons. Across levels, feedforward synaptic connections have similar, predominantly excitatory, neurotransmitter compositions, while feedback connections have a larger inhibitory proportion. The neck connective is a key bottleneck: descending neurons constrain information traveling from brain to VNC, and ascending neurons constrain flow in the opposite direction. However, neck connective neurons are integrators not just simple relays spatially bridging the two main divisions of the CNS. For example, ascending neurons output a similar number of synapses as descending neurons onto VNC-intrinsic neurons while also receiving more information from VNC-intrinsic neurons than from direct sensory inputs. Neck connective neurons also participate in the majority of feedback connections in this sensory-to-motor orientation. In the rest of this section, we examine the organization of sensory-to-motor information flow in the neck connective.

Unlike prior datasets spanning only the brain or VNC, an intact whole-CNS connectome allows all neurons and circuits to be contextualized with respect to their role in information flow from sensors to effectors. To investigate, we employed maximum flow analysis, a commonly used network analysis technique^48^. In brief, this finds the greatest possible rate at which information can move through the network from a source to a destination without exceeding capacity limits of the connections (here set by the synaptic weights between neurons). By systematically redistributing flow across alternative routes, the analysis identifies bottlenecks and the overall throughput limit of the network. Compared with the alternatives, maximum flow is better able to reveal long range information flow and can yield distinct measures for each source and sink pairing; however, like most methods it does not leverage the sign of connections, and because flow can be maximal without actually using every possible path, negative inferences should be made with caution.

Specifically, we used maximum flow analysis originating from all sensory neurons of a particular modality `class` (as a group), and terminating at motor neurons from each motor domain `subclass` (Fig 2a,d). This analysis was performed for all pairs of sensory modalities and motor domains on a presynaptic-input-normalized connectome graph, recording the resulting flow over all neuron-to-neuron edges and neuronal nodes. For example, DNp44 descending neurons have high maximum flow to many motor domains from olfactory and hygrosensory neurons, indicating they may prefer information flow from these modalities (Fig 2f). In the strongest flow routes from olfaction to the front leg motor domain (Fig 2e), DNp44’s strongest upstream flow contributors are VP4 thermohygrosensory projection neurons known to be dry air sensing^49^. Upstream of these, the strongest flow originates from olfactory receptor neurons including several associated with aversive odorants (DP1l/m, DL5), passing through interneurons including GABAergic inhibition. Thus DNp44 carries hygrosensory information, which may be gated by olfactory information.

Examining sensory-motor flow over the entire CNS highlights the distinct role of neck connective neurons in information processing. First, ascending and descending neurons have a much higher flow utilization (i.e. the mean percentage of their synaptic capacity utilized across all sensory-motor flows) than brain- and VNC-intrinsic neurons, indicative of their role as information bottlenecks (Fig 2g, left). Second, these flows are more specific towards particular motor domains (e.g. front or hind legs) than they are towards particular sensory modalities (e.g. gustation or thermosensation). This trend consolidates in the VNC, where circuits are more segregated as they fan out to control particular effectors (Fig 2g, middle). Third, while intrinsic neurons of both the brain and VNC are on average more specific to particular sensorimotor pairings, ascending and descending neurons tend to be specific for a particular sensory modality or a particular motor domain, but not both (Fig 2g, right, Fig S2b). In other words, neck connective neurons are multimodal (either in sensory or motor space); we propose that this lower specificity relative to other CNS neurons is driven by the joint constraints of the information bottleneck and the complex behaviours these neurons must orchestrate.

To better understand this multimodal structure within the neck connective, we clustered all descending and ascending neurons on the basis of an embedding of their sensorimotor flow (Fig 2h). While some clusters are predominantly composed of only ascending or descending neurons, or have high specificity to a single sensory modality or motor domain, many are more heterogeneous (Fig 2i). Neck connective cell types whose function has been described in previous literature were sorted into broad behavioral categories, which often correlated with the clusters (Fig 2j, top). We next created simple rules defining which sensory-motor pairings are compatible with which behavioral categories (Fig 2j, bottom, see <u>Methods</u>); this reliably captured known behavioural categories and should therefore help generate new functional hypotheses. Clusters also structure connectivity between ascending and descending neurons (Fig S2c), which could previously only be analyzed by matching of some cell types across partial connectome datasets from different animals^50^.

For a more granular understanding of how connectivity between neck connective neurons relates to information flow and connectivity with other populations, we analyzed type to type connections within and between ascending neurons (AN) and descending neurons (DN) (Fig 2k). There are few strongly connected types for each of the four superclass pairings. DN-DN and AN-DN synapses are predominantly in the brain, while AN-AN and DN-AN synapses are predominantly in the VNC. For AN-DN and DN-AN pairings, this is consistent with an expectation of axo-dendritic connectivity and archetypical localisation of arbours from these superclasses. While we observe that connectivity is predominantly axo-dendritic in all four cases, a larger proportion of connectivity within DNs (27%) and ANs (34%) is dendro-dendritic than for the other two cases (both 4%) (Fig 2l).

Our flow analysis allows discovery of new functional relationships, such as a motif involving 3 cell types that have yet to be experimentally investigated (Fig 2m). DNb07 and DNp63 only have one common strong upstream partner but both belong to cluster 11 since they share a flow of olfactory and visual information towards the same motor effectors (Fig S2d). Intriguingly, DNb07 makes inhibitory connections onto the axons of DNp63 suggesting that they form a functionally antagonistic pair, likely driven by different features of the same sensory modalities. The shared upstream partner is actually an inhibitory ascending neuron (glutamatergic AN27X015) with quite different tuning (tactile & proprioceptive modalities, cluster 14) suggesting that it could be a feedback signal relating to the motor state of the animal. Molecular reagents for these DNs are already available^51^ which will permit functional exploration of these and many other hypotheses.

### Identifying sexually dimorphic circuit elements

Sex differences in behaviour likely depend on the interaction of unique circuit elements, sex-specific circuit wiring and sex-dependent physiology of cells within^26,52–54^ and outside the nervous system^55^. Connectomics enables comparison of all structural differences in the nervous system at synaptic resolution. By comparing this male dataset with female brain connectomes (FAFB/FlyWire^30,31^ and hemibrain^32^) we now quantify the prevalence of sex-specifc neuronal cell types versus shared circuit elements that make different connections.

Cell types (most of which consist of multiple neurons in the adult fly), represent the fundamental unit of conservation across brains and brain hemispheres^31,56,57^. We assigned a type match for 96.4% of neurons in the male CNS brain and 98.8% of the optic lobe with one or both female brain datasets (Fig 1h, Fig S1e). This includes both *isomorphic* and *dimorphic* neurons, while unmatched neurons are candidate *sex-specific* types. We found that it was crucial to have specific operational definitions for these categories (detailed in Table 1 and schematised in Fig 3a). These definitions focus on neurons that exhibit clear morphological differences between the sexes. We will investigate “secondarily dimorphic” neurons that are morphologically isomorphic but still exhibit dimorphic connectivity later in the <u>results</u>. Due to the absence of a fully-proofread female VNC connectome, we extended dimorphism labels from previous literature describing the VNC^27,58–63^, especially recent work of Stürner *et al.*^50^ (see <u>Methods</u> for details).

**Figure 3.**
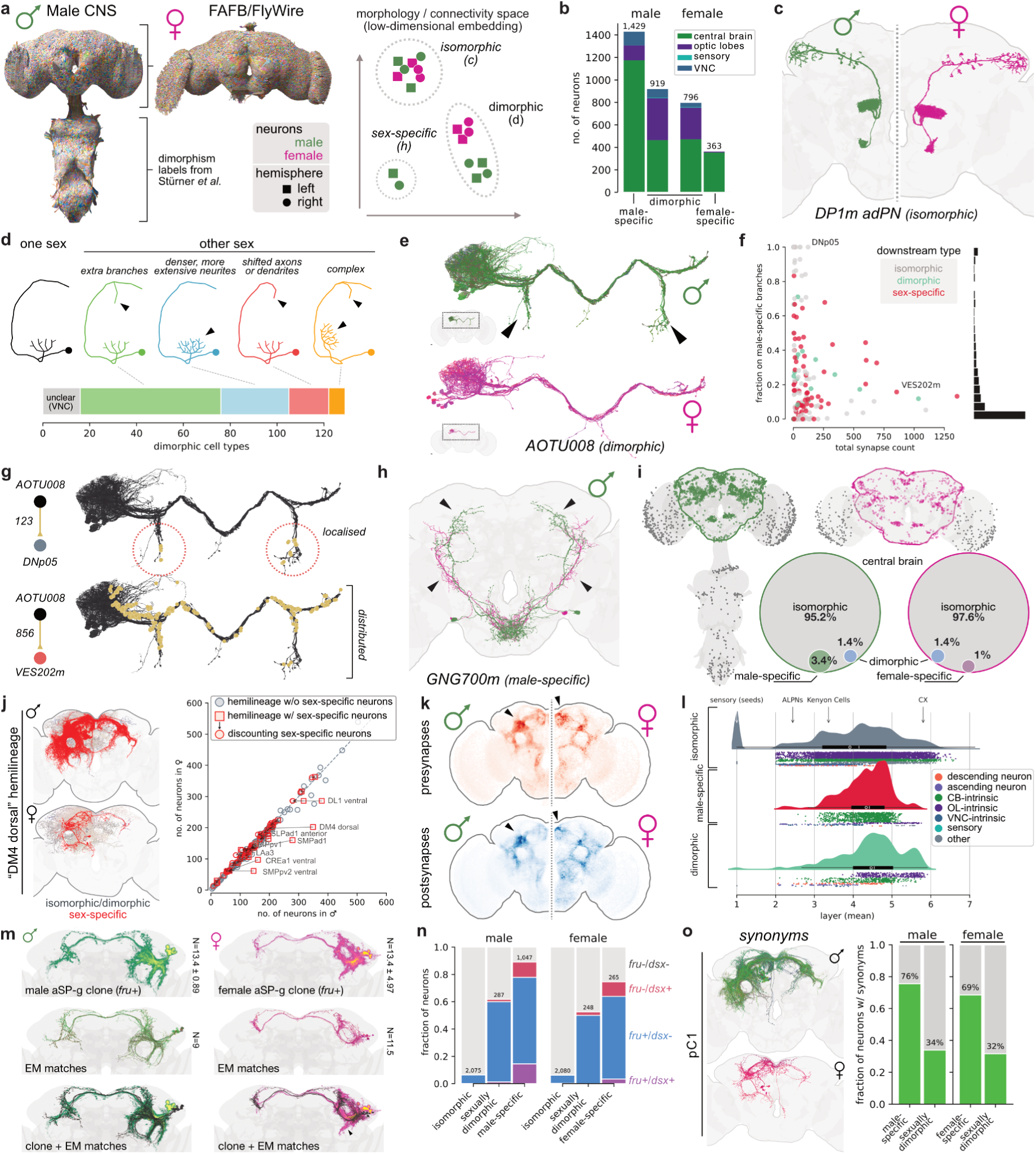
Sexual dimorphism in the fly brain. **a** Schematic illustrating differences between iso-dimorphic and sex-specific neurons in morphology or connectivity space. See also Table 1. **b** Number of dimorphic & sex-specific neurons broken down by superclass. **c** Example for isomorphic cell type. **d** Classification of dimorphic types based on morphological differences in the brain. **e** Example for dimorphic cell type with extra branches. Arrowheads highlight ventro-lateral axons only present in male neurons of this type. **f** Fraction of synapses on male-specific branches versus total synapse count for each outgoing connection made by AOTU008. **g** Examples for distributed connections and connections highly localised to male-specific branches. **h** Example for sex-specific cell type: GNG700m (green). The closest isomorphic cell type (AVLP613, magenta) is shown for comparison. Arrowheads point out differences in axonal projections. **i** Spatial distribution of dimorphic and sex-specific neurons’ somata. Central brain somas are coloured in green and magenta for male and female, respectively. Pie charts show the proportion of central brain neurons that are dimorphic/sex-specific. **j** Left: example hemilineage in male and female with sex-specific neurons highlighted in red. Right: neuron count per hemilineage in male versus female. Discounting sex-specific neurons tends to align numbers. The labelled hemilineages collectively produce >50% of all dimorphic & sex-specific neurons in the male. **k** Distribution of pre- (outputs) and postsynapses (inputs) of dimorphic & sex-specific neurons in male and female connectome. Arrowheads highlight differences between sexes. **l** Layer assignments relative to sensory inputs for all neurons in the male CNS split by dimorphism. Data points are coloured by superclass. Upper half shows the kernel density estimate of the underlying distribution. Mean layers for antennal lobe projection neurons (ALPNs), Kenyon Cells and central complex neurons are shown as landmarks. Boxplots shows median (vertical line), mean (circle), 1st-3rd quantile (box) and 1.5IQR). **m** Light-microscopy (LM) image of a *fruitless* (*fru+*) clone (top row) and matching neurons found in the EM (middle row)^63^. Arrowhead points at variable branches in female aSP-g clones. **n** Fraction of neurons in the central brain labeled as *fruitless*- and *doublesex*-expressing (*dsx+*) based on LM-EM matches. **o** Left: pC1 (includes P1 neurons) as an example for a synonym assigned to a group of dimorphic neurons. Right: fraction of dimorphic neurons matched to prior literature.

**Table 1:**
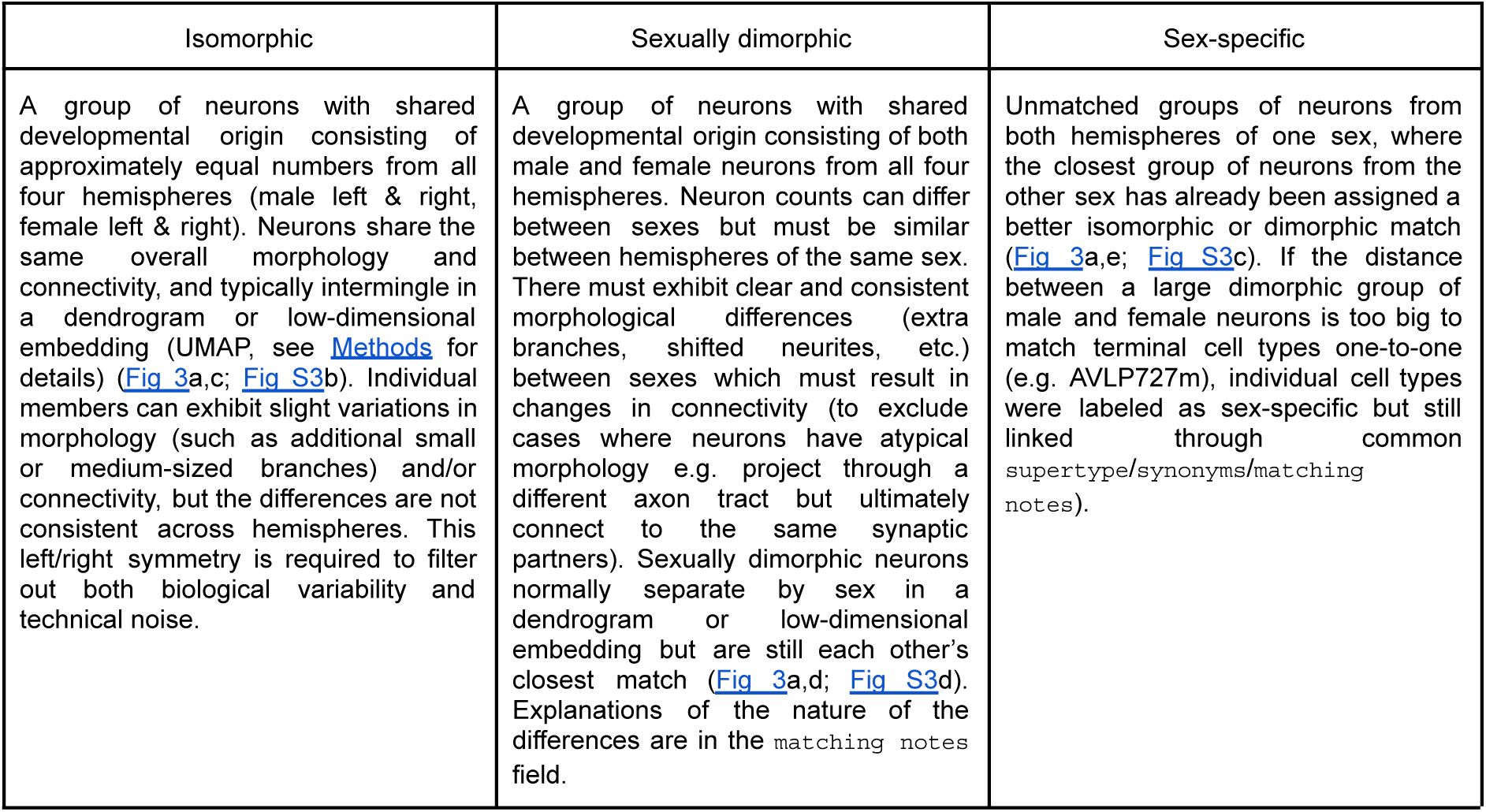
Definitions of dimorphism.

Applying these criteria, we identified 1,427 sex-specific neurons across the male CNS versus 363 female-specific (Fig 3b). We found 924 dimorphic neurons in the male CNS matching 811 neurons in the female; we would have expected the same number in both sexes and a large part appears to be biological variability (see Fig S3f), especially in the numbers of columns in the optic lobes^36^ (male CNS ∼900; FAFB ∼800). A relatively modest 0.1% of neurons in the male visual system are sex-specific and 0.3% are sexually dimorphic (Fig S3f). In the male central brain, dimorphism is much more prevalent: 3.4% of neurons are male-specific and 1.4% are sexually dimorphic (Fig 3i). By contrast, female-specific neurons make up only 1% of the central brain.

Where do sex-specific neurons originate? Female-specific neuronal apoptosis during pupal development is a key mechanism^64,65^ but it is not evenly distributed. Of the ∼200 hemilineages forming the central brain^66,67^, about a quarter produce sex-specific or sexually dimorphic neurons but just 8 of these (4%) produce more than half of all sex-specific and dimorphic neurons (Fig 3j, Fig S3g). For example, the DM4 dorsal lineage has 349 neurons in males versus 205 in females; but if we exclude sex-specific neurons, counts are – as expected – almost identical across sexes (199 versus 201). Approximately 8% of sexually dimorphic and sex-specific neurons are primary (i.e. early born neurons that function in the larva, Fig S3g).

#### Topology of sexual dimorphism

Is there a logic to the morphological differences in dimorphic neurons that underlie much of dimorphic connectivity? We observed three categories: completely new arbours, shifts, and expansions of existing branching patterns; these features were sometimes combined (Fig 3d). AOTU008 (part of aSP-i^63^) is a clear example for new arbours: it extends ventral axonal projections in males that are completely absent from female homologues (Fig 3e)^27^. As expected, this cell type has many dimorphic and sex-specific connection partners (5.5% of inputs and 57% of outputs). Surprisingly, the abundant dimorphic connections are not exclusively localized to the male-specific arbours (Fig 3f). For example, AOTU008 makes all of its connections onto DNp05 from its male-specific branches, but outputs to VES202m are distributed across the entire axon (Fig 3g). We can rationalise this as follows: AOTU008 can only connect to the isomorphic dendrites of DNp05 by extending an extra axon. But VES202m is a male-specific neuron directly adjacent to AOTU008, so connections can involve the entire neuron. We will further explore this anecdotal observation in later sections.

Where do sex-specific and dimorphic neurons make connections? We find their synaptic distribution is spatially highly non-uniform with a strong preference for integrative, higher brain regions of the protocerebrum (Fig 3k) consistent with earlier observations on the location of *fruitless*-expressing neurons^63^. We also quantify this by neuropil region, observing weaker but still significant enrichment in the lobula and gnathal ganglion which are closer to the sensory periphery (Fig S3h). The spatial distribution is very similar across sexes, but there are exceptions for example in the SMP and SIP, which are enriched in the male and female, respectively (Fig S3h,i).

These results suggest that dimorphic elements are concentrated in integrative neurons that we expect to be at intermediate locations in sensory-motor pathways in the connectome. We tested this intuition by applying a probabilistic graph traversal model, which starts from sensory neurons and assigns a layer to every neuron in the connectome graph^68^ (Fig 3l). Isomorphic neurons are distributed relatively evenly across 7 layers with a mean at layer 3.9. In contrast, sex-specific and dimorphic neurons tend to occupy deeper layers, with a mean at layer 4.3 and 4.5, respectively. Conversely the sensory periphery (corresponding to layers 1-3) contains many fewer such neurons: just 2.5% of male-specific versus 18.5% of isomorphic neurons.

#### Genetic basis for dimorphism

In *Drosophila* alternative splicing of two master regulator genes, *fruitless* (*fru*) and *doublesex* (*dsx*) shapes neuronal development to produce sex differences in morphology and connectivity^19,69,70^. Whole-brain connectomes now allow us to co-examine these gene expression patterns alongside wiring differences. The first step was molecular annotation of the connectome, achieved by co-registering CNS and FAFB/FlyWire connectomes with light microscopy datasets of *fruitless*^63,71^ or *doublesex*^27^ expression (Fig 3m); neurons were marked as low confidence where the evidence was less conclusive and counts were validated against previous studies^27,63,64^ (see <u>Methods</u>). Our analysis focuses on the central brain as technical limitations prevented exhaustive annotation of the VNC. We annotated 4,505 (2,695 high confidence) *fruitless*- and 407 (332 high confidence) *doublesex*-expressing neurons in the male CNS (2.7% and 0.2% of neurons, Fig S3j). 251 cells co-expressed both transcription factors. Most (73%) *fruitless*/*doublesex* annotations are in the central brain (Fig 3l) where 9.5% of male neurons are annotated as *fru+*; this represents an upper boundary compared with a lower boundary of 4.7% when using only high-confidence annotations.

Past studies almost exclusively examined *fruitless-* and *doublesex-* expressing neurons for sexual dimorphism but connectomics allows us to assess every neuron in the brain. How do these approaches compare? 89% of male-specific neurons in the central brain were annotated as *fru*+/*dsx*+, compared with 61% of dimorphic neurons and just 6.4% of isomorphic neurons (Fig 3n). Conversely, 11% of sex-specific and 39% of dimorphic neurons are *fru-*/*dsx-*. Some of this discrepancy likely results from incomplete gene expression data (see <u>Methods</u>); however, other genetic factors or non-cell autonomous mechanisms could still shape some of these dimorphisms.

There is a rich literature in *Drosophila* examining dimorphic cell types, connectivity, genes, circuits, and behaviours^10,19,20,22,24,26,27^. We provide `synonyms` to cross-reference cell types such as: aSP-g, Kohl *et al 2013*^26^. In total, we recorded synonyms for over 800 cell types corresponding to three quarters of sex-specific and one third of dimorphic types (Fig 3o). We propose many novel dimorphic/sex-specific cell types as 188 (37%) could not be matched to the literature.

### Visual system

*Drosophila* use their visual system to detect light cues for behaviors like navigation and circadian regulation, as well as specific object features such as shape, movement and colour. The visual system is housed primarily in the optic lobes, which consist of five neuropils (the lamina, medulla, lobula, lobula plate, and accessory medulla) and contain four neuronal superclasses (sensory, optic lobe intrinsic, visual projection, and visual centrifugal neurons) (Fig 4a). A full catalogue of the right optic lobe neurons in this male CNS connectome was presented in Nern *et al*.^36^. Here we expand our analyses to the left optic lobe. We observe excellent agreement between cell types, cell counts and connectivity between the two optic lobes (Fig S4Ba-e). We also improve our annotations of pale and yellow column types in the medulla of both optic lobes (Fig S4Bf-j and see <u>Methods</u>); see also the updated Cell Type Explorer web companion for comprehensive analyses of the optic lobe.

**Figure 4.**
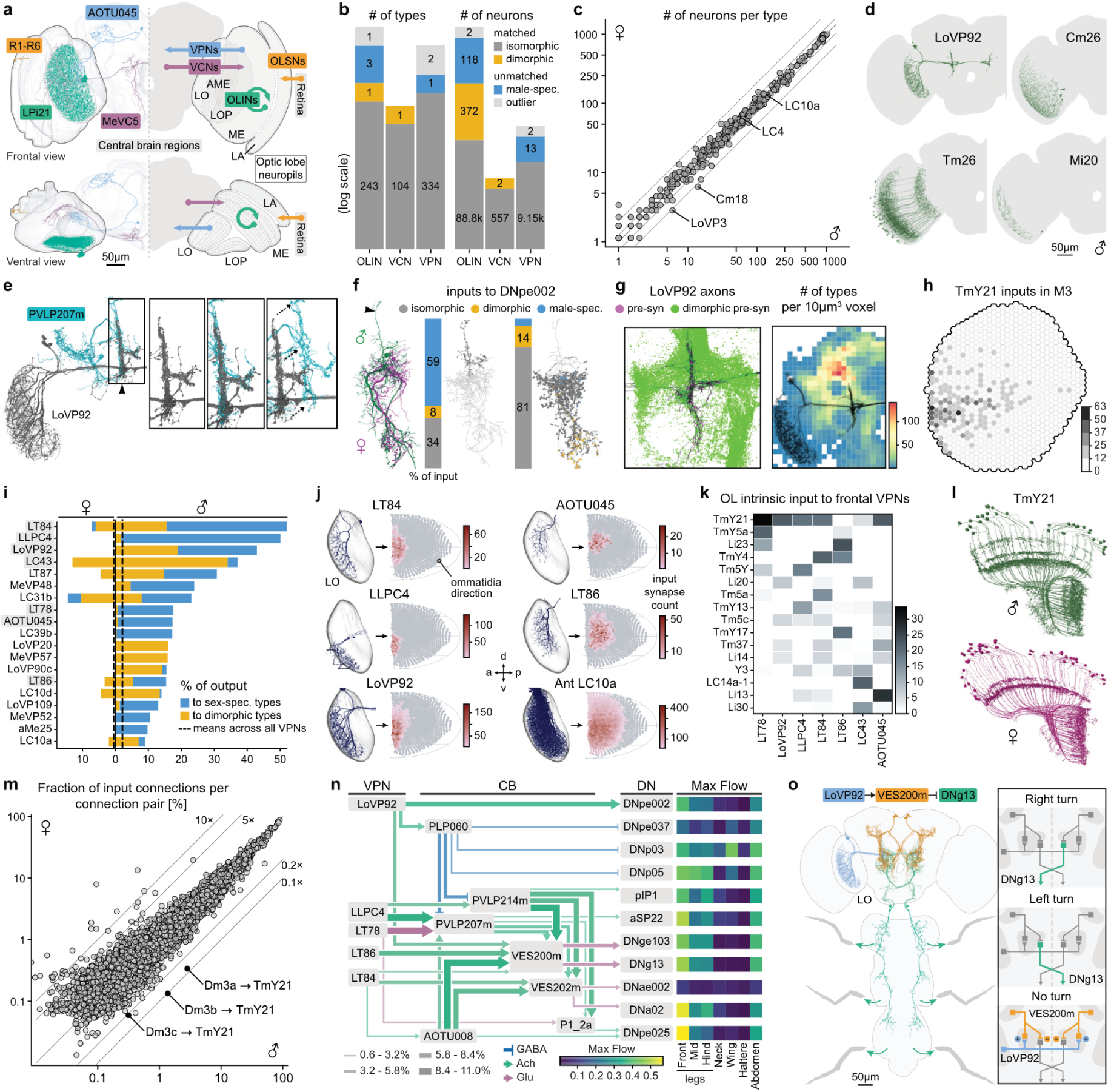
Sexual dimorphism in the visual system reveals a functional “love-spot”. **a** Diagram of the optic lobe showing the different neuropils and neuronal superclasses. LA: lamina; ME: medulla; LO: lobula; LP: lobula plate; AME: accessory medulla; OLSNs: optic lobe sensory neurons; OLINs: optic lobe intrinsic neurons; VPNs: visual projection neurons; VCNs: visual centrifugal neurons. **b** Number of types (left) and neurons (right) per superclass across several categories. A type was classified as male-specific only if it was symmetrical; otherwise, it was annotated as unmatched outlier. **c** Neuron count per type between male and female. Female counts were normalised (x1.134) to account for the ∼100 additional ommatidia in the male eye. Only types matched one-to-one between male and female are included. Photoreceptors and Lai are excluded, as many are missing in the male CNS. **d** The four male-specific visual types. Scale bar, 100 µm. **e-f** Examples male-specific (e) and dimorphic (f) types that converge onto LoVP92 axons. Inset: close-up and shifted view of the boxed area in e. Arrowheads indicate the location of LoVP92 axons. Stacked bar charts show the input-normalized proportion of dimorphism status of connection partners onto DNpe002. **g** Dimorphic synapse distribution around LoVP92 axons (black). Green dots show all synapses between dimorphic or sex-specific types. LoVP92 output synapses are shown in magenta. Right: heatmap showing the number of cell types per 10 µm³ voxel in the right hemisphere of the central brain. **h** Spatial map showing Dm3a,b,c inputs to TmY21 in the third layer of the ME. **i** Output-normalized percentage of VPN outputs to dimorphic or sex-specific types in the female (left) and the male (right). Dotted lines indicate the average percentage output from all VPNs onto sex-specific or dimorphic partners. Types highlighted in grey are frontally biased, as shown in panel j. **j** Examples of frontally-biased VPNs. Spatial coverage heatmaps show input synapse distributions mapped onto a Mollweide projection of the right compound eye’s visual field. Color scale bars show input synapse count. **k** Heatmap showing the strongest inputs from optic lobe intrinsic neurons to frontally-biased VPNs. **l** The dimorphic type TmY21 in the male (top) and female (bottom). Arrowheads indicate differences in arborisation between the sexes. **m** Fraction of input connections for each pair of connections between optic lobe intrinsic neurons in the male and female. Only edges with more than 500 synapses are shown. Lines indicate changes in relative weight by factors of 5 and 10. **n** Network diagram showing male-specific connections between frontal VPNs and DNs. Only DNs reached within two hops were included, using a 2% input threshold – except for DNa02 and DNae002, where a 0.5% threshold was used. AOTU008 did not meet the DN threshold but was included because its top two input partners are VES200m and VES202m. **o** Diagram of a frontal VPN to DN circuit in the fly brain and nerve cord (left), alongside a simplified circuit diagram (right). VES200m links visual detection – via LoVP92 inputs from the lobula – to the steering descending neurons DNg13. Unilateral activation or inhibition of DNg13 has been shown to cause ipsilateral, or contralateral, turning, respectively. LoVP92 activates VES200m bilaterally, and VES200m inhibits DNg13 ipsilaterally. As a result, activation of LoVP92 leads to inhibition of both DNg13 neurons, preventing turning.

The inclusion of the left optic lobe enables the bilateral visual networks of the male to be compared to those in the female. Male fruit flies rely heavily on visual information to direct courtship displays: chasing, ipsilateral wing preference, and overall courtship success are associated with the effective transmission of visual signals^22,72–74^. To investigate sexual dimorphism in the visual system, we compared the anatomy of cells across male^36^ and female^75^ optic lobes finding a high concordance in cell types and neuron counts (Fig 4b,c)^36,76^.

Of 249 cell types intrinsic to the optic lobe, we identified three (Cm26, Tm26 and Mi20) that are sex-specific (Fig 4b,d), and one (TmY21) that is dimorphic (Fig 4b,l). Strikingly, 99.9% of all visual projection neurons (VPN) are isomorphic, and we identified just one male-specific cell type: LoVP92 (Fig 4b,d). Three cell types remained unmatched due to bilateral inconsistency (Fig 4b, Fig S4Aa). We found no sensory or visual centrifugal neurons to be sex-specific/dimorphic (Fig 4b).

LoVP92 consists of about 6 cells per hemisphere. 18% of their input connections and 43% of their outputs are with male-specific or dimorphic partners (Fig 4i, Fig S4Ad). Furthermore, the distinctive LoVP92 axonal arbours co-converge with multiple male-specific and dimorphic types. This includes central brain interneurons (PVLP214m, PVLP207m, VES200 and VES202m, Fig 4e, Fig S4Ab), which are strong downstream partners of multiple isomorphic VPN types (Fig 4n). This architecture effectively converts these VPNs into functionally dimorphic types. We also see focal dendritic innervation of the same region defined by LoVP92 axonal arbours. Four descending neurons have large isomorphic dendrites innervating distinct territories but each sends a small dimorphic dendritic arbour to this region (Fig 4f, Fig S4Ac). Each dimorphic branch is itself heavily enriched in dimorphic and sex-specific partners (Fig 4f, Fig S4Ac). Therefore, the node defined by LoVP92 axons, the only male-specific VPN, may serve as a hotspot, where multiple dimorphic cell types converge to participate in male-specific circuits (Fig 4g, left). We anticipate numerous hotspots in other central brain regions characterised by a high density and diversity of dimorphic connections (Fig 3k, Fig 4g, right).

The gross anatomy of the peripheral visual system is largely isomorphic but visual information flows through VPNs to highly dimorphic circuits in the central brain. Nineteen out of 337 VPN types direct over 9% of their output to sex-specific/dimorphic downstream neurons in the male (Fig 4i). For example, the isomorphic LC10a neurons, critical for accurate tracking during courtship^32,74,77,78^, target sexually dimorphic AOTU008 neurons in both sexes^27^; but AOTU008 makes male-specific ventral axonal projections (Fig 3e-g) and commits 57% of its output to sex-specific and/or dimorphic downstream partners.

We next investigated the visual space sampled by the VPNs with strong connections to dimorphic partners. Seven of the 19 types (Fig 4i) have distinctive, frontally-biased visual fields (Fig 4j, Fig S4Ae,g)^79^. Most VPNs cover the entire lobula, but LoVP92 and 6 isomorphic types are confined to the anterior lobula which samples the frontal visual field of the compound eye. The same frontal bias is also observed in upstream visual processing. These VPNs have strong input from dimorphic TmY21 interneurons (Fig 4k). Inputs to TmY21 are frontally biased (Fig S4Af). This includes inputs from Dm3a-c edge-detecting neurons^36,80^ onto TmY21’s male-specific branches (Fig 4l,h). (Fig 4m). This organisation – and its likely behavioral significance – is reinforced by observations on LC10a VPNs. LC10a covers the whole visual field but recent work has shown that a subgroup targeting the same frontal region is required for precise steering toward the midline^78^.

We therefore propose that frontally biased VPNs connect onto strongly dimorphic central brain circuits rerouting the flow of visual information to support sex-specific social behaviours. This likely includes courtship, where males pursue females from behind, maintaining them within the frontal field of view^29,74,81,82^. This specialised processing of a distinct region of visual space is reminiscent of at least two long-appreciated features of visual organisation, the fovea and the insect “love spot”. Larger flies have a love spot^83^, where male eyes feature specialised photoreceptors^84^ that are proposed to be tuned for visual target tracking during courtship^85–87^. While *Drosophila* males lack the prominent external eye specialisations present in larger flies, the frontally-biased VPNs may serve an analogous function to male-specific/dimorphic neurons in other insects that sample the visual space imaged by the love spot^88,89^. Secondly, recent observations in both sexes show that the *Drosophila* visual system has features of the foveal specialisations found across species, including mammals: higher acuity eye regions^79^ and their over-representation in optic lobe circuits^36^.

To link these neurons to dimorphic motor behaviours, we investigated pathways downstream of the frontally-biased VPNs and identified 11 descending neurons, including DNa02 and aSP22, required for effective male steering during pursuit^78^ and later steps of courtship^90^ (Fig 4n, Fig S4Ah,i). Besides these pathways with known courtship functions, we found additional connections that could support male steering towards flies of either sex. For example, LoVP92 connects through inhibitory male-specific interneuron VES200m to descending steering neuron DNg13 (Fig 4o). DNg13 is a general purpose coarse steering pathway^91^. We therefore hypothesise that when the female is directly in front of the male, LoVP92 inhibits DNg13, favouring fine tracking near the midline through other descending pathways.

In summary we provide circuit-level evidence for a behaviourally relevant “love spot” in *Drosophila melanogaster*: male flies detect fly-sized objects in their central visual field ^74^, and this information traverses sexually dimorphic neural circuits through to descending pathways likely facilitating successful courtship pursuit.

### Auditory system

Auditory cues are central to *Drosophila* courtship. Male flies vibrate their wings to produce species-specific songs, enabling females to assess species identity and mate quality^92–95^. *D. melanogaster* males generate two main song types, pulse and sine song, controlled through nested networks in the wing neuropil of the nerve cord^58,96^. Songs elicit sexually dimorphic responses: receptive females slow down and open their vaginal plate for copulation^61,94,95,97–99^; males become aroused with increased locomotion, persistent courtship displays, and intermale competition^100–104^.

Song detection begins in the antennal Johnston’s organ (JO). Mechanosensory neurons have discrete frequency-tuning and project a tonotopic map to the AMMC, the primary auditory processing centre in the brain^105,106^. Low frequency courtship song is detected by JO-B sensory neurons^102,107^. Despite segmentation issues (Fig S5a), we found no evidence that these sensory neurons are themselves sexually dimorphic^100,102,103,108,109^. JO-B targets include aPN1, a *fru+* but isomorphic population^100,102^ (Fig 5a) essential for detecting courtship song, especially pulse song^102,109,110^. To enable comparison of central auditory processing across sexes, we annotated 8 further song-responsive cell types described in the literature^100,102,103,108,110–112^.

**Figure 5.**
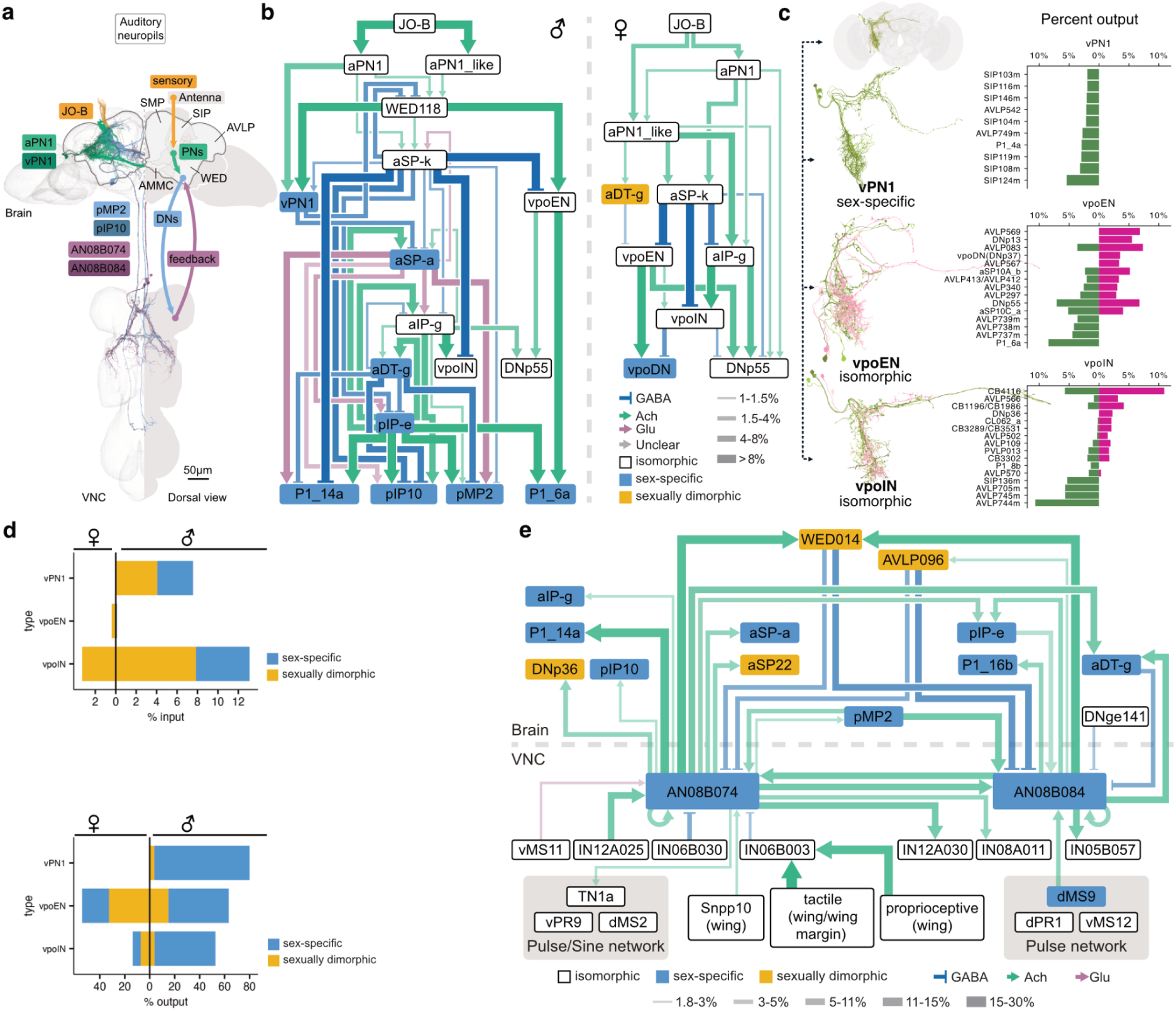
A recurrent feedback loop controls courtship song detection and action in the auditory system. **a** Overview of the auditory pathway and recurrent feedback circuit. Song induced vibrations move the arista to activate Johnston’s organ B neurons, activating a sensorimotor loop that provides feedback from song generating networks back to the central brain. **b** Circuit schematics for male (left) and female (right) from JO-B to higher order processing centres. Boxes denote cell types, edge thickness scales with synaptic weight, and edge colors indicate transmitter identity. Node colors mark sex classes: isomorphic (white), sexually dimorphic (gold), sex specific (blue). Line widths denote input-normalized connection strengths. **c** Top 10 output partners of vPN1 (top), vpoEN (middle) and vpoIN (bottom) in males and females. Bars show the fraction of total synaptic outputs (right) (output-normalized) to each identified type across sexes. **d** Quantification of partner identity for vPN1, vpoEN and vpoIN. Top: fraction of inputs from sex-specific/dimorphic neurons in the female (left) and male (right). Bottom: fraction of outputs to sex-specific/dimorphic neurons in female (left) and male (right). **e** Connectivity between central brain song detection and ventral nerve cord song production circuits. The established pulse and sine song network in the wing neuropil is reproduced and connected to identified descending neurons and to male specific ascending neurons that project back to central brain nodes. Line widths denote input-normalized connection strengths. Scale bars as indicated.

Receptive females allow copulation by opening their vaginal plate through vpoDNs (DNp37), which are modulated by excitatory vpoENs and inhibitory vpoINs, both tuned to conspecific courtship song^53^. Despite this female-specific behavioural role, we identified vpoEN and vpoIN in the male CNS and surprisingly found no clear morphological dimorphism (Fig 5c, Fig S5c). However, their connectivity is necessarily very different: vpoDNs (and the vaginal plate) are absent in males. Approximately half of vpoEN outputs (but not inputs) are onto sex-specific/dimorphic cell types in both males and females. In males, vpoIN also has strong outputs onto sex-specific/dimorphic neurons but female vpoINs primarily target isomorphic types (Fig 5d). Comparing vpoEN downstream partners between males and females, we find strong sex-shared (DNp55) and sex-specific downstream partners (vpoDN in females, P1_6a in males) (Fig 5b,c). This last pair defines a changeover switch: male and female brains reroute the same sensory information (in vpoEN) to sex-specific downstream neurons.

In males, courtship song induces chaining and functional analysis has begun to elucidate the circuit basis of this striking behaviour where multiple males court each other. Chaining depends on aPN1 and male-specific *fru+* vPN1 neurons^100^, which respond to pulse and sine songs and appear to excite male-specific P1 neurons^100^. However, vPN1 is strongly predicted as GABAergic so excitation cannot be direct (Fig S5e). Connectomic analysis likely resolves this by identifying a strong disinhibitory circuit: vPN1 inhibits a male-specific mAL which inhibits P1. This module may switch male behaviour between aggression and courtship.

Downstream, we identified song DNs (pIP10^23^ and pMP2^58^) and key wing pre-motor neurons in the ventral nerve cord (TN1A, vPR9, and dPR1^58^). Extending previous work^50,59^, we also identified a set of male-specific ascending neurons (AN08B074 and AN08B084). These feed information from VNC song generation neurons to *fru+* song detection circuits (aSP-a/pIP-e) in the central brain as well as song DNs. These connections are actually bidirectional: song detection neurons also synapse onto the AN axonal arbours in the brain (Fig 5b,e, Fig S5g). This creates a closed loop in which the motor output of singing likely influences auditory detection of song; indeed it may function as an efference copy allowing males to distinguish their own song from that of others^113^, in turn influencing the male’s arousal state.

### Olfactory system

From flies to mice, volatile pheromones can elicit different behavioural responses in males and females^114,115^. Changes in the morphology and connectivity of pheromone-responsive neurons have been found in higher brain areas such as the lateral horn^26,116^ but not in the early olfactory system^114,117^. Now we can follow pheromone signals into the antennal lobe (AL, analogous to the mammalian olfactory bulb, Fig 6a) and across the CNS connectome to identify likely sex differences in processing.

**Figure 6.**
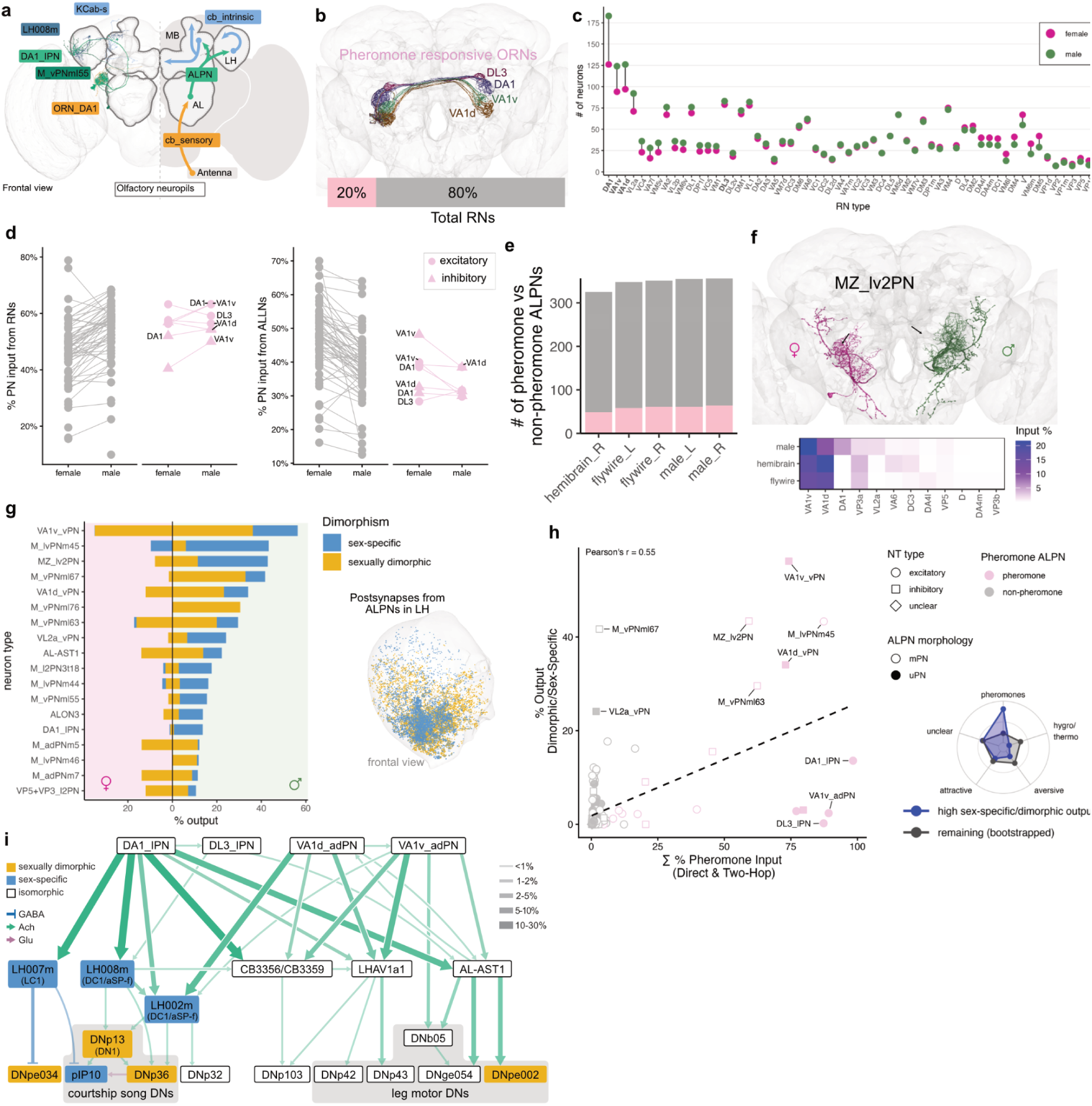
Dimorphism emerges in the third-order of the olfactory system. **a** Illustration of the olfactory system describing flow of information from sensory neurons to the antennal lobe (`cb_sensory`) and from there to the lateral horn and mushroom body (`cb_intrinsic`) via antennal lobe projection neurons (`class: ALPN`). The left hand side shows example neurons, the right hand side shows the diagrammatic flow of information. AL: antennal lobe, LH: lateral horn, MB: mushroom body, PED: peduncle, CA: calyx. **b** Rendering of pheromone responsive ORNs: DA1, DL3, VA1d, VA1v. 10 neurons are shown for each type. Below, the percentage of pheromone RNs relative to all RNs. **c** Comparison of the number of RNs per type in the male (green) and female (FAFB/FlyWire: magenta) brains. The thermo/hygrosensitive glomeruli (VP1-5) are shown on the right of the plot. **d** Comparison of the percentage input to uniglomerular ALPNs in the male and female brains (FAFB/FlyWire) for pheromone (pink, distinguishing excitatory from inhibitory) and non-pheromone (grey) types. **e** Number of pheromone and non-pheromone ALPNs per side in the male and female brains. An ALPN is defined as being either from class ALPN or ALON and receiving at least 5% of its input from pheromone ORNs and their uniglomerular PNs. **f** One of two sexually dimorphic ALPN, MZ_lv2PN, in the male (green) and FAFB/FlyWire (pink). Below, a heatmap of its top RN input (over 0.1% per type) in the male and female brains. **g** Left: ALPNs that provide at least 10% of their output to sex-specific or dimorphic neurons, in male or female brains. The plot shows percentage output in the male and female (FAFB/FlyWire) brains to dimorphic (yellow) and sex-specific (blue) neurons. Right: ALPN postsynapses of cell types shown in the plot, coloured by type of dimorphism. **h** Correlation analysis of the percentage of pheromone input for ALPNs (direct and two-hop) versus the percentage output to dimorphic or sex specific neurons. Pheromone ALPNs in pink. On the right: Bootstrapped comparisons of sex-specific/dimorphic targeting across ALPN groups defined by dominant RN input valence. **i** Network diagram showing strong and selected downstream targets of the uniglomerular pheromone ALPNs (up to 2 hops).

We identified and cross-matched all 53 olfactory and 7 thermo/hygro-glomeruli sensory neuron types, each projecting to a single AL glomerulus (Fig S6a,b, <u>Methods</u>). We began by comparing male and female sensory neuron numbers (Fig 6c). Sorting glomeruli by the difference in male and female counts revealed a male bias in just 4/60 types (DA1, VA1v, VA1d and VL2a). This includes 3/3 *fru*+ types, and 3/4 glomeruli known to detect sex pheromones^114,118,119^ (Fig 6b) while food responsive VL2a is pheromone-like^120^, promoting male courtship. For the remaining ORN types, there was good agreement between the numbers in female and male connectomes.

Classic studies in moths linked male-enlarged glomeruli to processing of specific pheromone blend components detected by receptor neurons that are numerically enriched on the male antenna^121^. Low resolution studies found volume differences^122^ in 3/4 *Drosophila* glomeruli highlighted above but no differences in numbers of receptor neurons^123^. Our examination of n=2 brains precludes statistical analysis, but the targeted numerical differences in four biologically relevant glomeruli strongly suggest a true sex difference. Increases in sensory neuron number may allow males to sustain pheromone responses during courtship despite exposure over tens of minutes, analogous to recent comparative studies across drosophilid species of sensory neurons that detect host plant volatiles^124^. This logic is also similar to our observations of a frontal sensory bias in the visual system (Fig 4).

Given the differences in sensory neuron (ORN) counts for some glomeruli, we looked for changes in connectivity with downstream antennal lobe neurons. We observed a trend towards higher ORN and lower local interneuron (ALLN) input onto AL projection neuron (ALPN) in males but this was global and not specific to particular glomeruli (Fig 6d). Crucially, when comparing ORNs, ALPN and ALLN connectivity between male CNS and both female datasets (FAFB/FlyWire and hemibrain) (Fig S6c) we found that FAFB/FlyWire is the outlier, very likely owing to differences in electron microscopy and image segmentation. This highlights the need for careful analysis of connectivity differences, particularly when interpreting global trends for a class of neurons as opposed to changes between specific cell types of the same general class.

We next annotated the output neurons of the antennal lobe (projection neurons, ALPNs, 188 types); many have dendrites innervating a single olfactory glomerulus, but others, which have historically received less attention despite constituting 60% of the population^68^, integrate information across multiple olfactory channels. A threshold of 5% input from pheromone ORNs defined 136 pheromone ALPNs (27 types, of which 21 are multiglomerular), many of which have not been functionally investigated (Fig 6e). Just 2/188 types are sexually dimorphic; one of them, multiglomerular MZ_lv2PNs, receives pheromone input from DA1 and VL2a ORNs in males but not females as well as non-pheromone olfactory and thermosensory input (VP3a+b) in both sexes (Fig 6f). These two pheromone pathways are known to regulate male courtship behaviour^125^ but the reason to combine all these inputs in a single neuron remains to be explored.

Consistent with our observations in the visual and auditory system, dimorphism of second order neurons is thus extremely limited. Instead key transformations occur at the third order, where isomorphic projection neurons target sex-specific/dimorphic downstream partners. 17/188 ALPN types send at least 10% of their output to sex-specific/dimorphic third-order lateral horn neurons in the male (Fig 6g). 12/17 are multiglomerular types and strikingly, half are GABAergic vPNs, such as the inhibitory VA1v vPN which sends >50% of its output to sex-specific/dimorphic neurons. These connections are primarily in the anteroventral lateral horn (LH), consistent with previous studies of pheromone representation^126^ (Fig 6g).

We would predict that pheromone PNs would have many dimorphic partners and this is mostly the case (Fig 6h) but pheromone-responsive DA1 lPNs, VA1v adPNs, and DL3 lPNs have a low fraction of sex-specific/dimorphic partners. This can be explained because unlike most multiglomerular PNs, these uniglomerular PNs make strong output connections in the mushroom body, the principal site of learning and memory^127–129^ in addition to the LH. We see no obvious sex differences in the output of pheromone PNs in the mushroom body (Fig S6e). A detailed analysis of olfactory input to the mushroom body finds no obvious sex differences (Pleijzier *et al.,* ms in prep) but there are notable changes in dopaminergic punishment and reward signals (Silva *et al.*, ms in prep).

To begin investigating the flow of pheromone information across the brain and VNC, we followed the excitatory outputs of 4 glomeruli (Fig 6i). DA1 and DL3 detect 11-cis-vaccenyl acetate (cVA), a male-secreted sex pheromone that acts as a female aphrodisiac and promotes aggression in males^114,130^; VA1v and VA1d detect ligands present in both sexes that likely act as aphrodisiacs for males^131^. We see a separation between cVA-specific (LH007m, LH008m) and multi-pheromone (AL-AST1, LHAV1a1, CB3356, CB3359) pathways, consistent with electrophysiological recordings from some of these neurons^26,116^. One novel observation is a dedicated cVA pathway that directly inhibits courtship song (Fig 6i): this may help prevent courtship displays towards other males^101^. When cVA is integrated with other pheromones, the descending pathways broaden. Unique combinations of descending neuron activation will likely enable dynamic pheromone regulation of a range of behaviours. An in-depth analysis of the logic of pheromone processing will be presented elsewhere (Beckett, Morris *et al.*, ms in prep).

### Gustatory system

Like volatile olfactory pheromones, non-volatile pheromones detected by gustatory receptor neurons influence sexually dimorphic behaviours, signalling sex and species identity^132–134^. However, their detection by GRNs usually requires physical contact, as occurs when male flies touch or lick potential mates during courtship^135^. GRNs fall into six anatomically distinct subclasses primarily based on their precise location on the mouth parts, legs and wings^136^ (Fig 7a,b). A detailed typing and cross-matching of all gustatory sensory neurons is reported in our companion paper^137^; combining this with identification and analysis of proboscis motor neurons reveals how sensory input is transformed into the motor programs underlying feeding and foraging behavior^138^.

**Figure 7.**
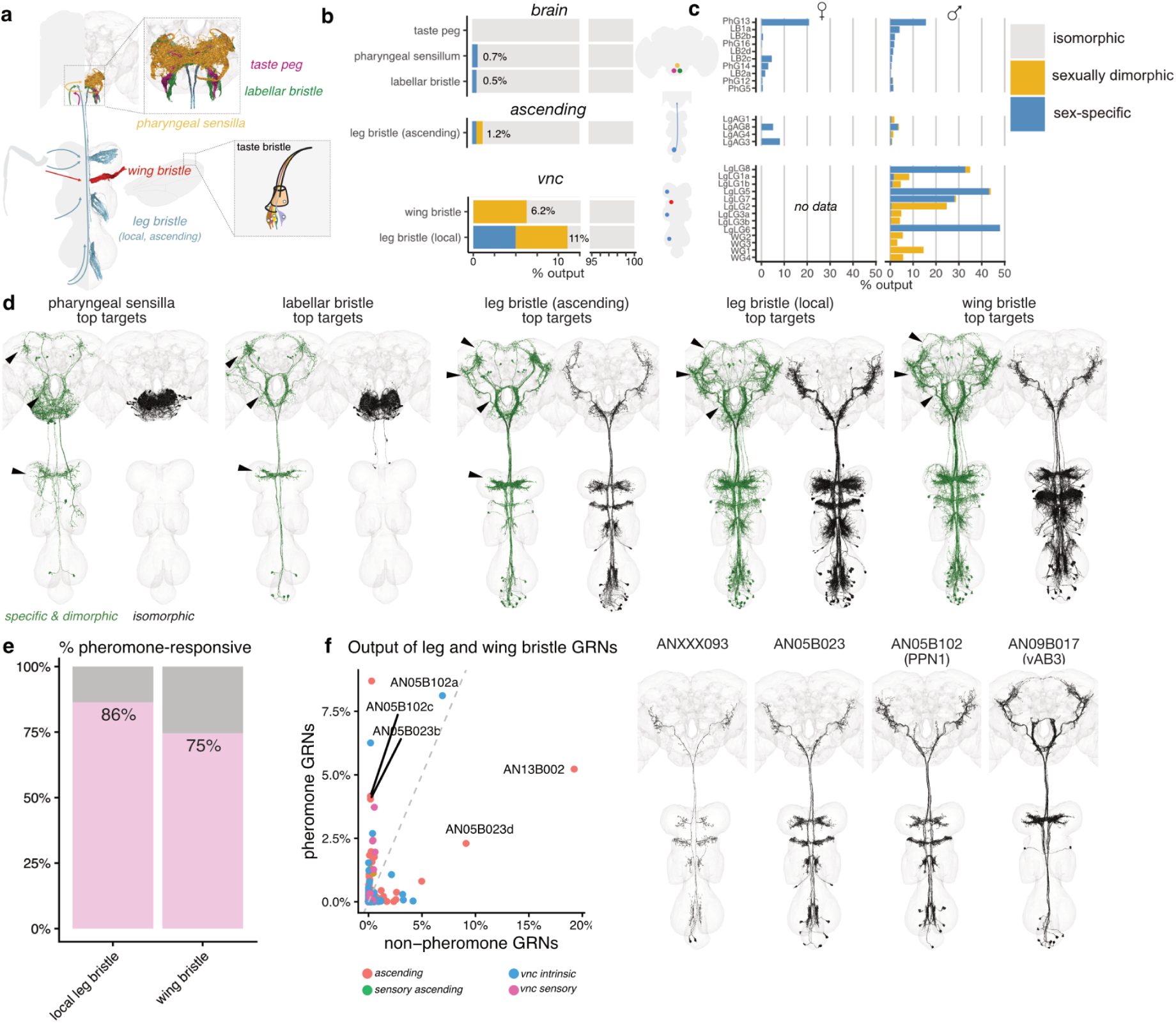
Dimorphic second-order gustatory neurons separate contact chemosensation from taste. **a** Schematic representation (left) and 3D representation of taste sensory subclasses. Taste peg, labellar bristle, and pharyngeal sensilla GRNs (top inset) enter the nervous system via nerves targeting the brain; leg and wing bristle GRNs enter via the VNC. Bottom inset: schematic representation of taste bristle external structure. **b** Output-normalized proportion (%) of GRNs output to types annotated as sex-specific (blue) or sexually dimorphic (yellow) according to subclass and anatomical location: brain (top), ascending (middle), VNC (bottom). **c** For each subclass, the main GRN types targeting sex-specific and sexually dimorphic partners in females (left) and males (right).Types with >1% dimorphic outputs in either sex are shown. See Fig S7a for subclasses. **d** Visualisation of the top ten sex-specific and sexually dimorphic (green) versus isomorphic (black) downstream partner cell types of taste sensory subclasses. Sex-specific and sexually dimorphic neurons extend distinct projections to AVLP, SMP, SLP, SIP, FLA, LegNpT1 (black arrows and Fig S7c). **e** Input-normalized proportion of putatively pheromone-responsive versus non-pheromone responsive neurons in leg and wing GRN subclasses according to receptor-type mapping. **f** Left: Downstream partners of putatively pheromone-responsive and non-pheromone responsive local leg and wing GRNs, coloured by superclass. Percentages are output-normalized. Within-type connections are removed; Right: AN types selective for pheromone-responsive GRN input include a single pair of ANXXX093 neurons and 3 serially homologous cell types, AN05B023, AN05B102, AN09B017, each containing multiple subtypes recorded in neuprint,

Foreleg gustatory sensory neurons have dimorphic axon projections^139^ but we otherwise see minimal dimorphism^137^. However, there are clear differences in downstream circuit organization. We first identified the proportion of sex-specific/dimorphic downstream outputs for each GRN subclass (Fig 7b,c). Local leg bristles engaged most strongly with dimorphic circuits (11% of outputs in males); wing bristles also had a high fraction (6%) but ascending leg, pharyngeal and labellar sensory neuron neurons were all much lower (∼1%). Early gustatory processing is therefore more dimorphic in the VNC than the brain. The more limited dimorphic connectivity in the brain is concentrated in a small subset of GRN types (8/42 types in males, Fig 7c). These data suggest a clear anatomical division: information entering via a larger number of gustatory information channels associated with the mouth parts directs general sex-shared behaviours like feeding, while a more specific set of receptors in the rest of the body is focussed on sex-relevant cues.

We next compared the anatomy of sex-specific/dimorphic (green) versus isomorphic (black) second-order gustatory neurons in the male CNS (Fig 7d). The isomorphic downstream partners of mouthpart sensory neurons are confined to primary taste processing areas in the gnathal ganglion (GNG). In contrast, their dimorphic partners immediately project widely across the central brain and VNC (Fig 7d, Fig S7b) including to higher brain areas characterised by high density of dimorphic connections (Fig 3k, Fig 4g). Dimorphic partners of nerve cord GRNs target these same brain areas and show some specificity for foreleg neuropils (likely associated with male tapping). Neuroanatomical comparison of dimorphic and isomorphic second-order gustatory neurons in females showed similar trends and included some female-specific taste projection neurons (Fig S7c).

### Global analysis of sexually dimorphic connectivity

We have seen that the *Drosophila* CNS consists of isomorphic, dimorphic, and sex-specific neurons (Fig 3) and examined how these are deployed in some specific circuits (Figs 4-7). We now investigate how these dimorphic elements contribute to wiring differences across the whole male and female central brain. We first developed a simple statistical approach to compare edges (i.e. connections) in the connectome graph. Although this explicitly considers the variability of connections within and between individuals, we found that expert review of the nodes i.e. the neurons themselves was essential to draw robust conclusions. We ultimately concluded that about 6% of connections in the male brain (and 1% in female) are dimorphic and that these involve 18% of neurons in male (and 8% in female).

#### A connectivity-based approach to find dimorphic edges

To provide a baseline for comparison we computed the rates at which the three categories of neurons from Fig 3 connect with sex-specific/dimorphic partners: the rates of 39% (for sex-specific neurons) and 20% (for dimorphic neurons) indicate strong interconnectivity (Fig 8a; see Fig S8a for female). In contrast, isomorphic neurons have 4% sex-specific/dimorphic partners; this is very similar to the fraction of sex-specific/dimorphic neurons in the brain (4.8%) suggesting that they may have no preference either way.

**Figure 8.**
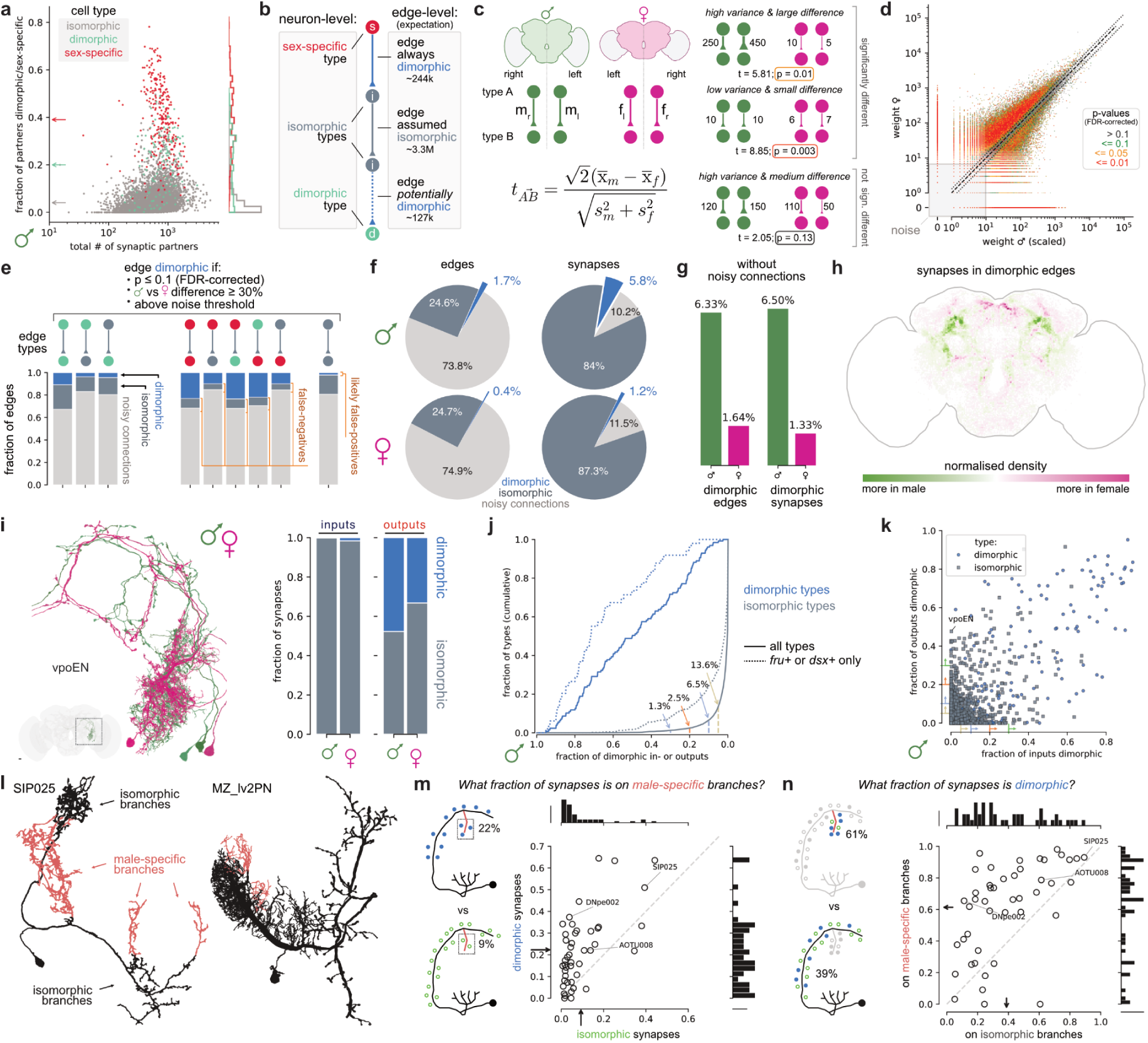
Sexual dimorphism in brain connectivity. **a** Dimorphic up- and downstream synaptic partners as a fraction of total synaptic partners for all cross-matched male cell types. Arrows on the y-axis indicate means for iso-, dimorphic and male-specific types. **b** Illustration of neuron-level dimorphism and expected resulting edge dimorphisms. **c** t-statistic to find significantly different edges between male and female. The denominator is a pooled standard deviation combining the separate standard deviations for male and female edge weights (see <u>Methods</u> for details). **d** Male (scaled) versus female type-to-type edge weights coloured by false discovery rate (FDR)-corrected p-value. Dotted envelope demarcates a 30% difference in connection weights. **e** Fraction of dimorphic, isomorphic and noisy connections (male + female) broken down by edge type. False-positives and -negatives are labeled based on expectation as laid out in a. **f** Proportion of edge types by total number (left) and by synapse count (right) in males (top) and females (bottom). False-positives and -negatives (g) were re-assigned to isomorphic and dimorphic edges, respectively. **g** Fraction of dimorphic edges/synapses not counting noisy connections. **h** Male-female differences in spatial distribution of synapses in dimorphic edges. **i** Example of an isomorphic cell type (vpoEN) with a large fraction of dimorphic outputs (by synapse count) in both male and female. **j** Fraction of iso- and dimorphic types with at least X% dimorphic in- or outputs (by synapse count) in male. **k** Fraction of dimorphic in-versus outputs (by synapse count) per cell type in the male. Arrows correspond to labeled thresholds in panel k. **l** Examples for dimorphic cell types that were split into isomorphic and male-specific branches. **m**,**n** For each split type, the fraction of total synapses from isomorphic and dimorphic connections found on male-specific branches (m), and the fraction of dimorphic synapses found on male-specific vs isomorphic branches (n). Arrows indicate the means along the respective axes.

We began our connection analysis by converting male and female brain connectomes to a pair of weighted graphs. 8,258 nodes represent cell types (cross-matched or sex-specific), while 3.74M edges are defined by the number of synaptic connections between types. Before continuing we stopped to make some simple predictions (Fig 8b). First, we imagined that all connections involving sex-specific neurons should be dimorphic (since they cannot exist in the opposite sex). Second, morphologically dimorphic neurons would have a lower rate of dimorphic edges (mirroring our observations in Fig 8a). Finally, we predicted that most if not all edges between isomorphic cell types are isomorphic: any edges flagged as dimorphic would probably result from biological variability or technical noise.

Crucially, we have previously shown that the distribution of connectome edge weights is highly skewed^31^: most synapses belong to strong edges that are reliably observed within and across datasets; but the majority of edges are weak and unreliable. We therefore defined a noise threshold for each dataset and ignored weak edges below these weights (<8 for male CNS, <11 for FAFB/FlyWire, Fig S8e), leaving a graph with 721k edges. Synapse counts also differ systematically across datasets mostly due to technical factors^31^; we therefore scaled the male connection strengths down to match the female (see <u>Methods</u> and Fig S8d). We then computed a t-statistic where the mean difference between male and female edge weights is divided by the within-sex (left vs right) standard deviation (Fig 8c). Intuitively, this highlights connections with large differences between datasets and low variability between the two hemispheres of the same dataset (Fig 8d). Given the large number of tests, we used false discovery rate (FDR) correction^140^ to balance sensitivity and specificity (see <u>Methods</u> and Fig S8d).

#### Contrasting morphology- and connectivity-based approaches

This procedure allowed us to label each edge as either dimorphic or isomorphic (or noisy) (Fig 8e; Fig S8h) quite independently of our annotations from Fig 3. Discounting noisy connections, 68% (32k) of edges involving male-specific neurons are flagged as dimorphic. This is satisfyingly high even if it does not match the 100% we propose in Fig 8b; the main reason is that some connections in the male were too variable across hemispheres to be distinguished from zero synapses. For dimorphic cell types, 23% (5.4k) of edges were flagged as dimorphic; this is consistent with our observations that in most cases only part of their arbours are dimorphic.

Finally, we examined edges involving two isomorphic cell types: 11% of these edges (72.2k) were flagged as significantly different between sexes. We worried that some dimorphic cell types might have evaded our initial neuron-level search (Fig 3). But reviewing the 500 isomorphic types that showed up most frequently led to <5 changes in dimorphism status. Rather, most apparent differences in male-female connectivity were explained by segmentation issues or inconsistent cell type definitions across datasets. Given the available evidence that most of these edges are false-positives (i.e. not actually dimorphic) we excluded them from further consideration. Our account therefore represents a lower bound on the extent of dimorphism in the brain.

Having completed this analysis, we conclude that 6.3% of edges in the male and 1.3% of edges in the female central brain are dimorphic with similar numbers for synapses (Fig 8f,g, ignoring noisy edges). There are therefore slightly more dimorphic connections than sex-specific/dimorphic neurons in males, but the opposite is true in females (compare Fig 3i). Identification of all dimorphic edges enables many additional analyses: for example the highly punctate spatial distribution of the contributing synapses (Fig 8h) reinforces our earlier observations about dimorphic hotspots (e.g. Fig 3k).

We can also now quantify the extent to which the sex-specific/dimorphic neurons interconnect with the rest of the brain. Many dimorphic edges flagged in the analysis involve one partner cell type categorised as morphologically isomorphic (Fig 8k,l; see also Fig S8i,k). Interestingly, if those isomorphic neurons are *fru+/dsx+* they participate in many more dimorphic connections (Fig 8j). We speculate that *fru/dsx* may control guidance molecules that make isomorphic neurons a target for neurites of their sex-specific/dimorphic partners. For example, *fru+* vpoEN neurons (Fig 5) – which link male courtship to song to female receptivity by promoting opening of the vaginal plate^53^ – do not have clear sex differences in morphology (Fig 8i) but 48% of their output connectivity is dimorphic (Fig 8j). We propose the term secondarily dimorphic for neurons with dimorphic connectivity without dimorphic morphology; accounting for them means that 17.9% of male central brain neurons are directly or indirectly dimorphic (versus 7.8% for the female).

How do neurons make dimorphic connections? We previously saw a case where dimorphic connections are highly concentrated in dimorphic arbours (DNpe002 in Fig 4f) and another where the relationship was less clear (AOTU008 in Fig 3e-g). We now extended our analysis to a further 46 cell types where we could resolve dimorphic axons or dendrites (Fig 8l, see <u>Methods</u> for details). Dimorphic synapses were almost always enriched on male-specific arbours (Fig 8m,n) and this clustering could affect their physiology. However, surprisingly in most cases the majority of dimorphic synapses are not on male-specific arbours (Fig 8m). We speculate that some sex-specific arbours have a developmental function, pioneering a physical connection that serves as staging post for outgrowth of partner neurites.

### Dimorphic neurons are strongly clustered and interconnected

Having defined both dimorphic cell types (nodes) and connections (edges), we investigated the organisational logic of these elements across the connectome. Starting from the type-to-type connectome graph for the male central brain, we obtained hierarchically grouped nodes into clusters of similar connectivity (Fig 9a,b; see <u>Methods</u> for details). This identified two key modes of involvement of the dimorphic and male-specific types in the network structure.

**Figure 9.**
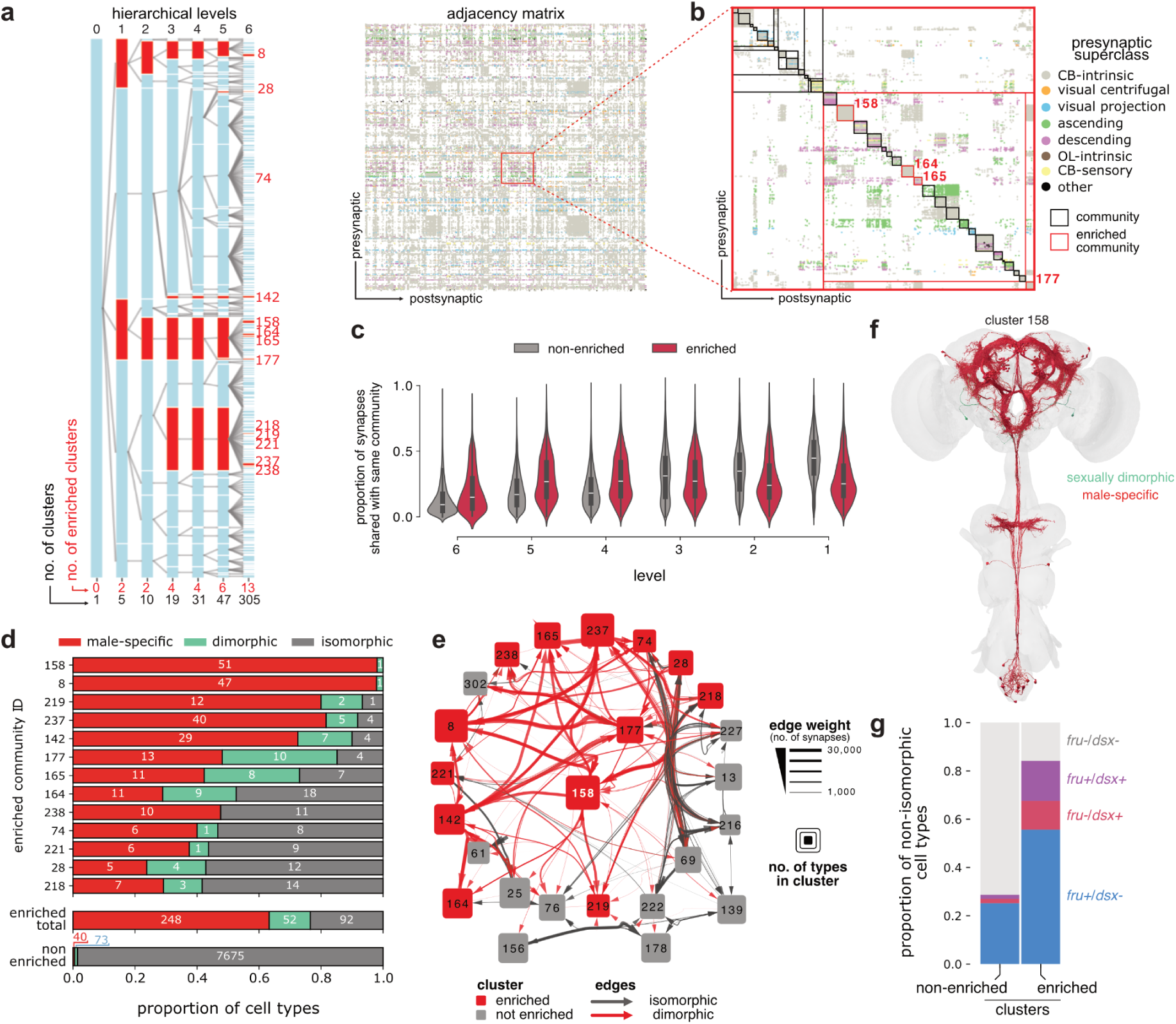
Sexual dimorphism in brain networks. **a** Hierarchical community detection partitions the cell types of the male central brain into blocks of similar connectivity. Clusters at 6 different levels of the hierarchy (left) are derived from the type-to-type connectivity matrix (right). The IDs of male-enriched clusters are shown in red. **b** Zoom-in on inset highlighted in panel a showing the hierarchical partitions in the adjacency matrix, with clusters enriched with non-isomorphic types in red. Rows are coloured by superclass. **c** Proportion of within-community synapse counts across the hierarchical levels. **d** Breakdown of cell type identity in the 13 enriched clusters at level 6. **e** Network graph of the neighbourhood (≥1,000 synapse threshold) around enriched cluster 158. **f** Rendering of enriched cluster 158, the most statistically overrepresented for male-specific or dimorphic cell types. **g** *Fruitless* and *doublesex* expression among non-isomorphic cell types in enriched versus non-enriched clusters.

First, only 13 out of 305 clusters were statistically enriched for dimorphic or male-specific types, constituting 300 of 413 (72%) non-isomorphic types. The enrichment persisted over six hierarchical levels, where 2 clusters remained statistically enriched even at the top level. On average, male enriched clusters connected more densely within their community than cell types in non-enriched clusters across hierarchical levels 4-6 (Fig 9c). Most dimorphic/sex-specific types therefore contribute to distinct subnetworks, and male-specific cell types are the main contributors to this enrichment (Fig 9d). Accordingly, 85% of dimorphic edges involve a type from an enriched cluster on one or both ends of the connection. However, the remaining 113 male-specific/dimorphic types showed a very different network organisation. This population was sparsely distributed across 64 non-enriched clusters and highly integrated into the isomorphic subgraph.

The most enriched cluster (158) contains many male-specific cell types, including many P1 subtypes (Fig 9e,f). A detailed analysis of P1 neurons is presented elsewhere^141^. We found that the other 12 enriched clusters all connect to cluster 158 with at least 1,000 synapses, mainly via male-specific connections (Fig 9e). This cluster exemplifies a second key organisational feature: male-enriched clusters connected to an average of 8.5 other enriched clusters by at least 1,000 synapses, compared to 1.1 for non-enriched clusters. The male-specific subunits of the connectome therefore connect densely within each cluster but are also globally interconnected with other male-specific communities. Finally, the proportion of *fru+/dsx+* neurons is much higher in enriched versus non-enriched clusters (Fig 9g), confirming the long-standing hypothesis that *fruitless* neurons preferentially connect with each other to form circuits regulating male behaviour^122^.

## Discussion

The complete connectome of the *Drosophila* male central nervous system represents a major step forward in the quest to link neural circuits and behaviour. The fully proofread connectome enables end-to-end analysis from eyes to legs.Our rich annotations link over 95% of neurons to previously reported cell types and to functional analysis in the literature.

The number of sex-specific and dimorphic neurons is relatively modest (approximately 5% of the male brain and ∼2% of the female brain, Fig 3i) but they can have major impacts on function. One explanation is that 18% of male brain neurons (8% for female) have significant dimorphic connectivity (Fig 8j). At the cellular level, perhaps surprisingly, we find male-specific neurons to be more common than dimorphic neurons (which have altered morphology), whereas the opposite is true in females. A greater proportion of male-specific versus female-specific neurons is also observed in *C. elegans* although the difference is more extreme (23.6% versus 2.6%)^142^; this consistent bias may reflect selection pressure on male behaviours.

We clearly demonstrate that sexually dimorphic elements are not evenly distributed in the brain but are concentrated in higher brain areas (Fig 3k). This is compatible with early observations of the pattern of *fruitless*-expressing neurons in the brain^122,143^ as well as volumetric differences between male and female brains^63^. However, we can now place these neurons in circuits. We see that they are in deep layers of the connectome between the sensory and motor periphery (Fig 3l). This suggests a hierarchy in which sex differences primarily modify integrative and decision-making areas while sensory detection and the highly tuned motor interface remain more constant. This logic appears distinct from *C. elegans,* where male-specific neurons are principally associated with circuits local to its dimorphic genitalia^142^. It will be very interesting to see which of these patterns apply to circuit evolution across species^144^.

Even within higher brain centres, there is additional structure: dimorphic neurons form synapses in a spatially clustered fashion (Fig 3k) and this can be associated with feedforward and feedback connections in particular sensorimotor pathways such as the visual elements in Fig 4g or the ascending taste neurons in Fig 7d. Sex-specific neurons contribute to multiple wiring motifs including sign inversion by inhibitory interneurons downstream of visual projection neurons (e.g. VES200m, Fig 4n) and switches in partner connectivity (e.g. PVLP207m, Fig 4e,n). In contrast, some sex-shared interneurons appear to act as a node for dimorphic connectivity without showing any sign of dimorphic morphology (vpoENs in Fig 5b,c). Zooming out, we see that the large majority of male-specific/dimorphic neurons form a small number of tightly coupled hubs (Fig 9). While some dimorphic neurons do integrate more sparsely across the connectome, the dominant organisational principle seems to be modules of sex-specific circuits. From an evolutionary perspective it may be easier to evolve such discrete modules in order to produce major changes in sex-specific behaviour. This is also consistent with our observations that the majority of sex-specific and dimorphic neurons originate from a small number of neuroblast lineages (Fig 3j).

We observed a strong correspondence between *fru* and *dsx* expression and dimorphic and male-specific neurons. However, there were potentially interesting discrepancies. Over 1,500 *fru*/*dsx* positive neurons were not dimorphic in the connectome; although technical limitations may play a role, this does raise the possibility that they are regulating factors other than wiring. To give a concrete example, we can confidently assign the cell type LHPV6q1 (pSP-e, FMRFa-WED). It has documented sex differences in its electrophysiological properties including its spontaneous firing rate^52^, but shows no wiring differences in the connectome. Conversely, we observe many neurons where *fru*/*dsx* expression is absent but neurons are dimorphic or sex-specific. Although technical limitations likely account for some of this discrepancy, we can be quite confident in some cases (e.g. 236 neurons in the central brain with known lineage) leaving open the possibility that alternative mechanisms are at play.

Connectomics is entering a rapid growth phase, but for comparative connectomics low sample numbers still pose significant challenges. Lessons learned during this project leave us cautious in some areas but still optimistic overall. We believe that attempts to extract biological differences from direct comparison of two raw connectome graphs are unlikely to be successful. Rather we recommend analysing cell type graphs, capturing the fundamental unit of conservation across connectomes while accounting for interindividual variation^31^. A strong quantitative understanding of that variability (and its technical and biological origins) is also essential. Crucially, we found that considering bilateral consistency of cell type morphology and wiring enabled us to draw robust conclusions about the identity of dimorphic elements. A specific example may be instructive. We previously noted a visual projection neuron (LoVP109/LTe12) present with one exemplar in each optic lobe of the FAFB/FlyWire female brain^30,31^ was missing in the single male optic lobe reported in Nern *et al*^36^. However, we now find this cell in the second optic lobe of the complete male CNS dataset, suggesting a developmental error in the first optic lobe. We estimate this error rate at <1% for neurons in each hemisphere^31^; if errors occur independently in the two hemispheres – as observed here – then the chance of missing an entire cell type is <0.01% or about 3 pairs of neurons in the central brain; this is a very low false positive rate compared with our observation of >1,000 male-specific neurons. Similar principles apply for connectivity, echoing recent work on cross-species comparisons of the larval *Drosophila* olfactory system^145^. Nevertheless, we found that expert curation was required to separate biological signal from cases of technical noise (which is more commonly correlated across both sides of one brain).

Although this is the largest finished connectome to date, limitations remain. First, while our dataset includes the entire male CNS, we were only able to make comprehensive across-sex comparisons to the FAFB/FlyWire female brain^30,31,42^; comparisons to the partially proofread female nerve cord (FANC) dataset^35,146^ were necessarily more restricted. This limitation could be removed by the very recent release of the female Brain and Nerve Cord (BANC) dataset^147^. Although BANC remains a work in progress at the time of writing, the nerve cord is at a more advanced stage and should allow male-female comparisons; nevertheless, we would caution that careful joint cell typing and a quantitative understanding of variation will be essential to draw robust conclusions about sex differences. We see huge value in integrative connectome analysis and already provide powerful tools (https://natverse.org/coconatfly) that enable analysis across all publicly available fly connectomes including the BANC and male CNS.

Second, the connectome provides the structural wiring diagram, but of course functional analysis will be required to confirm predictions about differences in information flow and their impact on behaviour. We are heartened by the recent success of brain-scaled simulation studies^148,149^. In the near future manipulation of neuronal connectivity and activity *in silico* may be used to understand the impact of the dimorphic elements we observe and to prioritise the many experimental hypotheses that emerge from connectome analysis.

Third, the connectome is the end state of a complex but highly stereotyped process of development. In some cases we may have failed to identify developmentally homologous neurons because their structure and connections are too divergent. Integration with molecular and developmental studies may be essential to resolve these issues^150–152^. However, by providing the most detailed picture yet of the result of sexually dimorphic development it should prove invaluable for studies examining the molecular specification of brain wiring^153^.

Similarly, this connectome represents a snapshot in the lifetime of an animal. This study confirmed our previous observations that the majority of the wiring diagram is stable^31^. However, even strongly innate dimorphic behaviours can be modified by experience^154,155^. Building on developments in comparative connectomics – including this study – there will be exciting opportunities to learn about the footprint that experience leaves in the wiring diagram of the brain.

## Supporting information

Higher Resolution Main Figures 1-9

Higher Resolution Supplementary Figures S1-S9

## Acknowledgements

This work was principally funded by Wellcome Trust collaborative awards (220343/Z/20/Z and 221300/Z/20/Z) to GSXEJ, GMR, SW and GC, by the Howard Hughes Medical Institute at Janelia for the FlyEM project team led by SB for proofreading, annotation and analysis of male CNS neurons, and core funding from the Medical Research Council (MC_U105188491) to GSXEJ. IRB was supported by Boehringer Ingelheim Fonds and the Cambridge Commonwealth, European and International Trust. Development of the natverse including the coconatfly package has been supported by the NIH BRAIN Initiative (grant 1RF1MH120679-01), NSF/MRC Neuronex2 (NSF 2014862/MC_EX_MR/T046279/1). We thank Sven Dorkenwald and Arie Matsliah for sharing data artefacts for the new synapse predictions for FAFB/FlyWire^156^, and members of the MRC LMB Scientific Computing group for assistance with compute and web infrastructure. We thank Hiroshi Shiozaki and Davi Bock for comments on the manuscript.

Rights retention: For the purpose of open access, the MRC Laboratory of Molecular Biology has applied a CC BY public copyright licence to any Author Accepted Manuscript version arising. This article is subject to HHMI’s Open Access to Publications policy. HHMI lab heads have previously granted a nonexclusive CC BY 4.0 license to the public and a sublicensable license to HHMI in their research articles. Pursuant to those licenses, the author-accepted manuscript of this article can be made freely available under a CC BY 4.0 license immediately upon publication.

## Author Contributions

KJH & ZL prepared the sample. WQ, AC, CSX & HFH imaged the sample. SP, ETT, TPi & SS aligned the EM data. SB, MJ & CO produced the automated segmentation. IRB, MC, ECM, S-yT, AMCF, MiC, MG, CO, TPa, FeL, BJM, IA, PB, BSC, IdHV, CRD, SFM, MAF, IH, GPH, LK, JK, SAL, ML, AL, CAM, IM, CM, EM, NO, AP, BP, EMP, AR, JRS, RJVR, ALS, LAS, LSC, CS, ST, AT, JJW, HW, EAY & TY proofread the segmentation. GBH & CO produced the automated synapse detections. SB, AN, DA, GBH & ASB identified neurotransmitters. PSc, AN, JB & SC developed spatial transforms. IRB, MC, PSc, ECM, AN, S-yT, AMCF, MG, TS, CRW, FeL, BJM, MWP, VS, GB, RJB, PB, BC, IdHV, GD, ED, MdS, CRD, KE, LK, IM, EM, AP, AR, IT, HW, CR & GSXEJ typed and annotated cells. SB, IRB, MC, PSc, ECM, AN, S-yT, AMCF, MiC, MG, CO, TPa, TS, KDL, CM, CK & GSXEJ provided quality control. SB, PSc, SP, WTK, ETT, LEB, JH, FrL, JC, PMH, DJO, RS, LU, AZ, SS & GSXEJ developed software tools. SB, IRB, PSc, AMCF, LEB, FrL, PMH & EMR developed visualizations and presentation tools. IRB, MC, PSc, ASC & GSXEJ drafted the main text with input from SB, AN, AMCF, SSM, CR, MBR, & GMR. SB, MC, PSc, MJ, AN, SP, DA, GBH, JH & MBR wrote the methods. SB, IRB, MC, PSc, AN, AMCF, ASC, MG, GBH, FK, TS, CRW, VS, YY, MD, SSM, EMR, PSe, AZ, MBR & GSXEJ analyzed the connectome and designed figures. MC, PSc, ECM, GWM, SR, SS, MBR & GSXEJ provided supervision. VJai, WK, SW, GMC, GMR & GSXEJ acquired funding. SB, SP, SMP, RG, VJay, WK, JF, LKS, GMC, HFH, GMR & GSXEJ guided the project.

## Supplementary Figures

**Figure S1.**
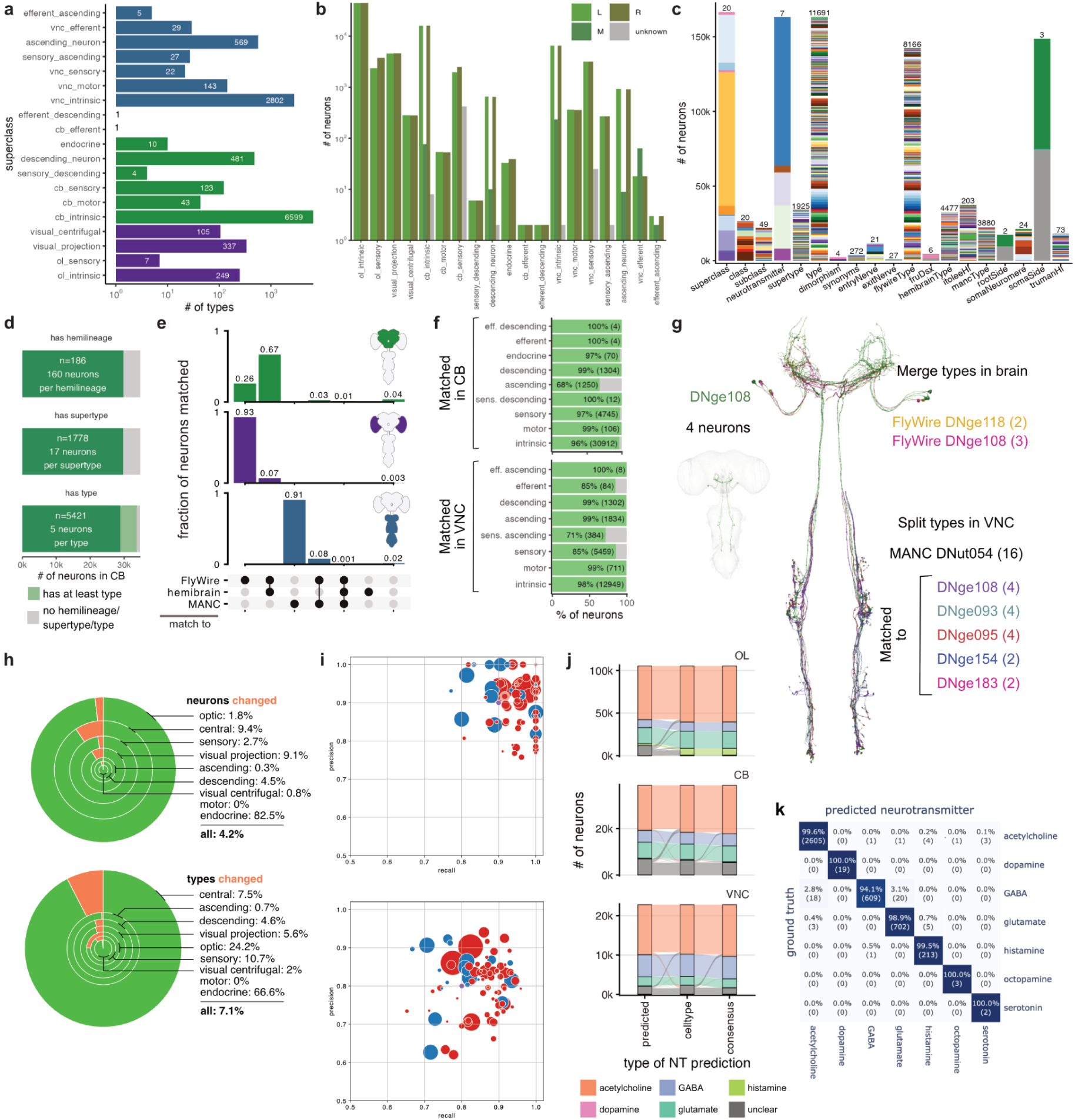
Annotation, cross-matching and synapse and neurotransmitter assignment, related to figure 1 and figure S9. **a** Number of types per superclass and per CNS region. **b** Number of neurons by soma or root side per superclass. **c** Number of neurons with annotations for several categories.The number of unique annotations for each one is shown at the top of the bar. **d** Number of neurons in the central brain with `itoleeHl`, `supertype` and/or `type` annotations. The number of distinct annotations, and the average number of neurons per category is shown in the bar. **e** Fraction of neurons cross-matched to existing datasets per CNS region. **f** Number of cross-matched neurons in the central brain and VNC, per superclass. The number of matched neurons and the corresponding percentage is shown. **g** The descending neuron type DNge108 was matched to 2 types in FAFB/FlyWire and a subset of neurons for type DNut054 in MANC. The remaining 12 neurons in DNut054 match 4 other types in male CNS. **h** FlyWire cell types updated via cross-matching with the male CNS broken down by superclass. Top: percentage of neurons for which the type changed; bottom: percentage of types for which at least one neuron changed. Surface area of wedges corresponds to the total number of neurons/types in that category. **i** The precision and recall of synapse T-bar predictions (top) and pre-post connections (bottom) in each region of interest (ROI). Markers are scaled by ROI size and color differentiates between brain (red) and VNC (blue) neuropils. **j** Change in neurotransmitter prediction for `predictedNt`, `celltypeNt` and `consensusNt` for each neuron in the CNS subregions. **k** Neuron-level confusion matrix for neurotransmitter prediction (`predictedNt`) on the held-out test set.

**Figure S2 related to figure 2.**
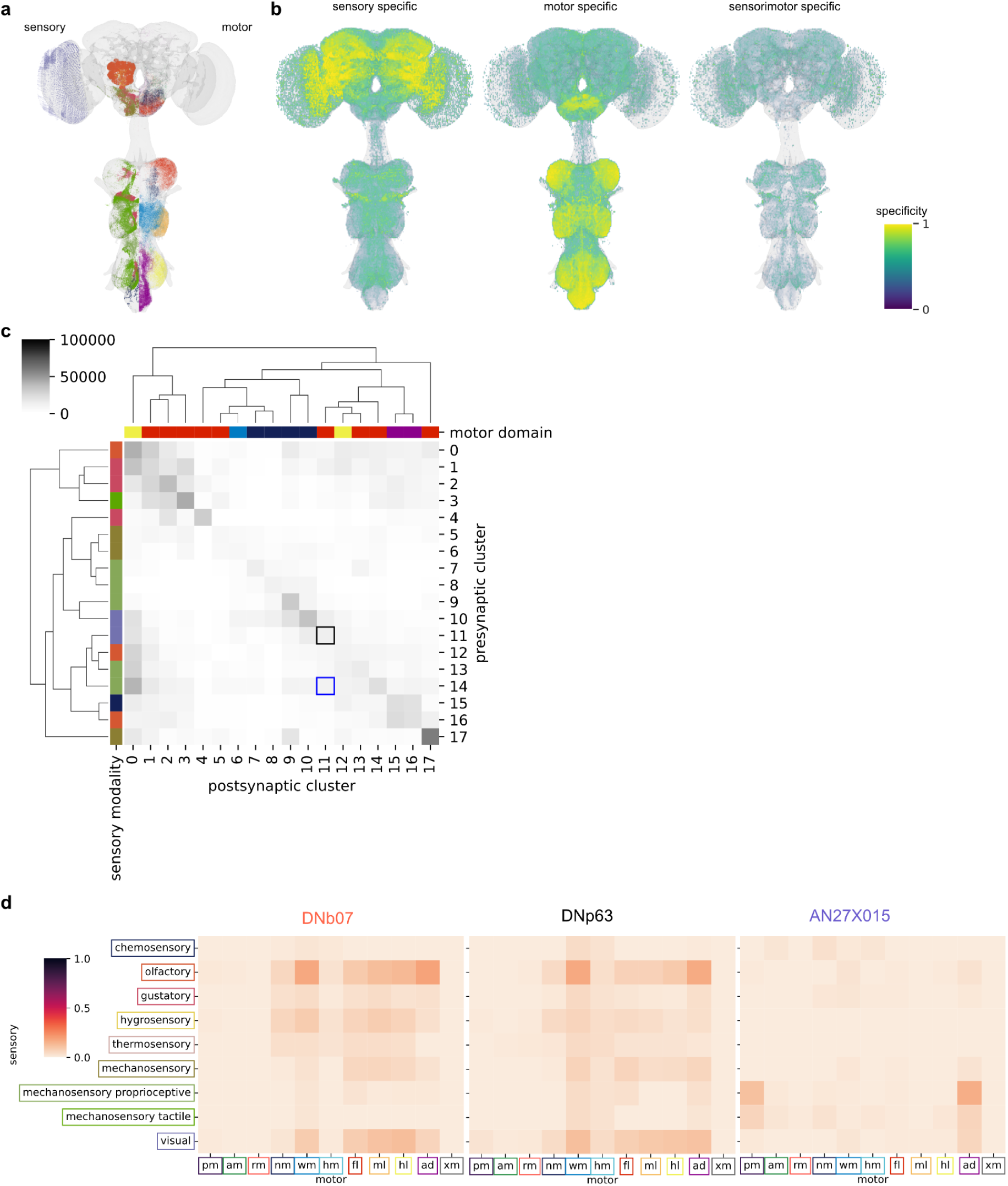
**a** Synapses from sensory (left) or motor groups (right) in the CNS. **b** Maximum intensity projection of synapses of neurons in the CNS that are specialised in flow for left: sensory group, middle: motor group, right: specific sensory-to-motor pairings. **c** Connectivity between clusters in Fig 2h clustered by cosine similarity. Mode preferred sensory modality and motor domain are labeled. Black and blue highlights indicate the elements containing the DN-DN and AN-DN connection in Fig 2m, respectively. **d** Sensory-motor flow for the neck connective types in Fig 2m.

**Figure S3.**
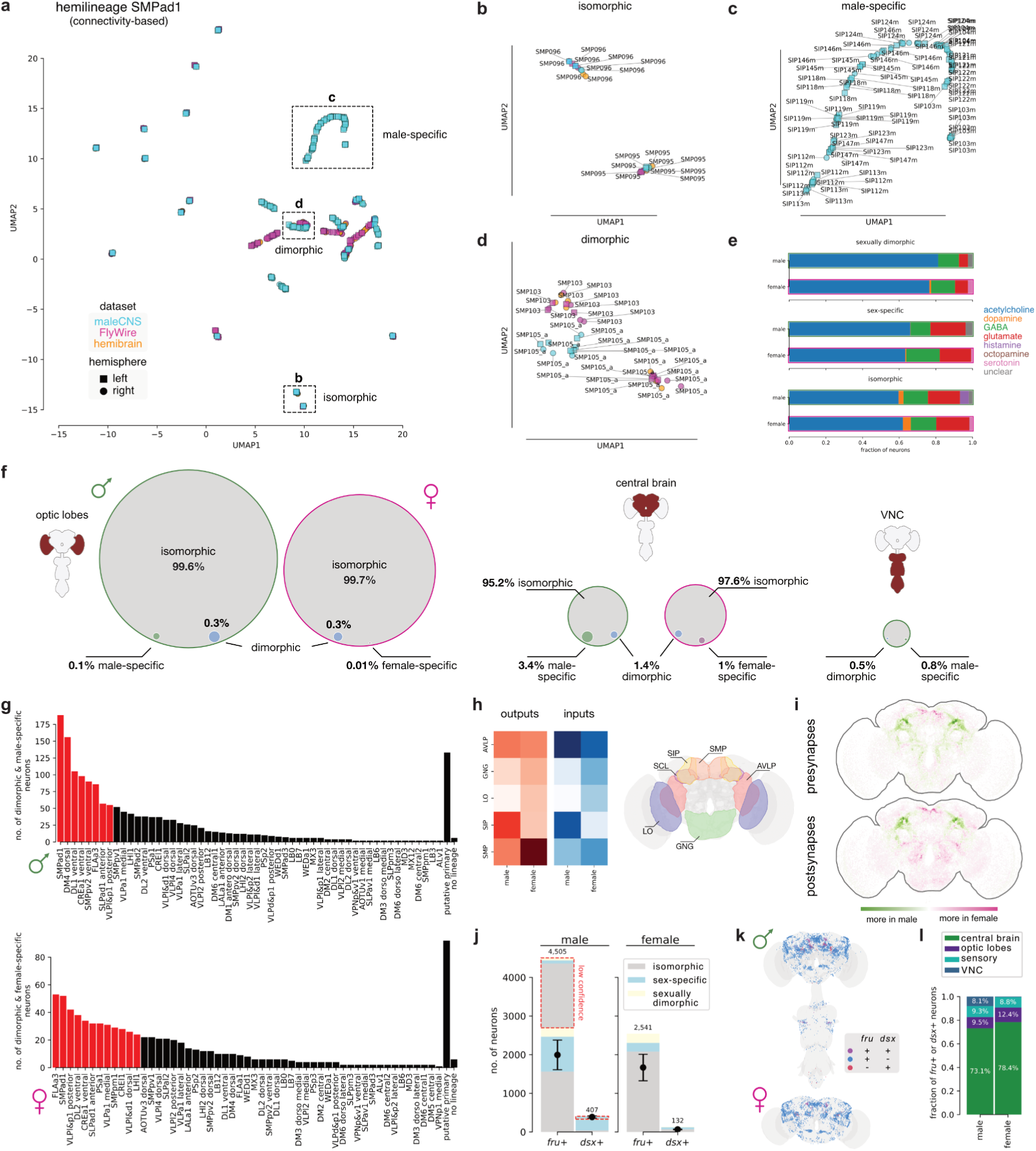
Additional dimorphism analysis, related to figure 3. **a** UMAP embedding of connectivity-based cosine distance for neurons of the SMPad1 hemilineage from three connectomes/five hemispheres: male CNS, FlyWire and hemibrain. **b-d** Zoom-ins on insets labeled in a showing isomorphic (b), male-specific (c) and dimorphic (d) cell types. **e** Distribution of predicted neurotransmitters for dimorphic, sex-specific and isomorphic neurons. **f** Pie charts showing the fraction of dimorphic and sex-specific neurons across optic lobes, central brain and VNC. Relative sizes correspond to the number of neurons. For the VNC, our annotations in the male include 15 dimorphic types (47 neurons) from previous publications^50^ that are either VNC-intrinsic (i.e. do not have a correlate in the female brain volume), or ascending neurons for which the axonal arbors in the brain are too small to identify them in FAFB/FlyWire. Note that VNC dimorphism labeling is partial and hence represents a lower bound. **g** Number of dimorphic and sex-specific neurons per hemilineage in male (top) and female (bottom). Lineages coloured in red collectively produce ≥50% of dimorphic and sex-specific neurons. **h** Top 5 neuropil with the most dimorphic/sex-specific synapses. **i** Difference between distribution of dimorphic/sex-specific synapses in males and females. **j** Breakdown of *fruitless* (*fru+*) and *doublesex* (*dsx+*) annotations by dimorphism. This represents an upper boundary because most of the light-level data labels large populations of neurons which makes identifying individual neurons difficult. Dotted red outline indicates low-confidence annotations in the male. Pointplots in black represent mean and standard deviation of expected counts based on prior literature. **k** Cell bodies of *fru+* and *dsx+* neurons in male (left) and female (right). A small number of *fruitless* and *doublesex*-expressing neurons were annotated in the VNC based primarily on previous annotations from the MANC dataset^58,150^. **l** Distribution of *fru+* and/or *dsx+* neurons across main brain regions. The majority (73%) of *fruitless*/*doublesex* annotations are in the central brain, 8.1% are in the VNC, and 9.5% are in the optic lobes.

**Figure S4A.**
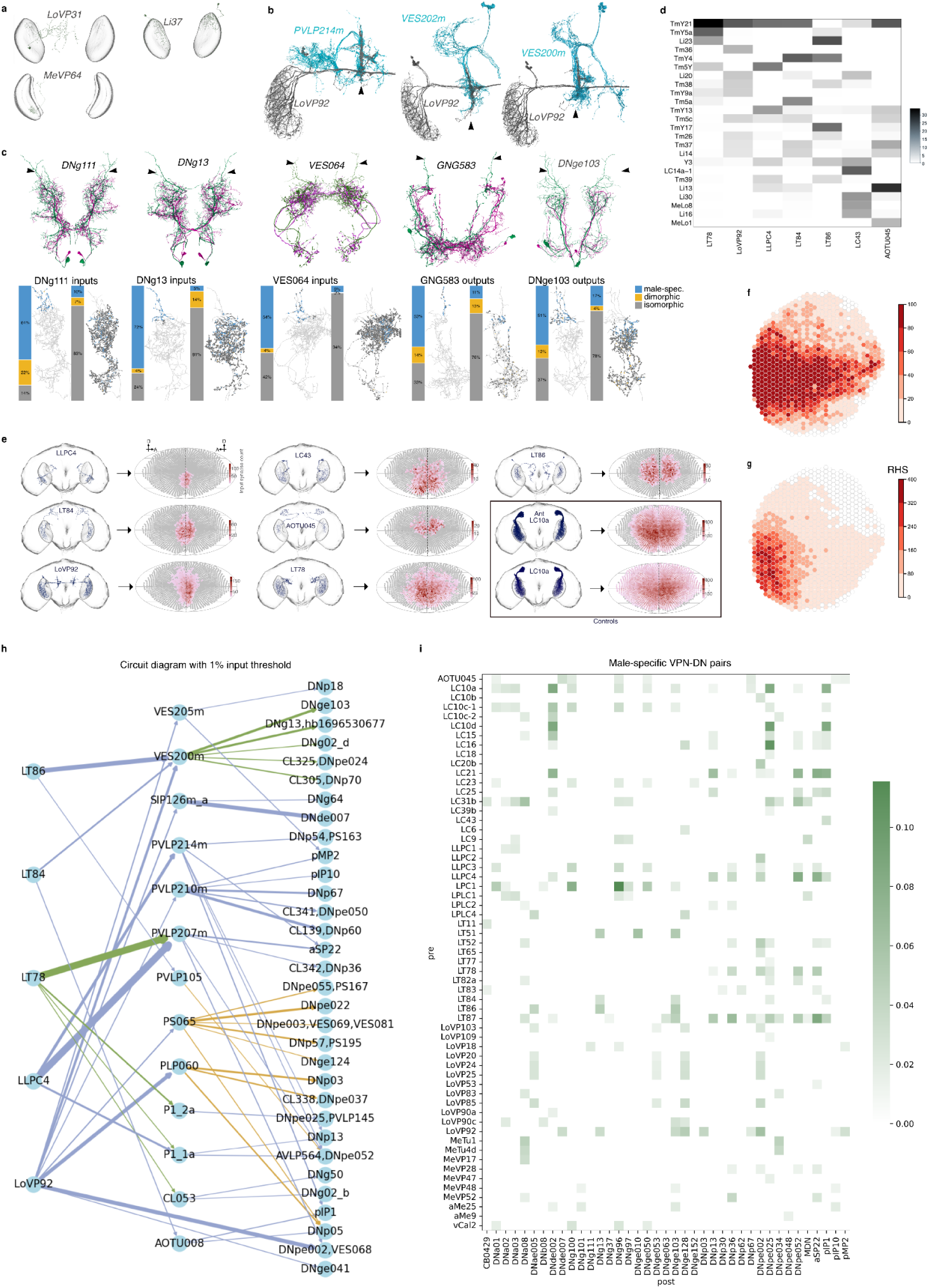
Additional visual system analysis, related to figure 4. **a** Unmatched outlier types not considered male-specific because not left-right symmetrical. Note that Tm40 is currently classified as isomorphic. However, FAFB/FlyWire has lower cell count compared to the male CNS (41 versus 180 ids). Consequently, it was excluded from the analysis shown in Fig 4c,l. **b** Male-specific types converging on the male-specific node defined by LoVP92 axons. **c** Dimorphic types converging on the male-specific node defined by LoVP92 axons. Stacked bar charts indicate the proportion of dimorphism status among connection partners of the dimorphic types. **d** Heatmap showing the strongest inputs from optic lobe intrinsic neurons to frontally-biased VPNs. **e** Examples of frontally-biased VPNs. Spatial coverage heatmaps show input synapse distributions mapped onto a Mollweide projection of the right compound eye’s visual field. Color scale bars show input synapse count. **f** Spatial map showing all inputs to TmY21 in the ME. **g** Inputs to male-specific, dimorphic and secondary dimorphic frontal types are biased towards columns devoted to the frontal field of view. **h** All pathways from VPNs in Fig 4l, to DNs with known functions or sexual dimorphism, within two synaptic hops, where all connections are stronger than 1% input. **i** Effective connectivity based on male specific pathways, involving at least one dimorphic or sex-specific cell type, within two synaptic hops, where each connection is stronger than 1% input, from all VPNs, to DNs with known functions or sexual dimorphism. The colour is based on the sum of direct connection strength, and the square root of the products of weights in the two-hop pathways.

**Figure S4B.**
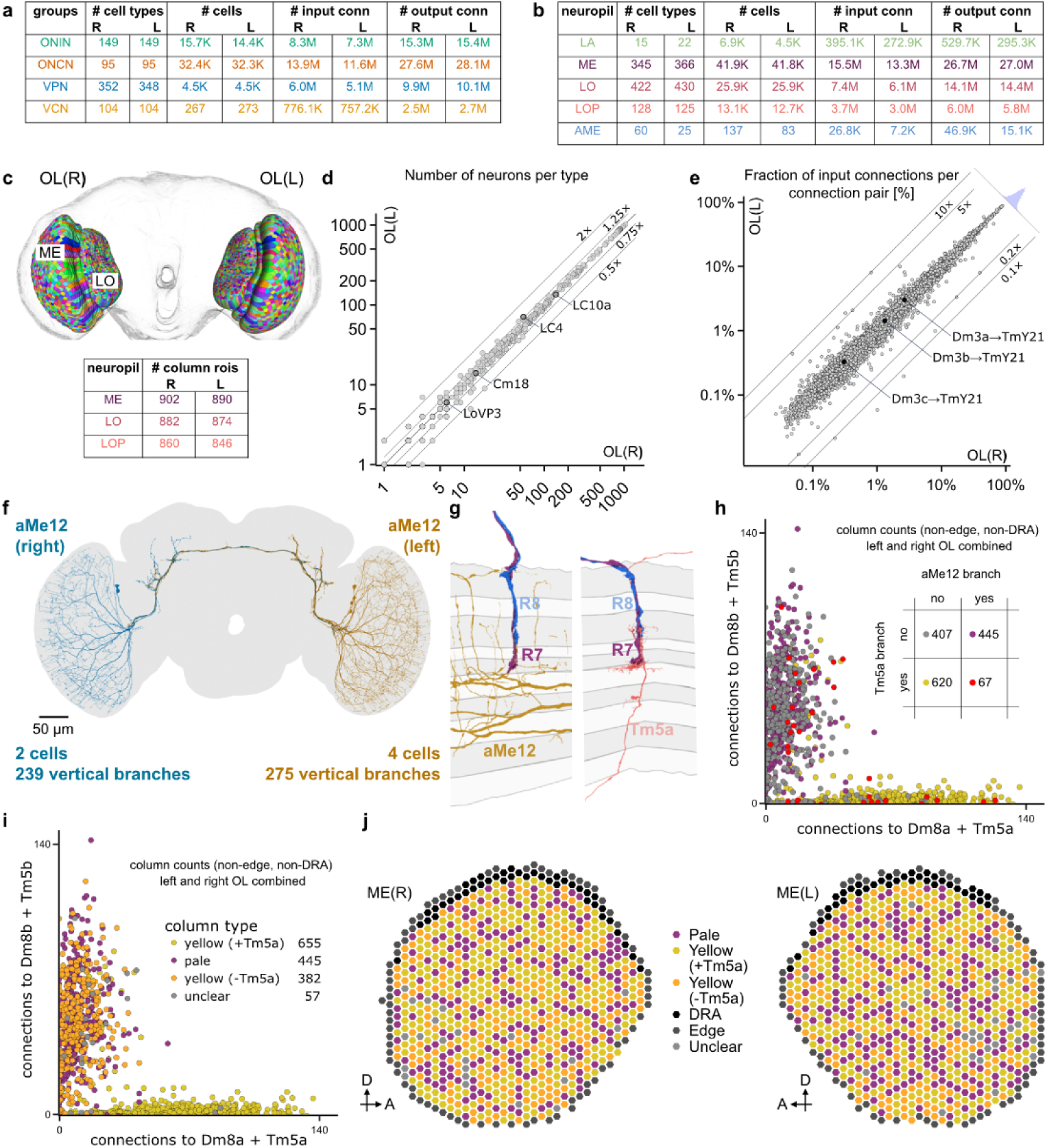
Complete inventory of optic lobe cell types across both brain hemispheres of the male CNS data set, related to figure 4. **a** Comparative summary of cell type groups showing numbers of cell types, cells, input and output connections across both optic lobes (Cell type groups follow Nern *et al.* 2025, with intrinsic neurons of the optic lobe, referred to as OLIN in Fig 4b, divided into ONIN and ONCN, Optic Neuropil Intrinsic and Connecting Neuron groups.). The table omits cell types not classified as ONIN, ONCN, VPN or VCN (the ‘other’ group in Nern *et al.* (2025), see <u>Methods</u>). Of the 701 cell types present in both hemispheres, 4 are only found on the right side, and 1 is only found on the left side. **b** The distribution of cells, cell types and connections across the optic lobe neuropils (LA: lamina; ME: medulla; LO: lobula; LOP: lobula plate; AME: accessory medulla). Some differences between left and right hemispheres appear to be due to differences in the position of neuropil boundaries between the two sides. **c** Column ROIs were updated in the right optic lobe, and developed for the left optic lobe, following the methods of Nern *et al.* (2025). This image shows a frontal view of the brain, highlighting the ‘patchwork’ of column ROIs across the main optic lobe neuropils. The table shows the number of column ROIs in the three columnar optic lobe neuropils. Bilateral layer ROIs are also implemented in the neuPrint database to facilitate analysis (not shown). **d** Scatter plot comparing cell counts per type between left and right optic lobes. Nearly all cell types lie near the unity line, indicating similar numbers of cells / type in both hemispheres. **e** Scatter plot comparing the number of connections per connection type (i.e. ordered pair of connected cell types), quantified as the percentage of connections relative to the total input connections of the postsynaptic type, in the left and right optic lobe. The histogram (blue), aligned with the plot, shows the distribution of ratios of left and right optic lobe input percentages. For cell types with synapses in both optic lobes, the left and right hemisphere cells were treated as different types. Connections show strong approximate bilateral symmetry across the diversity of synaptic connections between optic lobe types. The weights of the indicated connections are very similar between the two male optic lobes but differ in female flies (see Fig 4). **f** Rendering of the aMe12 neurons, whose fine arbors were systematically proofread to provide an improved anatomical marker for typing columns into pale/yellow (Kind *et al.* 2021; Nern *et al. 2025*). The number of aMe12 cells differs between the two hemispheres but the number of vertical branches (see next panel) is similar. **g** Examples of anatomical features used to type medulla columns – generally, columns with an ascending aMe12 branch (right) are typed as pale and those with a Tm5a branch (left) are typed as yellow. Columns with neither aMe12 nor Tm5a branches are also considered probable yellow columns (see below and <u>Methods</u>). When available, R7 photoreceptor connectivity in the column was also considered and some columns with conflicting features classified as unclear. **h** Scatter plot of R7 connections (combined for the left and right optic lobe) to the indicated neurons, with the color coding indicating the presence of Tm5a, aMe12, neither or both, by column. The scatter plot only includes columns with reconstructed R7 photoreceptors, the table inset shows counts for all columns (except in the DRA and at the medulla edge). **i** Scatter plot of R7 connections as in (h) with color-coding now indicating the different column types. Numbers included in the legend are counts for all columns (except DRA and edge). The general approach of classifying columns as pale and yellow followed Nern *et al.* (2025) but the now much more complete aMe12 reconstructions enabled a more complete typing of pale columns. Because we now expect most pale columns to be identified through aMe12 branches, we tentatively assign most of the remaining columns as yellow (yellow (-Tm5), see <u>Methods</u>.) **j** Medulla column maps showing left and right optic lobe column patterns with 6 column identities indicated: pale, two types of yellow columns (generally identified by the presence or absence of a vertical Tm5a branch), DRA (dorsal rim), edge columns which lack R7/R8s, and the few remaining columns indicated as ‘unclear.’

**Figure S5.**
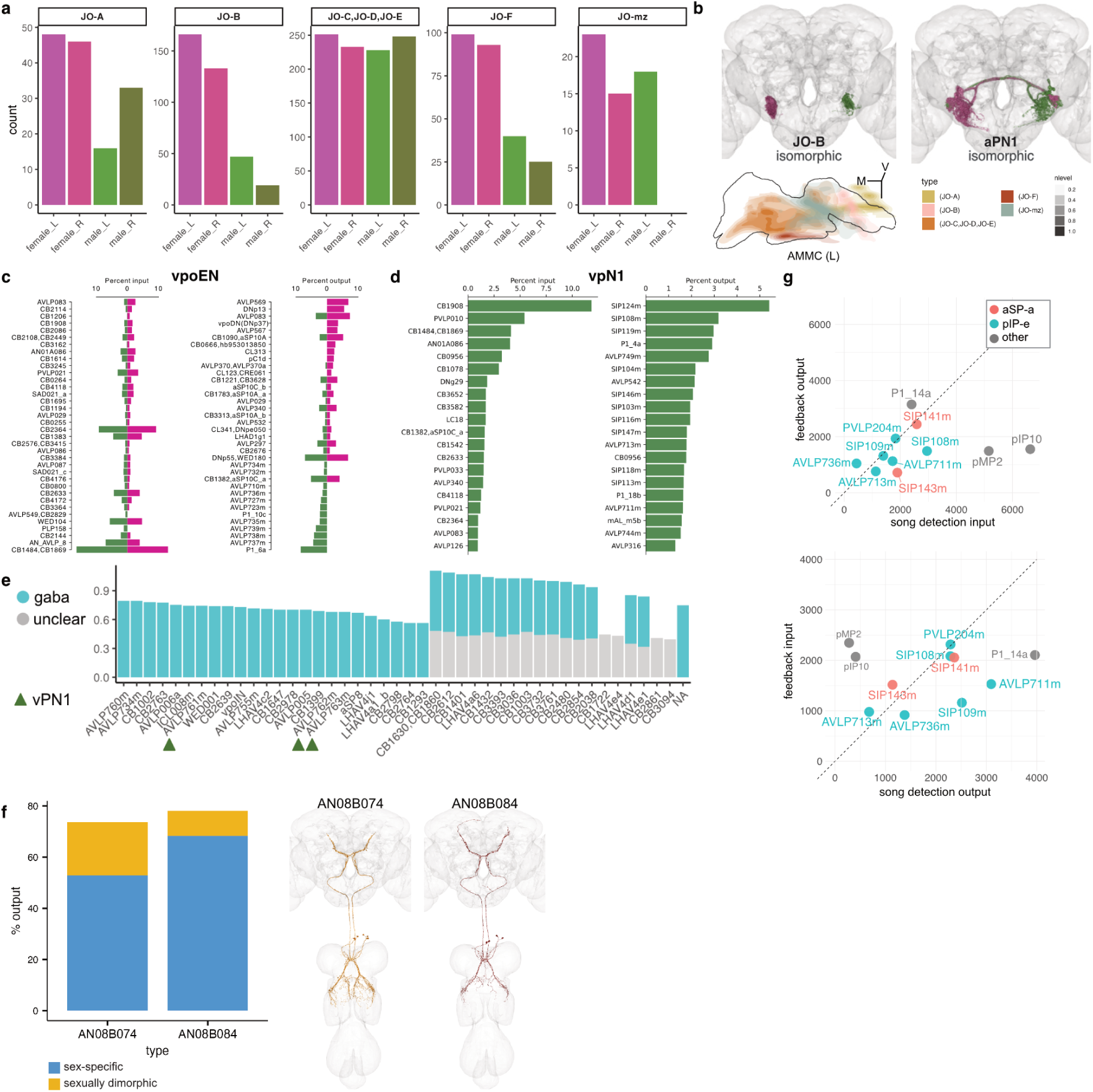
Additional auditory system analysis, related to figure 5. **a** Johnston’s Organ Neuron count by type in males (male CNS) and females (FAFB/FlyWire). Despite segmentation issues, we were able to identify around 20% of JO-B neurons. **b** Top left: JO-B neurons in female (FAFB/FlyWire, magenta) and male (male CNS, green) brains. Bottom left: Kernel distribution plots of AMMC innervation by JO neuron types in male CNS. Right: Second-order auditory neurons, aPN1, in female (FAFB/FlyWire, magenta) and male (male CNS, green) brains. **c** Top input (left) and output (right) partners of vpoEN in males (green) and females (magenta). **d** Same as in c for the male-specific vPN1 neurons (AVLP761m, AVLP762m, AVLP763m). **e** Average confidence score for consensus neurotransmitter predictions in the LHl1 lineage in male CNS; vPN1 cell types indicated by green arrowheads. **f** Left: Fraction of sexually dimorphic (gold) and sex specific (blue) outputs for AN08B074 and AN08B084. Right: AN08B074 and AN08B084 in male CNS EM space. **g** Scatter plot of the synaptic connection strength with song detection (x-axis) and AN feedback (y-axis) for cell types comprising *fruitless*-expressing clones in the circuit diagrams in Fig 5b, f.

**Figure S6.**
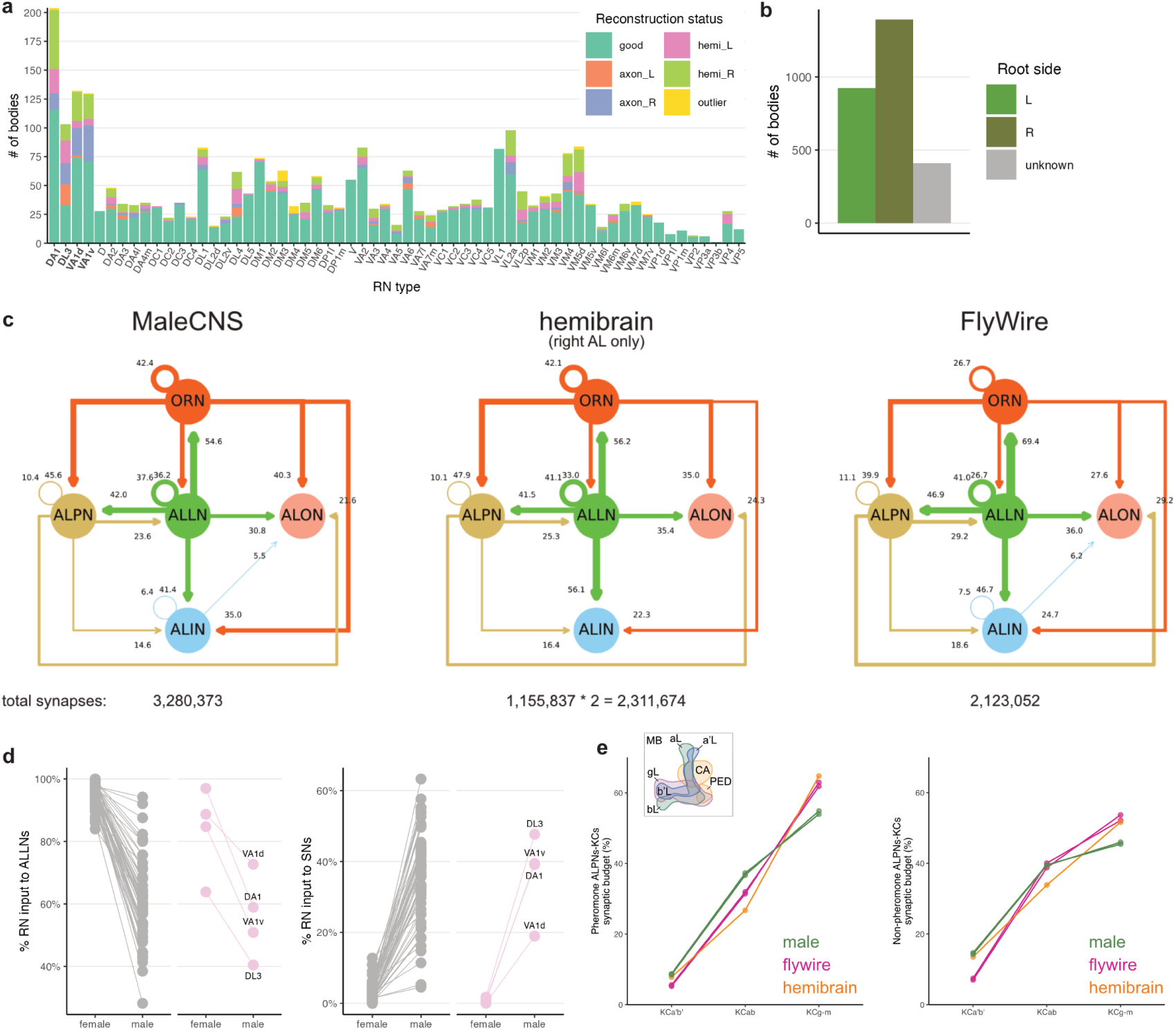
Proofreading and connectivity analysis in the olfactory system, related to figure 6. **a** Number of bodies per RN type and its reconstruction status. axon_L/R: axon (presynapses) only present on that side, though the body crosses the midline. hemi_R/L: body ends around the midline, and only exists on one side. Outlier: cases in which a body innervates more than one glomerulus on one side or does not arborise a glomerulus proper. **b** Number of bodies per RN type and per root side. **c** Input-normalized wiring diagram between AL cell classes for AL-intrinsic connectivity in males (left) and females (FAFB/FlyWire, right). **d** Comparison of the percentage input to RNs from ALLNs (left) and sensory neurons (right), for the female (FAFB/FlyWire) and male brains. ALLNs and sensory neurons are the top 2 classes that input to RNs. **e** Comparison of the synaptic budget, as percentage for the connections from ALPNs for the main types of Kenyon cells in the male and female brains; the differences observed between datasets for the KC classes are possibly due to biological variability. Left: for pheromone ALPNs; Right: for non-pheromone ALPNs. Data for each side is shown separately for FAFB/FlyWire and the male CNS brain.

**Figure S7.**
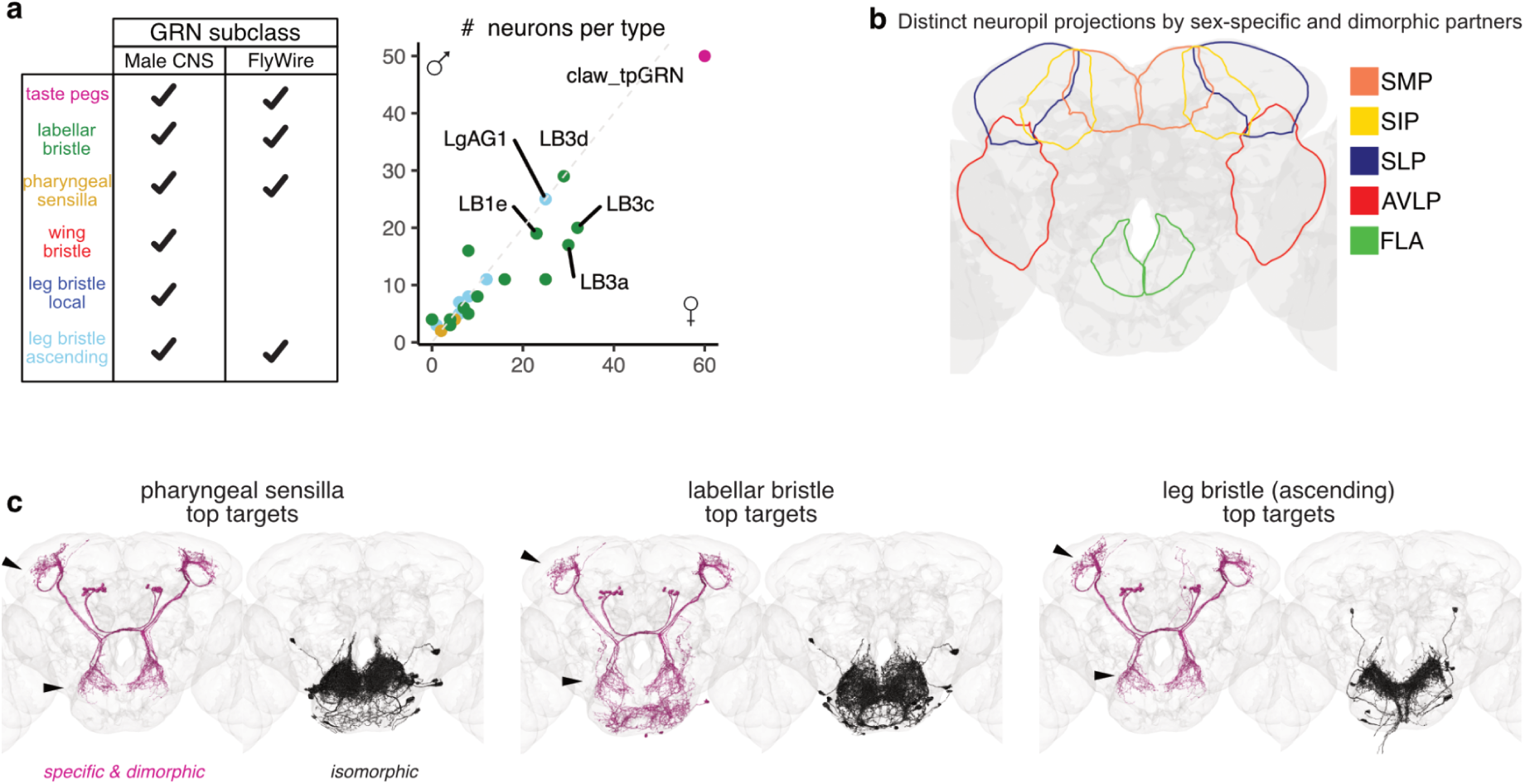
Additional gustatory system analysis, related to figure 7. **a** Left: Table comparing male CNS subclasses of taste sensory neurons with FAFB/FlyWire data. Wing bristles and local leg bristles are exclusive to the VNC. Right: Comparison of GRN cell type counts across males and females (pink: taste pegs; green: labellar bristles; yellow: pharyngeal sensilla; red: wing bristle; navy: local leg bristles; cyan: ascending leg bristles). Note that differences in labellar bristle count appear to be technical in origin rather than true sex differences^137^. **b** Strongest neuropils innervated by second-order sex-specific and sexually dimorphic gustatory neurons. **c** Top ten targets downstream of GRN types in female (FAFB/FlyWire), by type.

**Figure S8.**
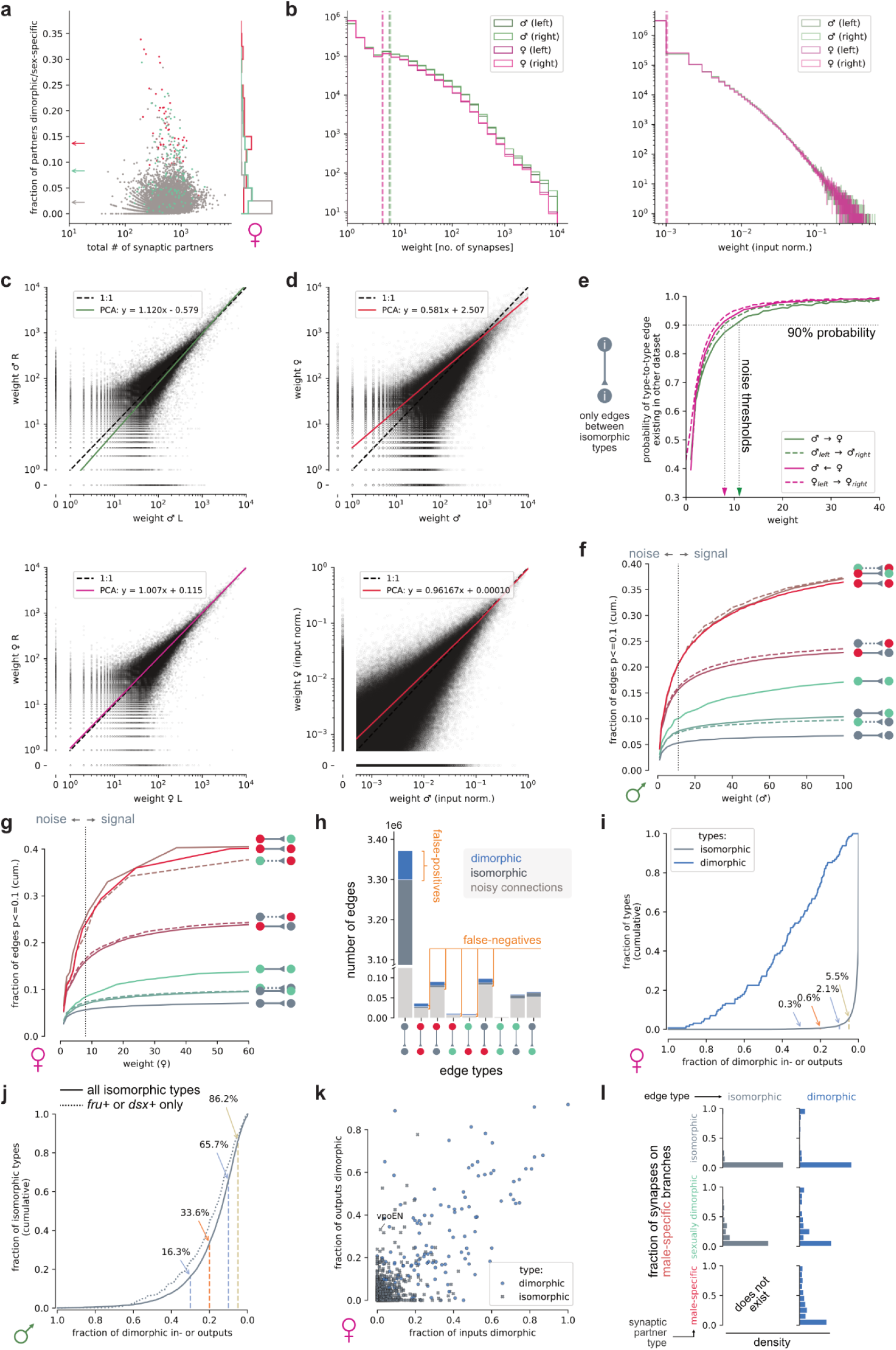
Supporting dimorphism analyses, related to figure 8. **a** Dimorphic up- and downstream synaptic partners as a fraction of total synaptic partners for all cross-matched female cell types. Arrows on the y-axis indicate means for iso-, dimorphic and male-specific types. **b** Histogram of total (left) and input-normalised (right) cross-matched type-to-type edge weights for both male and female. Dotted vertical lines indicate the means. **c** Left versus right weights for cross-matched type-to-type edges in male (top) and female (bottom). Green/magenta line represents PCA fit. **d** Male versus female edge weights for cross-matched types. Top: total number of synapses; bottom: input normalised. Red lines represent PCA fit. **e** Probability of finding an edge between isomorphic neurons of a given weight in another hemisphere of the same or different dataset. **f** Cumulative fraction of male edges with FDR-corrected p-values below 0.1 per edge type. Dotted vertical line marks 90% probability threshold from e. **g** Same as f but in female. **h** Total number of dimorphic, isomorphic and noisy connections (male + female) broken down by edge type. **i** Fraction of iso- and dimorphic types with at least X% dimorphic in- or outputs (by synapse count) in female. **j** Fraction of isomorphic types in male with at least X% dimorphic in- or outputs (by synapse count). Same analysis as Fig 8j but using only the iso-iso edges in the graph and without removing presumed false-positive dimorphic edges. Shows that if at least one partner is fru+/dsx+, there is a slightly higher chance of edges being flagged as dimorphic. While consistent with the idea that a minority of these connections are truly dimorphic, additional connectomes would be required to make precise conclusions. **k** Fraction of dimorphic in-versus outputs (by synapse count) per cell type in the female. **l** For each edge from/to a cell type that was split into male-specific and isomorphic branches, the fraction of synapses found on male-specific branches broken down by dimorphism of the synaptic partner (rows) and edge columns).

**Figure S9.**
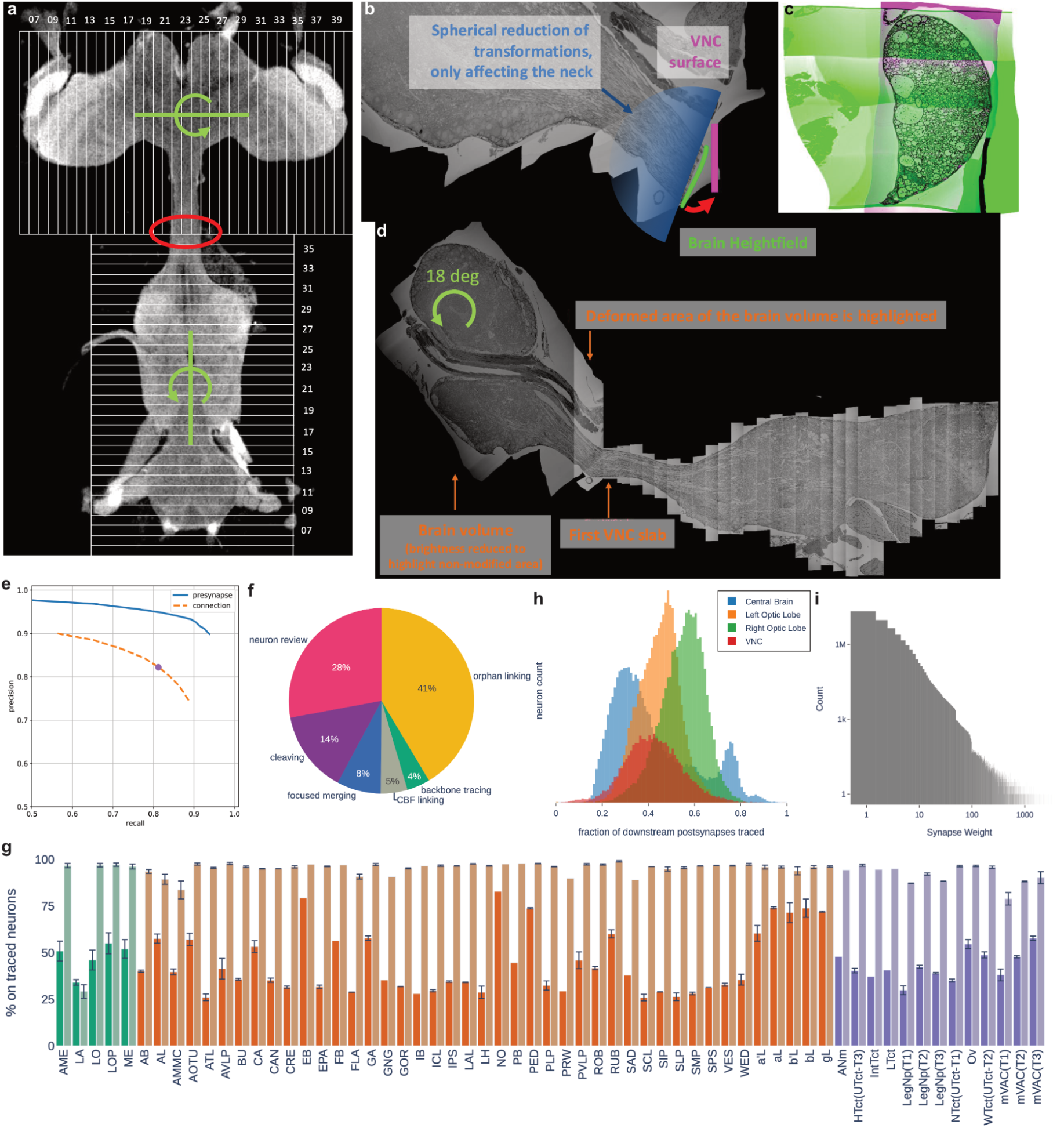
Dataset alignment and proofreading metrics, related to figure 1 and Methods. **a** Overview of the entire CNS and the slicing into slabs for FIB-SEM imaging. The red ellipse highlights the brain/VNC cut, green rotation axes show the degrees of freedom wrt to slab orientation. **b** Transformations that were applied to the brain and VNC volumes to stitch them seamlessly. **c** Overlay of the extracted surfaces of the brain (green) and VNC (magenta). Note the 4 slabs that are visible in the brain surface. **d** Cut through the entire CNS volume highlighting the connection between brain and CNS, the modified area of the brain for stitching both volumes, and the slabs of the VNC. **e** The precision-recall curves for presynapses and connections, sweeping across the model’s output confidence score. Our published synapses are filtered with a confidence threshold of 0.5 (purple dot), resulting in overall precision of 0.82 and recall of 0.81. **f** Estimated proportion of proofreading labor spent in each major proofreading protocol. **g** Traced completeness of synapses across neuropils. Light bars show presynapse completeness, dark bars show postsynapse completeness, and hue differentiates between optic lobe, central brain, and VNC neuropils. For paired left and right neuropils, the whiskers indicate the spread between left and right values and the bar shows the mean. **h** The distributions of downstream capture fraction for traced neurons in the central brain, optic lobes, and ventral nerve cord. **i** The distribution of synaptic partner counts of remaining unmerged ‘orphan’ fragments.

## Methods

### Data availability

A static download of the primary data artefacts for the male CNS connectome is available in the google bucket `gs://flyem-male-cns`. A landing page with ways to interactively explore the data can be found at https://male-cns.janelia.org. Derived data products specific to this paper are available at https://github.com/flyconnectome/2025malecns. Updated annotations for FlyWire are available at https://github.com/flyconnectome/flywire_annotations.

### EM Sample Preparation

Methods for sample preparation were described in detail in Nern *et al.*^36^, an earlier study of the same sample. Briefly, hundreds of five-day-old *Drosophila melanogaster* males from a cross between Canon S strain G1 × w^1118^ were dissected to extract the CNS intact, including the brain, VNC and neck connective. After a multi-stage fixation and staining process, each was inspected for significant damage, with special attention to the fragile neck connective. After initial inspection, 44 were examined in detail via X-ray CT imaging to check for more subtle flaws and to assess the staining. The sample identified internally as ‘Z0720-07m’ was selected as having the best quality.

### Hot-knife cutting

As the *Drosophila* CNS is too large to image by FIB-SEM as a contiguous block, we cut it into 20 µm-thick slabs via our “hot-knife” sectioning procedure^157^ (Fig S9a). The VNC was sectioned first into a total of 31 slabs using cuts along transverse planes. The brain was then sectioned into 35 slabs with sagittal cuts (orthogonal to those in the VNC). Quality of each slab was assessed by LM, followed by mounting, laser-trimming, and further inspection by X-ray CT.

### EM Volume Imaging

As explained in Nern *et al.*^36^, we followed the imaging methods first described in Takemura *et al.* ^34^ with minimal modifications. We employed a fleet of seven customized FIB-SEM machines to separately image each of the 66 hot-knife slabs over the course of a year. SEM images were acquired at isotropic 8nm in-plane resolution at 3 MHz using a 3 nA beam with 1.2 kV landing energy. FIB milling was performed with a nominal 8 nm step size using a 15 nA 30 kV Ga ion beam. The final 160 teravoxels (including fixative) occupied 0.082 mm^3^ total volume of which 0.054 mm^3^ contains tissue.

### EM Volume Alignment

As the brain and VNC volumes of our sample were imaged and hot-knife sectioned separately, we first assembled and stitched the hot-knife slabs of each volume independently, following the methods described in Nern *et al.*^36^.

Following a single cut through the neck that separated the sample into brain and VNC, the hot-knife sectioning of the brain and the VNC were performed orthogonally to each other (Fig S9a). Hot-knife sectioning fixes the two in-plane imaging axes, leaving the rotation around the milling axis as a free variable.

For convenient viewing and analysis, we chose to orient the brain volume such that all three of its anatomical axes approximately coincide with the volume coordinate axes. The faces of each hot-knife slab are sagittal sections, with the sagittal axis coinciding with the X axis of our aligned volume. We rotated the brain around the sagittal axis, tilting the head back by 18 degrees to align its dorsal-ventral axis with the Y axis of the volume and its anterior-posterior axis with the Z axis. In the VNC, the hot-knife cuts are transverse sections, with the Z axis reasonably aligned to the sample’s anterior-posterior axis.

We then proceeded to stitch together the brain and VNC volumes at their interface in the neck connective (Fig S9b-d). The cut through the neck spanned four hot-knife brain slabs of the brain volume on the anterior side, whereas it formed the first slab surface of the VNC volume on the posterior side. To extract the cut surface in the brain volume, we adjusted our hot-knife surface finding code and mostly relied on the BigDataViewer-based interactive tool to manually define the heightfield describing the exact location of the surface in the brain volume. An angle difference of ∼25 degrees remained between the surfaces. To match both volumes seamlessly, we developed code that applied a single non-rigid transformation consisting of the surface deformation field, the surface heightfield (including the ∼25 degree rotation), and a global translation. The software reduces the magnitude of the transformation as a function of distance from the stitching plane to ensure a smooth transition that is limited to the neck region, since proofreading had already started in the brain volume. After interactively confirming the correctness of this complex transformation on the two large VNC and brain volumes in BigDataViewer, we joined both volumes into a single, seamless volume of the entire CNS.

### EM Volume Segmentation

Segmentation for neurons and semantic masks (nuclei, glia, etc.) was performed with the same methods as the MANC VNC volume^34^, with some differences for this sample as described in Nern *et al.*^36^. Because the complete CNS volume was imaged and aligned as three separate subvolumes to be stitched together (right hemisphere, left hemisphere, and nerve cord), the segmentation procedure was run three times. For the nerve cord portion, a separate semantic segmentation model was trained to properly differentiate between muscle tissue and the nerve cord. Stitching the main segmentation for the left hemisphere onto the right hemisphere (and the VNC onto the brain) required care, as the earlier portion had already undergone significant manual proofreading by then. The segmentation region for each subsequent subvolume was constructed to overlap with that of the preceding subvolume. In the unified agglomeration, merges across the stitching boundary were selected according to the highest count of shared voxels among agglomerated segments within the overlapping region. Separation constraints to forbid merges between nuclei and fibers in the same nerve bundle (or the cervical fiber bundle) helped to avoid introducing false merges in already proofread segments.

### EM Volume Synapse Identification

We performed synapse prediction by extending the methods as described in the optic lobe reconstruction^36^. Two separate networks for presynaptic T-bar detection and postsynaptic partner detection from that reconstruction were further fine-tuned with additional training ground-truth synapse data collected in the brain and VNC regions of the full CNS sample.

As with the previous reconstruction, we collected separate validation ground-truth to assess the performance of the synapse identification. In total, synapses were densely labeled in 114 cubes of 300 × 300 × 300 voxels, spanning 81 ROIs. Cube locations were selected to cover distinct ROIs, and randomly selected within a given ROI. Altogether, the subvolumes contained 2,303 annotated T-bars and 16,870 annotated postsynaptic partners.

By selecting validation cubes that span many ROIs, we sought to better assess global synapse prediction accuracy, as well as identify any potential problematic ROIs. In our case, initially selected validation cubes in the lamina (LA) showed poor recall in the set of synapse predictions. This recall issue resulted from some presynapses in the lamina having a differing morphology, which was not well-represented in the original training set. We subsequently used these annotated lamina cubes to fine-tune a new T-bar detector, which we applied to the lamina ROIs, adding any predictions made by the new detector that did not already exist in our original set. (New validation cubes were then sampled and annotated for the lamina.)

Fig S9e gives the overall precision-recall plots for T-bars alone and synapses as units (both components correctly predicted); performance is comparable to the optic lobe subset. Fig S1i gives the precision/recall for each ROI, for T-bars alone, and for synapses. Source data for the plots are provided in the accompanying github.com/flyconnectome/2025malecns repository. Circle area is proportional to ROI volume, and color indicates ROI location, with red used for brain ROIs and blue for VNC ROIs. There is some variation in accuracy across ROIs, but performance is largely clustered, and generally above 0.8 recall for T-bars alone and 0.7 recall for synapses.

### Neurotransmitter prediction

#### Ground Truth

The ground truth for neurotransmitter prediction is made up of data from the literature, as well as experiments run at Janelia, and is available as a regularly updated resource [on Github](https://github.com/funkelab/drosophila_neurotransmitters). This resource maps cell types to the emission of ten neurotransmitters: acetylcholine, dopamine, GABA, glutamate, glycine, histamine, nitric oxide, octopamine, serotonin, tyramine. The data is kept in ternary form for each neurotransmitter, a value of `-1` signifying an absence, a value of `+1` signifying a presence, and a value of `0` signifying that the neurotransmitter was not tested for. The specific subset of cell types used to train our model can be found in the gt_sources directory of that repo.

The cell types from the resource were matched to male CNS cell types according to the `flywireType` and `mancType` neuprint columns, taking synonyms into account from FAFB/FlyWire. We then filtered the ground truth to exclude entries with an evidence confidence rating below 3, removed instances of co-transmission (multiple `+1` values when pooled across studies of the same cell type) and dropped any cell types for which positive and negative evidence exists for the same neurotransmitter. Disjoint training and validation datasets of cells were created (80% and 20% of the neurons, respectively), stratified so that they held a similar distribution of neurons from each class. The automatically detected T-bar locations for each cell were listed, and we verified that the split (80/20) was still valid at the synapse-level.

#### Model training

We trained an image classifier network following Eckstein *et al.*^40^ to predict neurotransmitter identity of a pre-synapse from a region of interest centered around a T-Bar, and spanning 640nm^3^. The network outputs a normalized score for each of the 7 neurotransmitters considered: the scores add up to 1, and are akin to a probability. The highest number is chosen as the predicted neurotransmitter.

The classifier network is a ResNet50^158^ model. It was trained using stochastic gradient descent on 100,000 mini-batches of 32 volumes each. All input volumes were intensity-normalized to the range [−1,1]. The data was randomly augmented during training with random axis flipping; affine transformations with rotations up to ±π radians, shear up to ±0.1, translations up to ±10 voxels, and scaling within ±10%; intensity scaling by ±20% with fixed mean; intensity shifts up to ±0.2; Gaussian noise with mean 0 and standard deviation 0.1; and Gaussian smoothing with sigmas randomly sampled between 0.1 and 0.5 along each spatial axis. We used the **`AdamW`** optimizer with a learning rate of 1×10^−4^ and parameters β=(0.9,0.999), λ=0.01, where β corresponds to the momentum parameters and λ to the rate of weight decay. Code for training and inference is [available on Github](https://github.com/funkelab/synister_malecns).

#### Assignment of neurotransmitter predictions to neurons

Each segment was assigned an aggregate neurotransmitter prediction according to the scheme described in Nern *et al.*^36^. Accompanying the predictions we provide a confidence score for each segment’s assigned prediction according to the formula in Eckstein *et al.*^40^. Each presynapse in a segment is assigned a confusion score by selecting an entry from the synapse-level confusion matrix: the row is determined by the owning segment’s overall prediction and the column determined by the presynapse’s model prediction. The segment confidence score is defined as the mean of its presynapse confusion scores.

The database records the most frequent prediction among a neuron’s presynapses in the `predictedNt` property, except for those marked `unclear` as noted below. The most frequent presynapse prediction when pooled across all cells of a common type is recorded in `celltypePredictedNt`, except for those marked `unclear`. The confidence score accompanying `predictedNt` is given in `predictedNtConfidence`, and the confidence accompanying `celltypePredictedNt` is given in `celltypePredictedNtConfidence`.

Neurons or fragments with fewer than 50 presynaptic sites or those whose `predictedNtConfidence` is below 0.5 are given `predictedNt` of `unclear`. Similarly, cell types with fewer than 100 presynapses when pooled across all corresponding neurons or cell types whose `celltypePredictedNtConfidence` is below 0.5 are given a `celltypePredictedNt` of `unclear`.

Finally, the `consensusNt` property is a copy of `celltypePredictedNt`, except in cases where experimental ground truth for the cell type is available to override the model prediction. Additionally, all octopamine and serotonin results are set to unclear in `consensusNt`, owing to the relatively scant validation data used for those neurotransmitters. The `consensusNt` is the recommended property to use in most analyses.

### EM Volume Proofreading

We carried out intensive manual and semi-automated proofreading, quality control, annotation and analysis targeting all neurons in the image volume to produce a connectome that can be considered finished and fully annotated^30–34^ (as compared with draft connectomes that are automatically segmented and partially proofread^35,159^ or partially traced^42,146^) by the emerging standards of the field.

#### Major phases of proofreading

Current state-of-the-art automated neuropil segmentation methods leave many unresolved errors requiring manual proofreading to identify and correct. The segmentation was proofread over 3 years by 29 expert proofreaders (estimated effort of 44 person-years). We executed a combination of bottom-up and top-down proofreading protocols using a software toolchain for connectomic reconstruction including NeuTu^160^, Neu3^161^, and DVID^162^ (Fig S9f). Further descriptions and rationale are provided in Nern *et al.*^36^. As the image volume was assembled in three major phases (right hemisphere, left hemisphere, and nerve cord), we repeated all major protocols in our proofreading process for each subvolume as it became available.

The first phase was cell-body fiber linking, in which we attached orphaned cell bodies (somata) to their main axons. Next, we followed our cleaving protocol in order to eliminate major false merges in all segments deemed significant according to their synaptic counts or other qualities, such as passing through the cervical fiber bundle.

With the dataset largely free of major false merges, we could begin efficiently assembling fragments together into recognizable cell shapes. In the backbone tracing protocol, we targeted synapse-rich segments for coarse tracing until no obvious false splits remained on large branches.

To increase the fraction of synapses captured by our traced backbones, we executed two bottom-up protocols aimed at connecting medium or small fragments to their target neurons. The focused merging protocol provided automatically selected merge proposals for directly adjacent segments to proofreaders as binary decisions. The orphan linking protocol tasked proofreaders with tracing a fragment to a backbone segment (without knowing which target it is destined for). We attempted to trace every fragment with 100 or more synaptic connections throughout the dataset. In the central brain and right optic lobe we went further, examining fragments with at least 50 synaptic connections.

The final phase of proofreading is purely top-down: neurons are assessed holistically to spot overlooked false merges or false splits. In this reconstruction of an entire CNS, we often had the benefit of examining pairs or groups of homologous neurons in tandem to spot gross inconsistencies between cells of the same type. This protocol was executed primarily by proofreaders in the central brain and right optic lobe, whereas we relied on the cell typing process to flag errors in the left optic lobe and VNC.

In addition to our major proofreading protocols, expert neuroanatomists from both Cambridge and Janelia examined all neurons as they typed and annotated them, providing a final quality check as poorly reconstructed neurons cannot be easily clustered and typed.

Our proofreading team is staffed almost entirely by annotators with experience from earlier connectomic reconstructions. Newly hired proofreaders received at least six weeks of training using a separate database before transitioning to the production environment. Regular “refresher” training was given to all proofreaders when we transitioned between major phases of proofreading. Two senior proofreaders took on a specialized role to provide oversight and quality control, reviewing the work of other proofreaders in addition to performing their own proofreading tasks. Based on an analysis of our edit logs and task management records, we estimate that our proofreading team spent approximately 44 person-years on the CNS dataset.

We proofread all fragments from the initial segmentation with > 100 synaptic connections. 98.9% of the 141,780 detected neuron-associated nuclei are part of a proofread neuron. After proofreading, only 0.6% of neurons with attached soma are composed entirely of fragments below this threshold. Thus, for the 1.1% of nuclei which remain unassociated with proofread arbours, we estimate that > 99% of the corresponding neurons have been proofread but have not been connected to their cognate nucleus in the cell body rind on the outside of the brain. For sensory neurons (without a cell body in the dataset), we seeded and traced axon profiles in all nerves entering the volume.

#### AutoProof

As described in the previous section, significant effort is spent manually proofreading the automated segmentation, and this process constitutes the most labor intensive component in generating a connectome reconstruction. With the accuracy of current automated segmentation methods, proofreading the entire segmentation is intractable. Manual effort is instead spent to target a specific connectivity coverage (see previous section). This workflow leaves a certain amount of the segmentation unexamined; within the orphan linking protocol, disconnected orphan fragments are only examined if they are above a threshold for synaptic weight.

We developed a new procedure, AutoProof, to automatically process these unexamined fragments and select a subset that can be joined to target neurons with high confidence. A machine learning model is first trained on the manual proofreading decisions from the focused merge protocol, thereby learning, given a pair of adjacent segments, whether the segments belong to the same body or not. This model is then applied to all unexamined orphan fragments in the central brain and nerve cord with a synaptic weight of ten or higher. The full details of Autoproof are given in Huang *et al.*^163^.

A random sample of orphan-to-target merges proposed by AutoProof was manually verified, and used to select a high conservative threshold, targeting a low estimated error rate of ∼3%. All proposed merges above this threshold were automatically accepted without explicit human verification; the risk is in part mitigated as any given orphan is of small synaptic weight and therefore individual errors would have minimal impact on the overall connectivity.

For the male CNS reconstruction, AutoProof helped the completion rate significantly. About 200 thousand orphan link proposals were automatically accepted, adding about 2.4 million PSDs and 309 thousand T-bars. These synapses added about 1.3% to the connectivity completion rate. At an estimated 200 tasks per day for a human annotator working in the orphan link protocol, this automated procedure was equivalent to about 4 person-years of work.

#### Quality metrics

The quality of a connectome can be assessed using a variety of metrics. One important metric is the overall fractions of presynapses and postsynapses which have been captured by ‘traced’ neurons, known as presynaptic and postsynaptic ‘completeness’. As can be seen in Fig S9g, presynaptic completeness tends to be much higher than postsynaptic completeness, as presynapses typically reside on larger-calibre neurites which are more easily merged via both automated segmentation and manual proofreading. This trend is also observed in prior large-scale *Drosophila* connectomes^30,32,34^.

A synaptic connection is only useful to downstream analyses if both its presynapse and postsynapse belong to traced neurons, so a more informative metric is the connection completeness, i.e. the fraction of synaptic pre-to-post connections in which the neurons on both sides have been traced. Fig 1d shows the fraction of completed connections in each neuropil, with an overall average of 40.1%. Empirically, connection completeness is approximately equivalent to the product of presynaptic completeness and postsynaptic completeness.

However, not all neurons in a compartment are equally well traced. An informative neuron-centric metric is the ‘downstream capture’ of each neuron, which is defined as the fraction of synaptic outputs which connect to traced neurons. Even when a neuron is itself completely reconstructed, if its postsynaptic partners are poorly reconstructed then its connectivity to downstream neurons will be poorly characterized. The discrepancy between compartment-level completeness and neuron-level connection recall is revealed by the downstream capture metric. The distribution of downstream capture fractions among neurons in the major compartments of the CNS is shown in Fig S9h.

It is worth noting that the downstream capture distribution for the central brain is bimodal. The second peak is chiefly due to populations of neurons whose outputs are in areas with relatively high postsynaptic completeness: the EB, FB, and NO in the central complex; the antennal lobes; and all compartments of the mushroom bodies.

Another quality metric is the number of unmerged ‘orphan’ fragments in the dataset and their distribution of synapse counts. The largest of such orphans remain unmerged in our dataset due to data artifacts or especially complex morphologies which make it difficult or impossible to find the site at which they should be traced to a neuron. We inspected and attempted to eliminate all orphan fragments with 100 or more synaptic partners. There are 5,329 remaining orphan fragments at or above that threshold in the proofread dataset. In total, there are 84.6 million orphan fragments in the dataset, most of which contain very few synapses. The final distribution is shown in Fig S9i.

### Spatial Transforms

We generated spatial transforms to register the male CNS to other coordinate spaces, including the JRC2018M brain light microscopy template^76^ (procedure described in Nern *et al.*^36^), and the MANC, FAFB/FlyWire and hemibrain connectome datasets. In all cases, we began with an automated registration between synapse point clouds using either Elastix^164^ or CMTK (RRID:SCR_002234) to produce a non-linear transform. Where necessary, we fine-tuned the transform using BigWarp^165^ in ImageJ/Fiji by manually adjusting pairs of corresponding landmarks. In addition to these bridging transforms, we also generated a transform to mirror the male CNS across the midline which aligns homologous neurons from the left hemisphere to the right or *vice versa*. All transforms are accessible with *navis-flybrains* (see Table 2) and can be chained with existing transforms to map to additional coordinate spaces, such as JRC2018 unisex brain, the JRC2018 unisex VNC, etc using *navis*. Template-to-template transformation fields can be obtained from https://www.janelia.org/open-science/jrc-2018-brain-templates.

**Table 2:**
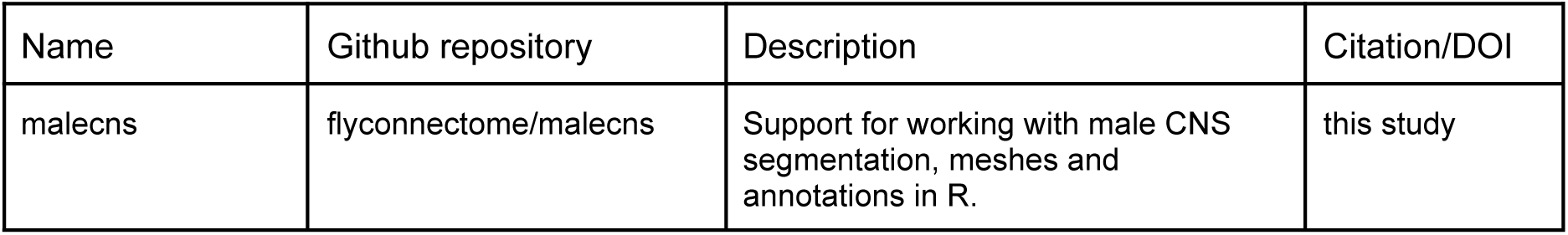
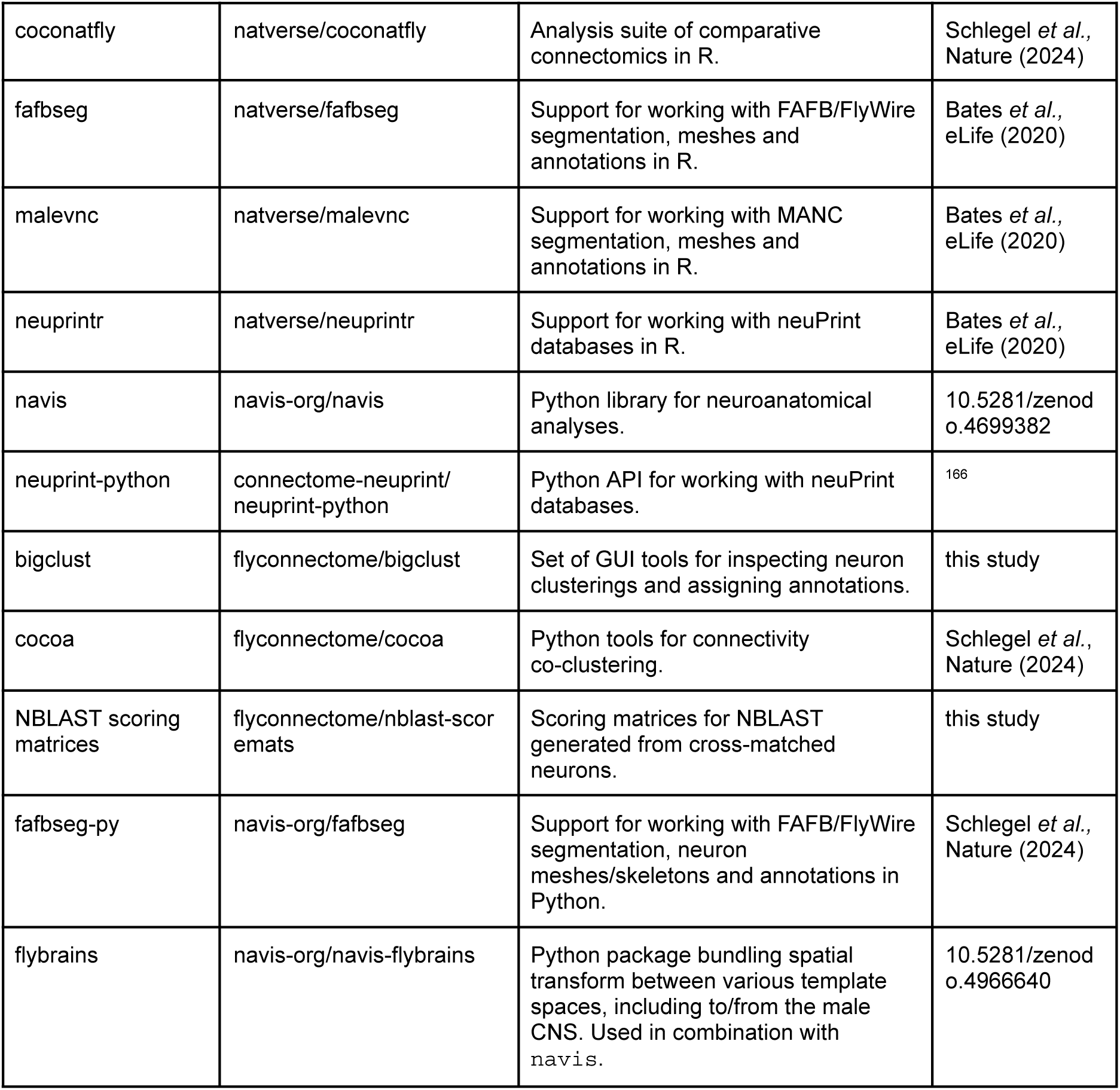
Software products used for analysis.

### Defining anatomical regions in the EM volume

As described in Nern *et al.*^36^, the primary neuropil regions were initialized via transfer from the JRC2018M template^76^ using the synapse point cloud to register the volume. We then refined the boundaries of those regions which did not align cleanly to the underlying EM data or relevant populations of segmented neurons.

### Analysis Software

Data curation and analysis was carried out with a combination of free and open source software tools for R and Python, principally from the *natverse* (https://natverse.org/) and *navis* (https://navis-org.github.io/navis/) ecosystems developed in Cambridge.

### Annotations

Each neuron in the dataset was given a set of annotations for various anatomical, developmental and descriptive features. We highlight some of the main ones below.

#### Superclass

Superclasses describe broad sets of neurons, grouped together by anatomy or general function. Bodies in the dataset are defined as neurons if they have a superclass. Bodies without one are fragments of neurons. We define 20 superclasses: intrinsic neurons are those contained entirely within the optic lobes, central brain or VNC; visual projection neurons connect the optic lobes and the central brain while visual centrifugal neurons do the opposite; sensory neurons extend their axons from the periphery into the CNS; sensory ascending or sensory descending neurons are sensory neurons that extend from the VNC into the brain or from the brain to the VNC, respectively; ascending neurons have their somas in the VNC and extend into the brain while descending neurons do the opposite;. motor neurons extend their axons out of the CNS to connect to muscle; efferent neurons are non-motor efferent that can be found entirely contained in the central brain or VNC or connecting these 2 regions (descending or ascending); finally endocrine neurons connect the central brain to the corpora allata or cardiaca endocrine glands.

Superclasses that are found across the CNS such as intrinsic and sensory, are given a prefix to distinguish between the populations in the optic lobe (ol-), central brain (cb-) and VNC (vnc-).

#### Hemilineage

Hemilineage annotations were transferred from the FAFB/FlyWire^30,31^, hemibrain^32^, and MANC^33,34^ datasets according to cross-matched neuron types. Unmatched types and many:1 matches were reviewed as described in Schlegel *et al.*^31^.

#### Supertype

Supertypes were assigned on a per hemilineage basis to morphologically similar neurons in the central brain using morphological (NBLAST) clustering^31,41^ and subsequent visual inspection. Neurons were grouped into a supertype if their main arbours aligned and when dendrites and axons were mostly overlapping in 3D space.

In some cases, as with the SLPal2 hemilineage, sex-specific terminal cell types can be linked together through their supertype designation. For example, supertype “12551” includes a group of sex-specific neurons previously described as the sexually dimorphic aSP-g^26,63^ or aSP6^59^ cluster. While these neurons differ widely in morphology and connectivity, preventing them from being linked at the level of terminal cell types, their classification under a common supertype reflects a shared development underpinning sex-specific functions within their respective neural circuits^20,26,167^.

#### Dimorphism

Dimorphism annotations were added to neurons on a per cell type basis according to a set of operational definitions used to describe three categories (1) sex isomorphic, (2) sexually dimorphic, and (3) sex-specific. Neurons from the FAFB/FlyWire, hemibrain (where possible), and male CNS datasets were clustered based on morphological (NBLAST) and connectivity (cosine similarity) metrics, and visualized with a dendrogram (Ward’s method) or a low-dimensional embedding (UMAP) (see also Fig S8a-d). Isomorphic neurons were matched across hemispheres and datasets, while sex-specific neurons were defined as having no probable match across opposite-sex datasets despite bilateral consistency within sex. Sexually dimorphic neurons were annotated when neurons could be closely associated enough such that they were clearly arising from the same cell type, but demonstrated numerical and/or morphological differences across sexes despite their shared developmental history. In some developmental lineages (e.g. FLAa3), high morphological similarity between types obscured our type-to-type matches. In many cases, types that fall into this category are considered “sex-specific”, but integration of future datasets may enable stronger type matching and these cells could turn out to instead be sexually dimorphic. Due to the lack of a fully annotated female VNC connectome, our dimorphism annotations are focused on the central brain and optic lobes. While we were able to add some additional dimorphism annotations for ascending neurons with extensive arbours in the brain, the majority of dimorphism annotations in the VNC come from existing literature^50,58^.

We offer a basic confidence assessment for our dimorphism annotations: dimorphic neurons are labeled either “sexually dimorphic” or “potentially sexually dimorphic”, and sex-specific neurons are labeled “male-specific” or “potentially male-specific”. We consider types with the qualifier “potentially” as lower-confidence annotations. However, analyses of dimorphism throughout the paper consider all dimorphism labels regardless of their confidence.

#### fruitless, doublesex and synonyms

Neurons in EM data were annotated as putatively expressing *fruitless* and/or *doublesex* according to comparison with light microscopy data transformed and projected into a registered template space^76,168^. Our analysis focuses on the central brain but we did annotate a small number of fruitless and doublesex-expressing neurons in the VNC based primarily on previous annotations from the MANC dataset^58,150^. The light level image data were obtained from Virtual Fly Brain^169^. Original data were sourced from Cachero *et al.*^63^ and Chiang *et al.*^71^ for *fruitless,* and Nojima *et al.*^27^ for *doublesex*. EM neurons were chosen for inspection against light data based on hemilineage assignment, since each light-level image was generated from individual lineages marked as ‘clones’^71,170^. Limitations due to lack of hemilineage assignment, ambiguity in lineage/cell type delineation, and technical limitations imposed by the genetic labeling prevented annotation for certain groups of cells. Specifically MARCM clone data are missing most primary neurons. The optic lobes where most neurons are have a different pattern of neurogenesis have not been well-characterised. Finally we excluded expression in mushroom body Kenyon cells where the subpopulation expressing *fruitless* cannot be clearly delineated.

EM neurons were compared to light images based on overlap in three-dimensional space and scored as ‘high’, ‘low’, or ‘NA’ indicating the amount of overlap with the clone and/or other evidence of gene expression according to previously published immunohistochemistry or transcriptome data, including Allen *et al.* 2025^152^. Hemilineages were putatively assigned to clones according to Costa *et al*.^171^. Annotations were further reviewed to ensure that neurons with the same cell type received the same annotation except in exceptional circumstances.

To validate the *fru/dsx* annotations, we compared our observed values to those expected from previous studies^27,63,64^. Considering only the brain lineages observed in Cachero *et al.*^63^, we identify 2,905 *fru*+ neurons in males and 2,068 *fru*+ neurons in females, compared to the expected 1,993.6(±75.7) and 1,668.4(±60.6), respectively. For *dsx*+ lineages reported in Nojima *et al.*^27^, we find 342 *dsx*+ cells in males and 134 in females, compared to 387.4(±18.6) and 66(±4.4) expected neurons (Fig S3j). For FAFB/FlyWire, our estimates are broadly consistent with a recent study in the same dataset^172^. Breaking this down per lineage shows that much of the discrepancy stems from lineages where prolific cell types have common morphologies, and this is consistent across sexes (Fig S3g). It is difficult to discern whether our labels are too inclusive, or whether methodological limitations in prior studies resulted in an underestimate. If the former is true, cell types with similar morphology may not always collectively express *fru* or *dsx*.

The synonyms column was filled for EM neurons which could be matched to previously published literature, especially when one or more names are used to describe the same cells. For putatively *fruitless*- and *doublesex*-expressing neurons, the name of the matching light-level clone^59,63,71^ were added to neurons with ‘high’ and ‘low’ confidence scores according to manual visual assessment.

#### Updated FlyWire annotations

In the process of adding annotations to the male CNS, we also added new (`fru_dsx` for *fruitless* and *doublesex* expression; `dimorphism` and `matching_notes` for dimorphic neurons; `supertype` matching the same field in neuPrint) and revised existing annotations for FAFB/FlyWire. The latter mostly affected `cell_type` and `hemibrain_type` but we also made minor adjustments to `super_class`, `ito_lee_hemilineage` and `side`. To reconcile cell types for optic lobe-intrinsic neurons, we also integrated the cell types from Matsliah *et al.* ^75^ with our original annotations. The `flywireType` field in neuPrint corresponds to these updated annotations (either `cell_type` or `hemibrain_type`) which have been deposited at https://github.com/flyconnectome/flywire_annotations.

### Typing and cross matching

#### Typing in the central brain and the ventral nerve cord

Neurons were principally typed by cross-matching to the FAFB/FlyWire^30,31,75^, the hemibrain^32^ and the male adult nerve cord (MANC)^33,34^. The initial typing was principally morphology-based (NBLAST) and generated a pool of cross-matched types for uniquely identifiable, often large neurons such as MBONs, uniglomerular olfactory projection neurons, or peptidergic neurons. This provided a scaffold on which to compute connectivity-based similarities between neurons from the different datasets. The connectivity-based approach was often required to make matches at the level of terminal cell types. In order to accelerate running a neuron matching pipeline, we developed R and Python tools to generate and efficiently review large-scale clusterings (see *coconatfly*, *cocoa* and *bigclust* in Table 2; sections below for details). Connectivity and skeletons (NBLAST) for the male CNS, MANC and the hemibrain were obtained from neuPrint. For FlyWire, we used the same L2-based skeletons as in Schlegel *et al.*^31^. For connectivity, we used the new FlyWire synapse predictions^156^ (materialization version 783) available from https://codex.flywire.ai.

The above matching pipeline was run iteratively: cell type labels were added/refined and then used to update the connectivity similarities. While the terminal cell typing was primarily connectivity-based, we frequently used NBLAST to cross-check labels. Matches to each of the other datasets are recorded in a set of fields in neuPrint: `hemibrainType`, `flywireType`, `mancType` and `mancGroup`. For cases when male CNS neurons match multiple types in the other dataset, the types are shown separated by a comma. The field `matchingNotes` includes, when necessary, relevant information for the cross-matching or typing. The primary `type` annotation represents the consensus type across all matched types. In a small number of cases, we modified existing types by defining a finer resolution subtype, merging two or more types, renaming the type or creating new types. For example, some of the tentative `CBXXXX` types introduced in FAFB/FlyWire were given new names following the hemibrain convention of `{main input region}{number}`. New male-specific types also follow this convention with an additional ‘`m`’ suffix, e.g. “`SIP136m`”.

The `group` field was used to record candidate matches early in the annotation process but retained later on as a more granular label than `type`. For the VNC, serially homologous neurons may have the same type but different groups in different neuromeres. In other cases, we think that differences between neurons with a different `group` but the same `type` may be the result of natural variation and not stereotypical differences observed across individuals. The field `instance` merges together `type` and `somaSide` or `rootSide` for sensory neurons.

#### Optic lobe typing

Cell typing for the left optic lobe largely followed the methods described in Nern *et al.*^36^, combining morphological similarity with connectivity. Extensive comparisons were made to the typed cells of the right side, greatly accelerating the process compared to the previous *de novo* cell typing effort. For Fig S4A and S4B we retain the cell type groups introduced in Nern *et al.*^36^: optic neuropil intrinsic neurons (ONINs), optic neuropil connecting neurons (ONCNs), visual projection neurons (VPNs) and visual centrifugal neurons (VCNs). We note that Fig 4 combines ONINs and ONCNs into one group OLINs. We are omitting the category of cell types grouped as “other” in Nern *et al.*^36^ in Fig S4A and S4B, as these cells are central brain neurons with minor optic lobe connectivity, and in many cases, while present in both hemispheres, do not have synapses in both optic lobes. The latter difference may be due to natural variability or reflect differences in precise boundaries of neuropil ROIs between hemispheres. The new, complete dataset provides a more complete description of the connectivity and brain region innervation of optic lobe associated cell types with synapses outside the (right) optic lobe. We found that three previously assigned^36^ optic lobe cell types were present in only one hemisphere (or, in one case, only had a marginal match between the candidate cell on each side) and appear to be also absent from other datasets. These cases may represent aberrant cells of unknown type rather than well-defined types.

#### Unmatched neurons

We cross-matched 96.4% of central brain and 98.8% of optic lobe neurons to FAFB/FlyWire and/or hemibrain datasets, and 93.1% of VNC neurons to MANC (Fig 1h). We were unable to match a fraction of efferent (15%) and sensory (15%) neurons in the VNC due to limitations in data quality and intrinsic variability which was not well-captured in cell typing in previous datasets. In addition, while ascending neurons could be confidently matched between male CNS and the previous MANC nerve cord connectome, around one third of these neurons could not be matched with the FAFB/FlyWire brain connectome since their arbours in the brain are too small for confident matching. Similar issues prevented confident matching of 29% of sensory ascending neurons that project from VNC to brain with the MANC dataset – reliable identification requires analysis of both brain and VNC

### Connectivity co-clustering

For terminal cell-typing and across-dataset matching of neurons we calculated a connectivity-based (cosine) similarity score. To efficiently run this pipeline on hundreds of thousands neurons we developed various R and Python tools: *coconatfly* and *cocoa*, respectively (see Table 2). For a given population of neurons – e.g. central brain neurons in FlyWire and male CNS, or VNC-intrinsic neurons in MANC and male CNS – we generated an observation vector where each row is a neuron and each column is an already cross-matched cell type. The values in that vector represent the number of synapses between a given neuron and the cross-matched cell type. We typically considered both the neurons’ in- and outputs to generate the vector, i.e. each cross-typed cell type has one column for connections to (inputs) and one column for connections from it (outputs). On a case-by-case basis would also occasionally just consider the in- or the outputs. Based on the connectivity vector we then computed pairwise cosine similarity, which is defined as the cosine of the angle between two vectors. Because this metric only considers direction and ignores magnitude, it automatically normalises for systematic differences in absolute edge weights between neurons from different datasets. The cosine similarities were converted to distances from which we calculated either linkages (using Ward’s method) for visualisation as dendrograms, or low-dimensional embeddings (UMAP) for visualisation as 2-dimensional scatter plots. See section below on the manual inspection of clustering results.

Connectivity data for the male CNS, MANC and the hemibrain were obtained from neuPrint. For FlyWire, we used the new FlyWire synapse predictions^156^ at materialization version 783 available for download from https://codex.flywire.ai.

### Morphological co-clustering

Morphological similarity scores were generated using NBLAST^41^ as previously described^31^. In brief, we obtained neuron skeletons from either neuPrint (for male CNS, hemibrain and MANC) or via the L2 cache (FlyWire) and pruned twigs smaller than 5 microns. Male CNS, hemibrain and MANC skeletons were additionally downsampled to average resolution of the L2 FlyWire skeletons (one node every 2.7um of cable). Depending on the NBLAST, we additionally either transformed skeletons into a common brain space and/or mirrored them onto the same side. From the processed skeletons we generated dotprop representations using the 5 nearest-neighbors (k=5) for each point to calculate the tangent vector. Dotprops were fed into the NBLAST function to compute both forward (neuron A → neuron B) and reverse (neuron B → neuron A) similarity scores. The final score was then calculated as the minimum between the forward and reverse scores. All relevant functions are implemented in *nat*, *neuprintr*, *fafbseg* (R) and *navis*, *neuprint-python*, *fafbseg-py*, *flybrains* (Python) (see Table 2). For inspection of the results, similarities were converted to distances from which we calculated either linkages (using Ward’s method) for visualisation as dendrograms, or low-dimensional embeddings (UMAP) for visualisation as 2-dimensional scatter plots. See section below on the manual inspection of the results.

### Inspection of clusterings

We developed various tools to efficiently inspect and integrate results of morphological or connectivity co-clusterings. For small-scale (10s to 100s of neurons) clusterings, we used functionality implemented in *coconatfly* (R) and *cocoa* (Python) to visualize similarities as either dendrograms and/or heatmap. For larger datasets, we developed *bigclust* (Table 2) which enables interactive exploration of clusterings of 100,000s of neurons as either dendrograms or low-dimensional embedding (UMAP) with linked visualisation of neuronal morphologies across datasets.

### Sensorimotor information flow analysis

Long-range flow of information from sensory inputs to motor neurons was assessed via maximum flow analysis of the synaptic graph. Unlike analyses of indirect connectivity^173,174^, graph traversal^68^, or dynamics of indirect connectivity traversal (influence)^147^ from only a source population of neurons, maximum flow can yield a distinct measure of information flow for each pairing of sensory source and motor sink populations. This analysis is also suited to long-range information flow, since maximum flow values do not necessarily exponentially attenuate like most multiplicative indirect connectivity measures. However, a limitation of maximum flow is that it does not identify alternate routes with equal or slightly less total flow value, so negative inferences about information flow based on low maximum flow values must be made with caution. Note that information flow does not consider the sign of interactions, i.e. neurotransmitter predictions were not considered in this analysis.

Maximum flow was computed for each combination of sensory source modality class annotation and motor domain subclass annotation. Efferents, endocrine neurons, and other nervous system outputs were not included. Input-normalized synaptic connectivity was used as edge capacities. Pseudo-source and -sink nodes were added to the graph with unit capacity connections to all source and from all sink neurons, respectively. All other inputs to sources and outputs from sinks were ignored. Thus the upper bound of each potential maximum flow was the minimum of the number of source and sink neurons. Maximum flow was computed using Dinic’s method^48^ on a sparse graph representation^175^, with floating point edge capacities in [0, 1] discretized into integral capacities in [0, 10000] and flows renormalized to [0, 1] after computation.

For each neuron, preference for a sensory modality was computed as the mean flow through the neuron across all maximal flows from that sensory source to all motor domain sinks. Likewise a preference score for motor domains for all neurons was computed as a mean over all flows from all sensory modality sources. Overall sensory or motor specificity for each neuron was computed as the maximum preference normalized by the sum preference for all sensory modality preferences and motor domain preferences, respectively. A corresponding combined sensorimotor specificity score was computed as the maximal flow value through a neuron normalized by the sum flow for all pairs of sensorimotor flows.

Due to the large number of abdominal motor neurons and their distance from most sensories, flows targeting them often spread broadly over neck connective neurons and caused these neurons to have relatively high preference values for the abdominal domain. To aid motor domain preference analysis for this population, a separate maxflow analysis was performed with the same sensory and motor annotation categories, except with abdominal motor neurons partitioned into separate domains for each annotated neuromere. This flow was not used for any other purposes; all other mentions of the sensorimotor flow analysis refer to the version with a single abdominal motor domain.

Analysis and visualization of sequential relationships in flows was performed using a pseudo-layering order, where neurons are assigned a layer value preferring to be one greater than the layer value of their flow inputs, weighted by the input-normalized flow value. Since flow is a directed acyclic graph, these layer assignments for all neurons, *l*, are computed using the (pseudo-)inverse of the Laplacian of the input-normalized flow adjacency matrix, *F*:

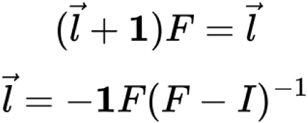

The data for all sensorimotor flows are provided in the accompanying github.com/flyconnectome/2025malecns repository.

### Sensorimotor clusters

For analysis of sensorimotor flow structure, descending and ascending neurons were clustered on the basis of a UMAP embedding of their sensorimotor flow. For clustering, the embedding was generated with 20 neighbors and a minimum distance of 0, using Euclidean distance. Clustering was performed on the mean sensorimotor flow aggregated by neuron type, projected into this embedding. Clustering was performed by simple k-means, with the number of clusters selected from a parameter sweep between [2, 60] maximizing the silhouette score. Cluster labels were reordered by clustering them via average agglomerative linkage of the cosine distance of their concatenated incoming and outgoing cluster connectivity.

Behavioural category compatibility of clusters was computed by multiplying the mean sensorimotor flow matrix of each cluster with a compatibility matrix for each behavioral category. Because the purpose was to demonstrate ease of interpretation of sensorimotor flow as it relates to putative function and behavioral hypotheses, rather than produce an optimal predictive algorithm, the compatibility matrices were manually constructed using simple rules of which sensorimotor pairings are likely predictive of compatibility with which behaviours. For those that are, those entries in the compatibility matrix are 1, while all others are 0. These pairing rules are as follows: for VNC sensation, any flow originating from gustatory, tactile, or proprioceptive modalities; for escape, any flow originating from vision targeting the legs or wings; for feeding, any flow originating from gustation or olfaction not targeting the abdominal domain, and any flow targeting the proboscis; for flight, any flow targeting the wings or halteres; for grooming, any flow originating from mechanosensory or tactile modalities targeting proboscis, neck, or front leg domains; for reproduction, any flow targeting the abdominal domain; for walking, any flow originating from any mechanosensory modality targeting leg domains. Both the compatibility and flow matrices are L_1_ normalized before multiplication.

The data for all sensorimotor clusters are provided in the accompanying github.com/flyconnectome/2025malecns repository.

### Axon-dendrite splits

Neurons were split into axons and dendrites (Fig 2) using the synapse flow centrality algorithm from Schneider-Mizell *et al.*^176^ as implemented in *navis* (see Table 2). In brief, for a given neuron we associate its in- (postsynapses) and outputs (presynapses) with the neuron’s skeleton. For each input, we draw a path to each output and count the number of paths going through each segment of the skeleton. The segment(s) with the highest “flow” typically represent the link between axon and dendrite. Removal of the linker results then splits the neuron into axon and dendrites. Splits for neurons shown in Fig 2 were manually confirmed. Based on the assignment of individual pre- and postsynapses to either axon or dendrites, we then re-calculated the neuron-to-neuron and from there the type-to-type connectivity split into axo-dendritic, axo-axonic and dendro-axonic connections.

### Effective connectivity

In Fig 4 (Visual system) we used the Connectome Interpreter tool^177^ to compute and visualise measures of effective connectivity (which can be computed for both direct and multi hop pathways). We first found direct and two-hop paths from all visual projection neurons (VPN) to either descending neurons (DN) that are sexually-dimorphic or -specific, or descending neurons for which there is existing research on the function of the cell type. We next filter the paths for both male and female, such that all direct connections are stronger than 1% of the input of the postsynaptic cell type. We further select from the filtered paths, ones that are unique to one sex, by removing paths that exist in the brain of the other sex, regardless of connection weight. We finally select only the paths that contain at least one sexually-dimorphic or -specific cell type.

We calculate the effective connectivity based on the selected paths: if the VPN connects directly to the DN, we use the connection strength. If the connection is through an intermediate neuron (i.e. two-hop), we use the square-root of the product of the connection weights.

For Fig S4A, panel h, we extract all the direct and two-hop connections from visual projection neurons in Fig 4I, to descending neurons with known functions or sexual dimorphism. We then filter the pathways such that the direct connections are stronger than 1% normalized input of the postsynaptic cell type.

### Edge normalization

Connections are either input- or output-normalized as defined in the text and figure legends.

**Input normalization** quantifies the relative contribution of each presynaptic partner to a postsynaptic neuron’s total input. For a connection from neuron A to neuron B, input normalization is calculated as:

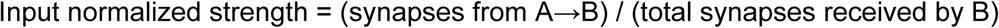

**Output normalization** quantifies how a presynaptic neuron distributes its synaptic output across its postsynaptic targets. For the connection from neuron A to neuron B, output normalization is calculated as:

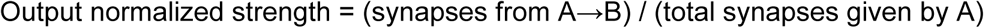

Input normalization reveals the relative influence of individual presynaptic partners on their downstream targets, while output normalization reveals the specificity and distribution of synaptic outputs.

### Definition of dimorphic edges

To define edges as either dimorphic or isomorphic (Fig 8), we first constructed a type-to-type graph for all cross-matched neurons with arbours in the central brain: superclasses `ascending_neuron`, `cb_intrinsic`, `cb_motor`, `cb_sensory`, `descending_neuron`, `endocrine`, `visual_projection` and `visual_centrigufal` for the male CNS and corresponding superclasses for FAFB/FlyWire. Connections made in the ventral nerve cord (VNC) of the male CNS were excluded. For FlyWire, we used the new synapse predictions^156^ for materialization version 783 (available from Codex).

Connection weights (i.e. number of synapses) were divided up between the left and right hemisphere using the source neuron’s soma (`somaSide`) as point of reference. For the small number of unpaired medial neurons that belong to neither side of the brain, we used the target’s side if possible. For connections between unpaired medial neurons (i.e. where neither source nor target belong to the left or right hemisphere), we split the weight evenly between left and right. For sensory neurons, we used the side of the nerve through which their axons enter the nervous system. To align connection weights between the male and female dataset, we modeled the systematic differences between the male and female dataset using a simple principal component analysis (PCA). This yields a scaling factor of 0.581 which we apply to all male CNS edge weights for cross-dataset comparisons (i.e. there are more synapses in male CNS; see Fig S8d).

This produced a table like this:

|  |  | weight in male (scaled) |  |  | weight in female |  |  |
| --- | --- | --- | --- | --- | --- | --- | --- |
| type_pre | type_post | weight_total | weight_left | weight_right | weight_total | weight_left | weight_right |
| AVLP532 | CL110 | 10.45 | 5.81 | 4.64 | 20 | 12 | 8 |

To extract connections that are significantly different between male and female we used a standard t-statistic, i.e. the ratio of the difference in a number’s estimated value from its assumed value to its standard error. The t-statistic *t_AB_* for a connection between cell types A and B is defined as:

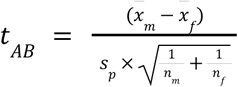

where *x̅_m_* and *x̅_f_* are the means between the number of synapses connecting A and B on the left and right hemisphere in male (*m*) and female (*f*), respectively.

The pooled standard deviation *s*_*p*_ is defined as:

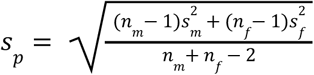

where *s_m_* and *s_f_* are the standard deviations over the number of synapses between A and B for the left and right hemispheres in male and female, respectively. The number of observations (*n_m_* and *n_f_*) is 2: one from the left and one from the right hemisphere, so this expression for *s_p_* simplifies to

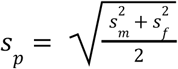

And the combined expression is therefore:

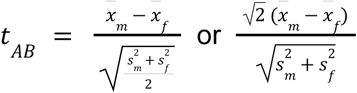

From the t-statistic, p-values were calculated using a two-sample t-test with 3 degrees of freedom as implemented in `scipy.stats.sf` ^175^. The resulting p-values were adjusted to control the false-discovery rate using the Benjamini-Hochberg procedure as implemented in `scipy.stats.false_discovery_control`.

We considered all edges with *p* ≤ 0. 1 to be potentially dimorphic. Across all 3.74M edges between cross-matched neurons in the central brain, we find 381k edges (10%) with significant (FDR-corrected p <= 0.1) male-female differences, 182k (4.9%) of which are highly significant (FDR-corrected p <= 0.01). This pool of candidate dimorphic edges includes a number of edges with small but very consistent male-female differences. We therefore additionally apply a threshold of at least 30% difference which reduces the number of candidate dimorphic edges to 378k. As demonstrated previously^31,178^, we find that weak isomorphic connections are unstable: around 60% of single-synapse connections in one hemisphere are not at all present in another hemisphere from either the same or another brain (Fig S8e). To achieve a 90% probability of an isomorphic edge in the male also being present in the female dataset, it needs to consist of >10 synapses. In the opposite direction, that threshold is >7 synapses – likely due to the generally stronger edges in the male CNS. We consider edges below these thresholds to be unreliable and treat them as noise. Because edge weight distribution is heavily skewed with many weak edges (Fig S8b), around 80% of all cross-matched edges fall below these thresholds. Crucially though, the remaining above-threshold edges collectively contain 90% of all synapses in the male (88% in female) (Fig S8f,g).

For subsequent analyses (Fig 8f and following), this initial set of edge-level dimorphism labels was corrected using the cell type-level dimorphism labels: false-negatives (edges involving sex-specific types on either end) were set to “dimorphic”, and likely false-positives (edges involving only isomorphic types) were set to “isomorphic”. The cross-matched type-to-type edges including dimorphism labels are provided in the accompanying github.com/flyconnectome/2025malecns repository.

### Graph traversal

We employed a probabilistic graph traversal model first published in Schlegel *et al*.^68^ to calculate the distance from sensory inputs for isomorphic, dimorphic and sex-specifc neurons (Fig 3l). Using a pool of all sensory neurons as seeds, the model pulls direct downstream partners and repeats the process until all nodes in the graph have been traversed. We used a linear function to determine the probability of traversal such that if neuron A makes up 30% or more of neuron B’s inputs, there is a 100% chance of traversal. The chance of traversal for any neuron outside the pool connected to a neuron already in the pool is independent. We repeated the traversal 10,000 times and calculated the layers as the mean across all runs. The results of this analysis are provided in the accompanying github.com/flyconnectome/2025malecns repository.

### Olfactory system analysis

The *Drosophila* olfactory system consists of approximately 1,300 olfactory receptor neurons (ORNs) in each hemisphere grouped into 53 types according to their receptor expression profiles^136,179–181^. ORNs expressing the same receptors project to the same antennal lobe compartment, or glomerulus, where they synapse with second-order projection neurons (ALPNs) that carry sensory information from particular glomeruli to higher brain centres. Antennal lobe local neurons (ALLNs) provide lateral inhibition and excitation^182^. In parallel, the AL also integrates thermo- (heat) and hygro- (humidity) sensation in distinct glomerular subcompartments in the AL (VP glomeruli, VP1-5) through their own receptor neurons (TRNs and HRNs, respectively). There are around 100 of these neurons grouped into 7-8 types^49^. We use “RNs” to collectively refer to olfactory, thermosensory, and hygrosensory receptor neurons, and “ORNs” when referring exclusively to olfactory neurons.

Sensory neurons are often difficult to reconstruct due to their fine calibre particularly as they cross the antennal commissure and artefacts likely associated with truncation of their axons during specimen preparation. Receptor neurons (RNs) in the male CNS were no exception with reconstruction status summarised in Fig S6a. Almost all RNs are bilateral (projecting to homologous glomeruli in left and right antennal lobes) with the side of entry into the neuropil annotated as `rootSide`. While the majority of RNs (∼70%) were fully reconstructed, some neurons could not be followed as they crossed over the midline leaving two separate fragments for which the side of origin was uncertain. When quantifying the number of RNs we use the term ‘neurons’ when we have accounted for the duplication of bodies where their side of origin is unknown, and ‘bodies’ when we have not. However, even accounting for this, there appeared to be asymmetries in both neuron numbers and synapse numbers between the two hemispheres (Fig S6b). For these reasons we opted to use the total number of RNs in our analysis rather than presenting an interhemispheric comparison (our approach in most other parts of the CNS). Furthermore, for most of our analyses, we were able to leverage the hemibrain dataset to provide an additional data point. However, this comparison was not satisfactory since ∼50% of glomeruli in that dataset are truncated and/or have fragmented RNs^68^.

In the male CNS we annotated 71 types of uniglomerular ALPNs (284 neurons) and 117 types of multiglomerular ALPNs (409 neurons) and 6 ALON types (14 output neurons with distinctive morphology compared with classic ALPNs). In the text, when we refer to ALPNs, this includes ALONs. To test whether ALPNs with high sex-specific or sexually dimorphic output (≥10% of presynapses; Fig 6g) differ in their odour input composition, we first assigned each ORN/uPN to one of five mutually exclusive valence categories (adapted from ref^68^). For bootstrapping, we resampled (with replacement) the remaining set of ALPNs 1000 times, and for each sample calculated the mean percentage input received from ORNs/uPNs in each valence category. This produced a bootstrap distribution of means per category, which we compared with the observed values for the high sex-specific/dimorphic output group.

### Network analyses

#### Hierarchical community detection

The hierarchical community structure is determined to analyse the contributions of isomorphic and dimorphic/sex-specific cell types in the network structure. Here, we fit a nested stochastic block model (SBM) to the data to infer partitions of nodes by their network connectivity. The SBM is a generative model for networks in which nodes are partitioned into blocks (or groups), and edges are placed between blocks according to block-specific connection probabilities. Given a particular partition, the likelihood that the observed network was generated under the model can be computed. To identify the best partition, this inference relies on the minimum description length (MDL) principle: the best model is the one that provides the most concise description of the data. In practice, this involves jointly encoding (i) the model parameters and assumptions (e.g., the block structure and probabilities) and (ii) the data given the model. The description length, measured in bits, is minimized when the model strikes the best trade-off between complexity (number of parameters) and goodness of fit (how well it explains the observed network). Importantly, this balance means that it does not overfit noisy data as is the case for modularity maximization based approaches^183^. The hierarchical partitions and the number of communities inferred are optimal by the above criteria. The procedure was employed with male cell-type connectivity graph of the central brain, where the synapse counts (edge weights) are modelled with a discrete geometric distribution, as determined to be optimal in the larval brain^184^. The graph-tool library was used to run the hierarchical block assignment^183^. We first obtained a greedy nested block model assignment and performed 1000 iterations of 10 Markov Chain Monte Carlo (MCMC) merge-split sweeps to improve the solution^185^.

#### Cluster enrichment analysis

Our hierarchical block structure provides a set of clusters which we evaluate for statistically significant enrichment in male-specific or dimorphic types. We used the hypergeometric distribution to assign each cluster a probability of randomly drawing at least the observed number of non-isomorphic types. We calculated this probability for each cluster across the whole hierarchy. Then, using the Benjamini-Hochberg method accounting for the false discovery rate (FDR), we designated clusters as enriched at α = 0.01. We obtained 13 clusters at the highest granularity level.

### Optic lobe analysis

#### Pale and yellow column identification

Medulla columns were classified as pale or yellow using both anatomical markers and synaptic connectivity, essentially following the previously described procedure^36^. Briefly, R7 photoreceptors were classified based on connectivity patterns with Dm8a/Tm5a (yellow) versus Dm8b/Tm5b (pale) cell pairs, and the presence of aMe12 and Tm5a branches in a column, used as anatomical markers. For most columns, column classification based on morphology and connectivity markers was consistent, the remaining cases were assigned based on the balance of evidence and sometimes left as ‘unclear’. As previously reported for the right optic lobe, we found some photoreceptors to be missing or incompletely reconstructed; the corresponding columns were assigned based on the anatomical markers alone. The main difference in the new dataset is that systematic additional proofreading of aMe12 cells, including in the right optic lobe, enabled us to identify many more of the aMe12s’ fine vertical branches, which serve as the most reliable known markers of pale-specific columns^186^, achieving substantial improvement on the prior effort (summarized in Extended Data Fig 5 of Nern *et al.*^36^). However, even with these improvements, the new analysis still revealed greater complexity than previously appreciated: the available anatomical markers are insufficient to place all columns (excluding edge and DRA columns) in two uniform groups, pale or yellow (as also discussed in the Methods of Nern *et al.*^36^). Making the assuming that with the new reconstructions the set of aMe12 branches is now close to complete, we implemented a refined classification scheme that identifies likely yellow columns (i.e. columns without aMe12 vertical branches) as two subtypes – those with and without Tm5a branches/connections. This updated column assignment yields column counts consistent with previous expectation^186–188^ while minimizing the number of ‘unclear’ unassigned columns (Fig S4Bf-j). Column type assignments for each medulla column are provided in the accompanying github.com/flyconnectome/2025malecns repository.

#### Neuron visualization

Three-dimensional neuron renderings for showing whole-brain projection views and optic lobe slice-views (Fig 4, Fig S4B) were generated using the Blender-based pipelines developed in Nern *et al.*^36^.

#### Comparison of edge weights in the optic lobes

Plots showing the percentage of inputs to the postsynaptic cells for pairs of connected cell types (for comparing males and females) or instances (for comparing the left and right optic lobe) in the optic lobes (Fig 4m, Fig S4B), include only connections with a combined synapse count above a threshold on at least one side of each comparison. Connection pairs that show high technical variability (e.g. those involving photoreceptors or cells in the lamina) or are entirely absent from one side of a comparison (i.e. sex-specific or present in only one optic lobe) are also omitted.

#### Eye maps for visual projection neuron spatial analysis

For the spatial coverage heatmaps of visual projection neurons in Fig 4 and Fig S4Ae, we first mapped the spatial locations of input synapses to column ROIs (described below). This spatial pattern together with the synapse count in each column were then visualized on a new eye map, representing the visual field of the compound eyes. This eye map was based on the data and methods of Zhao *et al.*^79^, but extended to accommodate the medulla column ROIs in the male CNS brain (detailed in Zhao *et al.*, ms in prep).

#### Cell Type Explorer interactive web resources

The Cell Type Explorer web resource was substantially updated to include bilateral optic lobe analyses and expanded to encompass connectivity summaries for all central brain neurons. The web interface now features enhanced interactivity, enabling users to filter and sort connectivity tables and browse cell types across the entire male CNS connectome. The right optic lobe web resource will remain at its original URL:

https://reiserlab.github.io/male-drosophila-visual-system-connectome/, while the updated version is hosted from https://reiserlab.github.io/celltype-explorer-drosophila-male-cns/.

#### Column and layer ROIs

##### Column and layer creation for the left optic lobe

Columns and layers for the left OL were created using similar methods to the right OL^36^, with one major difference: neurons were assigned to columns and hexagonal coordinates automatically rather than manually. We also updated several parameters used in column and layer creation and in the T4-to-Mi1 assignment for columns and layers in the LOP. We used these new parameters to also update the columns and layers in the right LOP.

##### Automated column assignment via community detection

To automatically assign columnar neurons in the ME to columns, we selected all 10626 neurons of cell types L1, L2, L5, Mi1, Mi4, Mi9, C3, Tm1, Tm2, Tm9, Tm20 and T1 in the left OL. For the right-side ME columns we had used these columnar cells in addition to C2, L3 and Tm4, but our previous analysis^36^ showed that these cell types are less columnar than the others (smaller fraction of their synapses could be assigned to single columns, with median < 0.65).

The key idea behind the automatic column assignment is that columnar neurons within a column should be more strongly connected to each other than to columnar neurons in other columns. Such an assignment can be achieved via community detection algorithms of networks. We applied generalized modularity maximization^189,190^, implemented with the Leiden algorithm (https://leidenalg.readthedocs.io/en/stable/intro.html). Modules (or communities) are non-overlapping sets of nodes that maximize a function called generalized modularity. Intuitively, this function compares the observed connectivity with that of an averaged random network that has matched in/out-degrees, i.e., the same total number of outgoing and incoming connections for each pair of neurons; but the latter connectivity is also scaled by a parameter γ. If the observed connectivity between a pair of neurons is larger than that of γ times the averaged random network, neurons are more strongly connected than expected and are therefore likely to be part of the same module; if below, they are less likely. Theoretical work has shown that γ controls the total number of modules found by generalized modularity maximization^190^ and that using a specific γ is equivalent to maximum likelihood estimation in certain stochastic block models with statistically similar communities^191^. In our application, we expect columns to form statistically similar communities, which makes generalized modularity maximization an appropriate choice of community detection algorithm.

The connectivity between the columnar neurons in the left ME determines the modules, and hence their column assignment. We removed weakly connected neurons (15 cells with ≤3 connections to any other columnar neuron or ≤10 total connections to other columnar neurons) and one clear merge error. We optimized the parameter γ by sweeping it over a wide range of values and observed how the number of modules changed. If robust modules exist, the number of modules should remain approximately constant over a range of γ’s and be close to the number of columns in the right ME (892 columns). We used the *leidenalg* Python library (https://leidenalg.readthedocs.io/en/stable/intro.html) and the functions “Optimiser”, “RBConfigurationVertexPartition”, “optimize_partition”, and “resolution_profile” with “resolution_range=(50,150)”. We found that the number of modules changed more slowly around 880 modules. We picked γ=100 which gave 884 modules—the same as the number of L1 neurons in the left OL. (We note that 892 also equals the number of L1 neurons in the right OL.)

Since we will use these modules to divide the ME ROI into column ROIs (see below), we further required that all neurons must have synapses in the ME ROI; due to imperfections of the ME ROI, this removed 7 more neurons, leaving 10603 neurons. Of the 884 modules, 790 modules contained exactly one neuron per cell type. 58 modules contained less than 12 columnar cell types but up to 12 neurons, and 36 modules contained all 12 columnar cell types but more than 12 neurons. Most of these 96 imperfect modules were at the boundary of the ME. Conversely, 113 neurons were over-assigned to modules (i.e., if for each module we picked a single assigned neuron per cell type, then 113 neurons were left over).

##### Hexagonal coordinate assignment and ROI creation

Column modules were assigned hexagonal coordinates by applying an algorithm developed to assign hexagonal coordinates to ommatidia of the fly eye^79^. This algorithm uses local difference vectors between a point and its 6 nearest neighbors (7 manually identified points) to find all the other neighbors and works reliably for nearly regularly spaced points. We applied this algorithm to the mean synapse positions of all columnar neurons in each module. We manually verified the assigned hexagonal coordinates and made ∼10 manual re-assignments to edge columns. We removed 5 edge modules (outside of the ME ROI or duplicated), resulting in 879 column modules with hexagonal coordinates. Column ROIs were created using one neuron per cell type per hexagonal coordinate, with the same algorithm (and identical parameters) used for the right OL^36^. To make layer ROIs, we also applied the same algorithm as for the right OL but with slightly adjusted parameters: in the ME, we used “alpha=0.0004” and “fac_ext=1”; in the LO, “frac_peaks=0.75”, “alpha=0.0002”, “fac_ext=0.3”; in the LOP, “frac_peaks=0.85”, “alpha=0.0002”, “fac_ext=0.3”.

##### Extending the coordinate system to the LO and LOP

We relaxed the criterion for T4-to-Mi1 assignments from requiring all four T4 types to requiring at least three types to be assigned to an Mi1 neuron. This increased the valid assignments in the right lobula plate. These results are summarized in Fig S4Bb. The number of layers on the left matches those on the right.

### Sex-specific branch analysis

For the analysis of connectivity on sex-specific branches (Fig 8l-n) we examined the ∼120 dimorphic cell types. 46 types (135 individual neurons) with well-defined extra axonal or dendritic branches in the male were split into the isomorphic and male-specific parts (Fig 8l). We manually placed annotations in Neuroglancer to define cut points that would separate sex-specific from isomorphic branches. We then split skeletons for each of the 46 types using `navis` (see Table 2) and associated their synaptic connections with either sex-specific or isomorphic branches using nearest-neighbor lookup of synapse locations to skeletons. All splits were manually inspected to confirm correctness. Depending on whether the sex-specific branches represented the neurons’ dendrites or axons, we only quantified their incoming or outgoing connections, respectively.

For each connection to or from these types (4,418 isomorphic and 1,856 dimorphic edges), we determined the number of synapses on male-specific versus isomorphic branches. On average, male-specific branches contained 22.2% of all dimorphic synapses and only 9.2% of all isomorphic synapses (Fig 8m). The mean for isomorphic synapses was skewed by a small number of types with large male-specific branches (e.g. SIP025) that contained a high fraction of isomorphic synapses. The median is 20% for dimorphic and only 4.8% for isomorphic synapses. Conversely, 61% of synapses on male-specific branches were dimorphic and 39% are isomorphic, compared to 39% and 61% on isomorphic branches, respectively (Fig 8n). Again, this mean was skewed by a small number of outliers: male-specific branches of DNg83, DNbe005 and DNge128 contain little to no dimorphic synapses, suggesting that their male-specific branches are either the result of biological variability or not the primary driver of dimorphic connectivity. Importantly, for 32 out of the 46 analysed types, the male-specific branches carry mostly dimorphic synapses. These findings suggest that sex-specific branches tend to be enriched in dimorphic connectivity, but the opposite is not necessarily true: dimorphic in- or outputs can be found across the entire neuron. Only a small fraction of dimorphic connections are localised exclusively to sex-specific branches (Fig S8l).

