## Supplementary figures and images for "Sexual dimorphism in the complete connectome of the *Drosophila* male central nervous system"

### Higher Resolution Main Figures 1-9

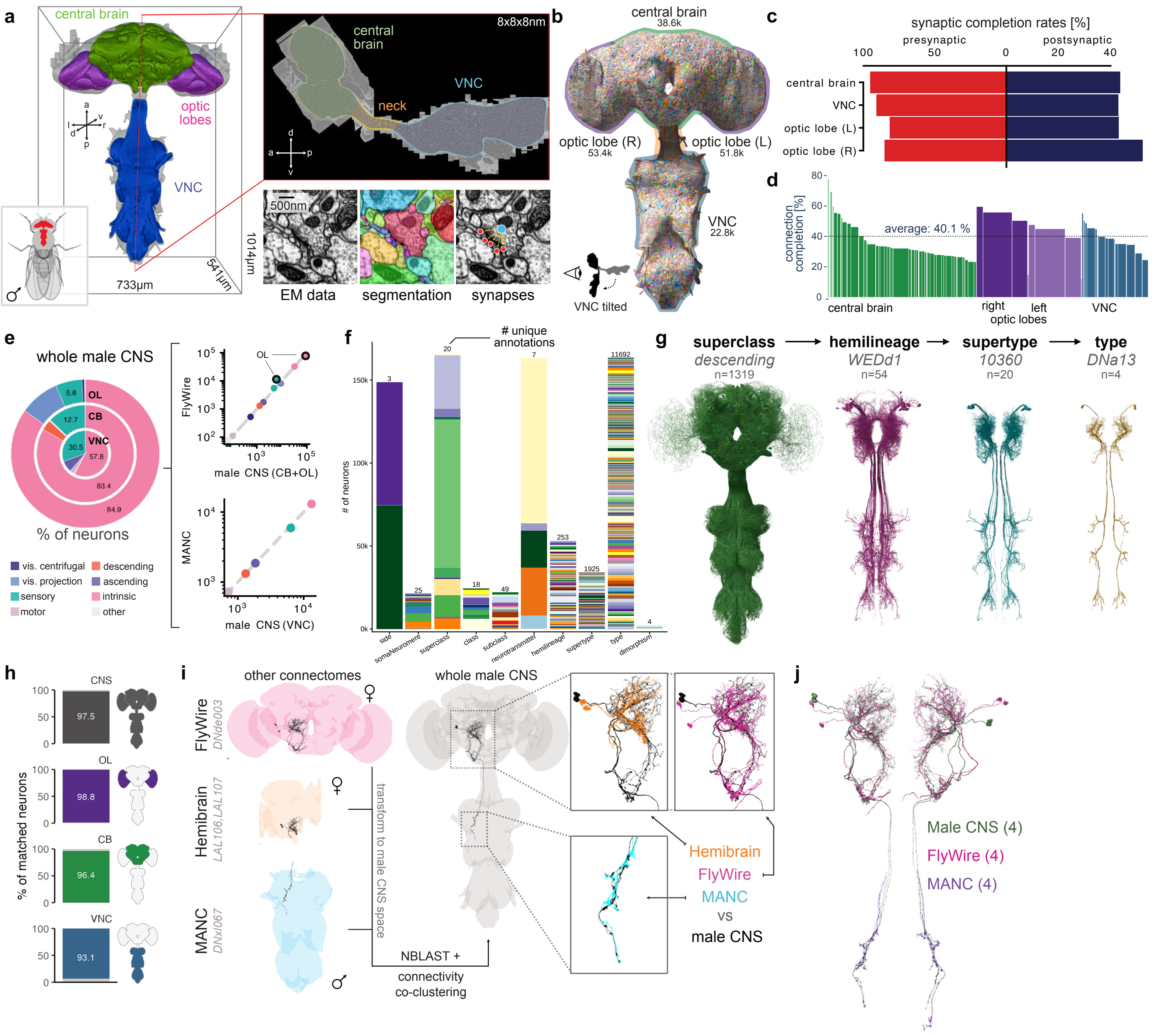

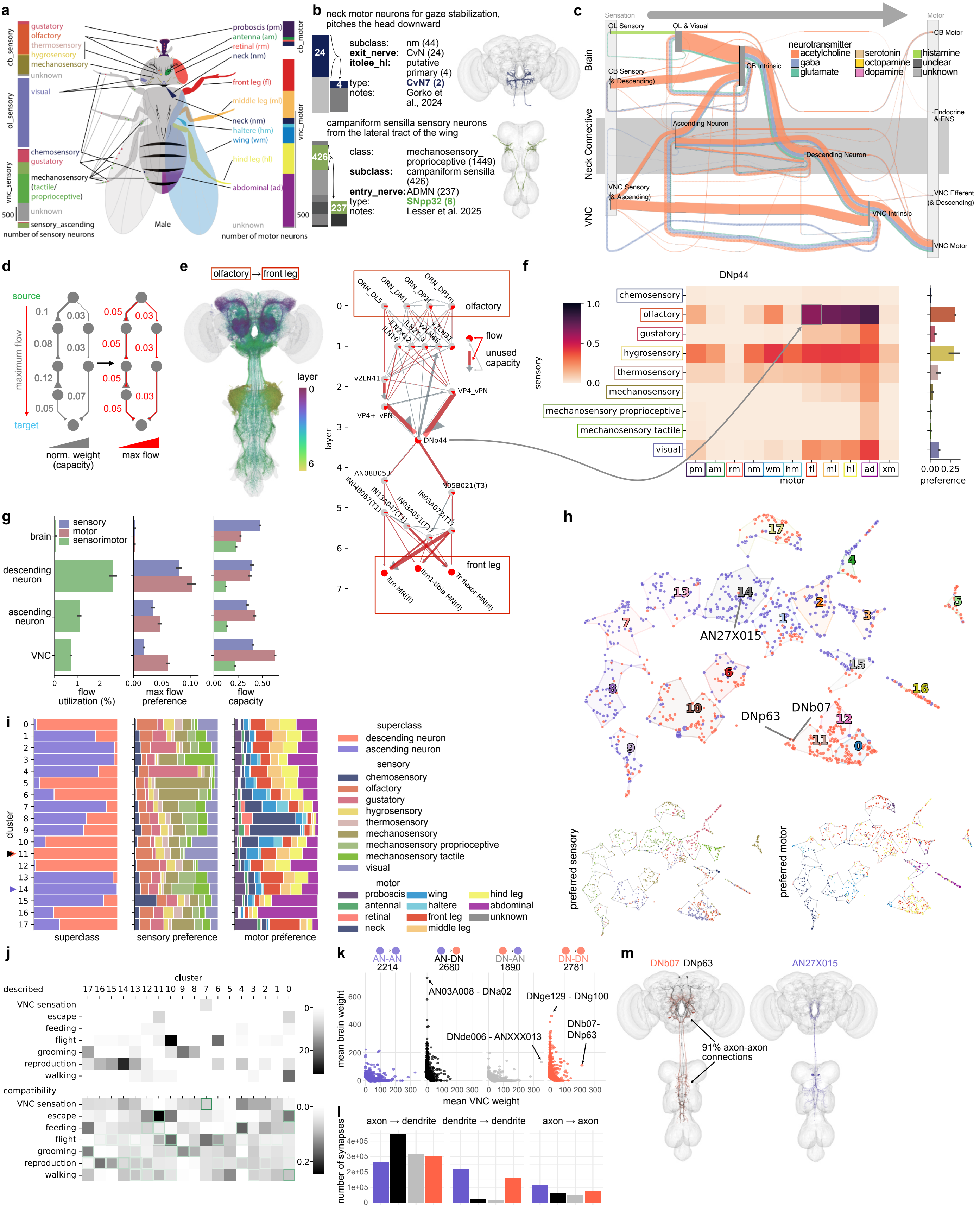

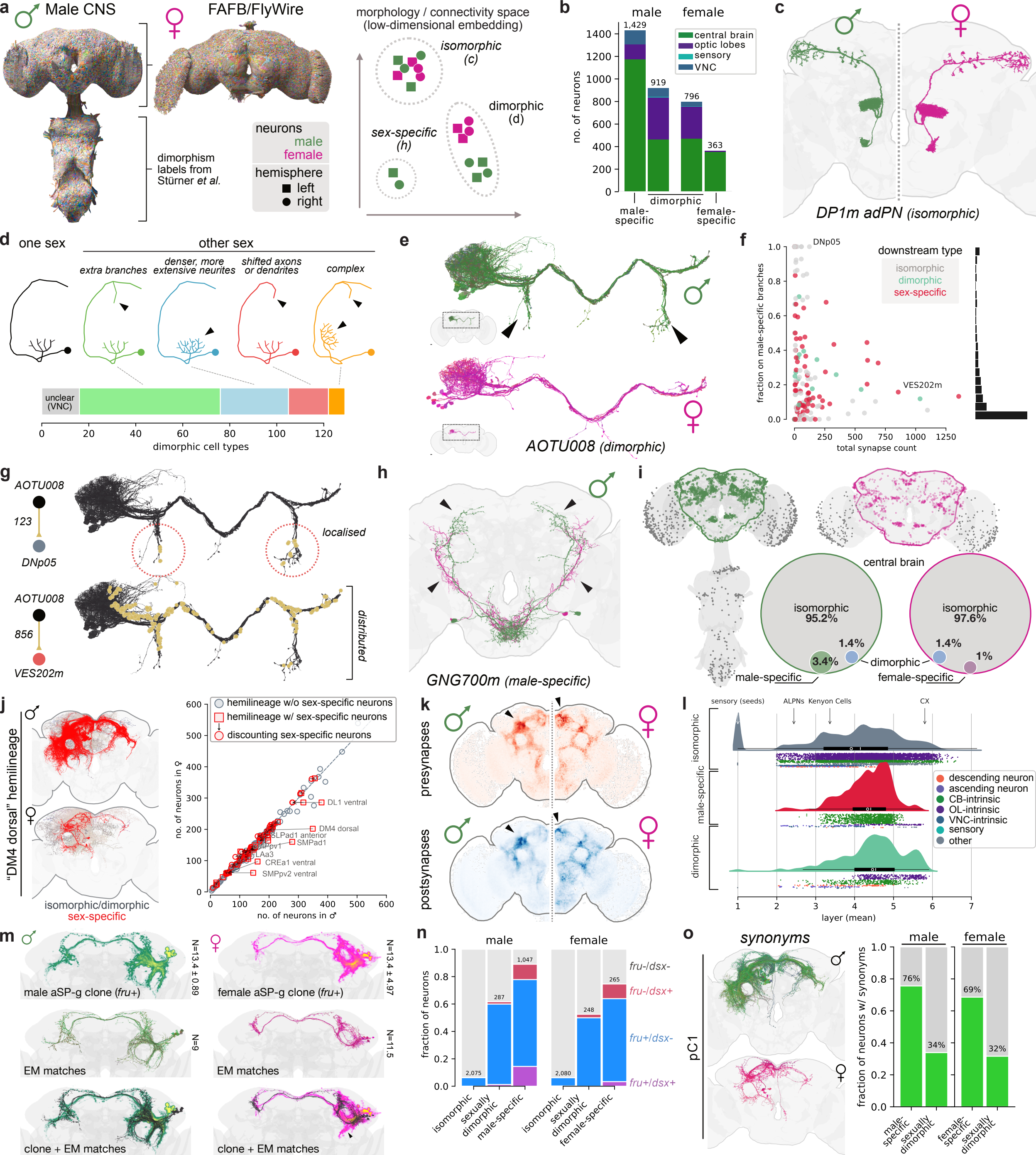

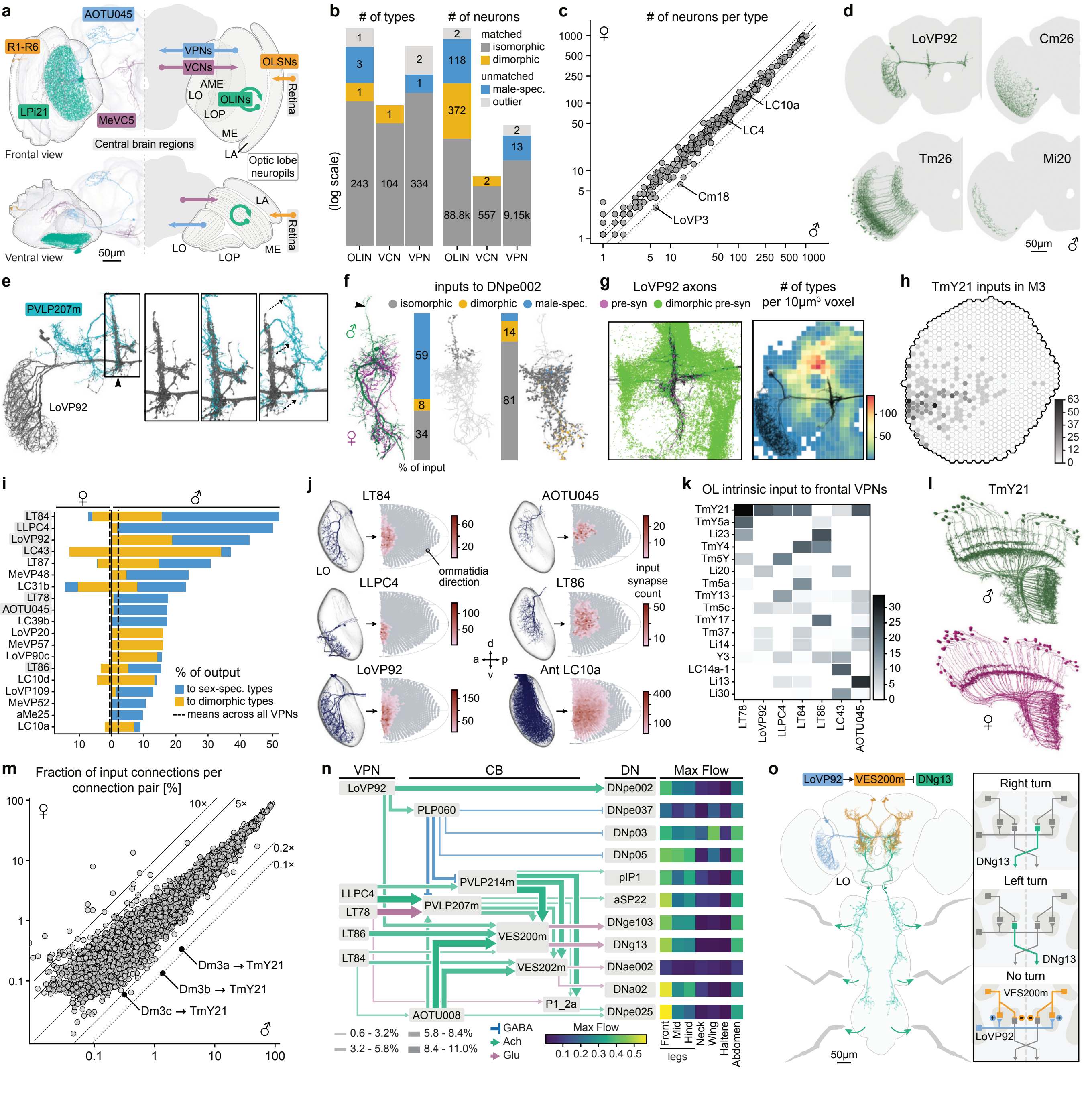

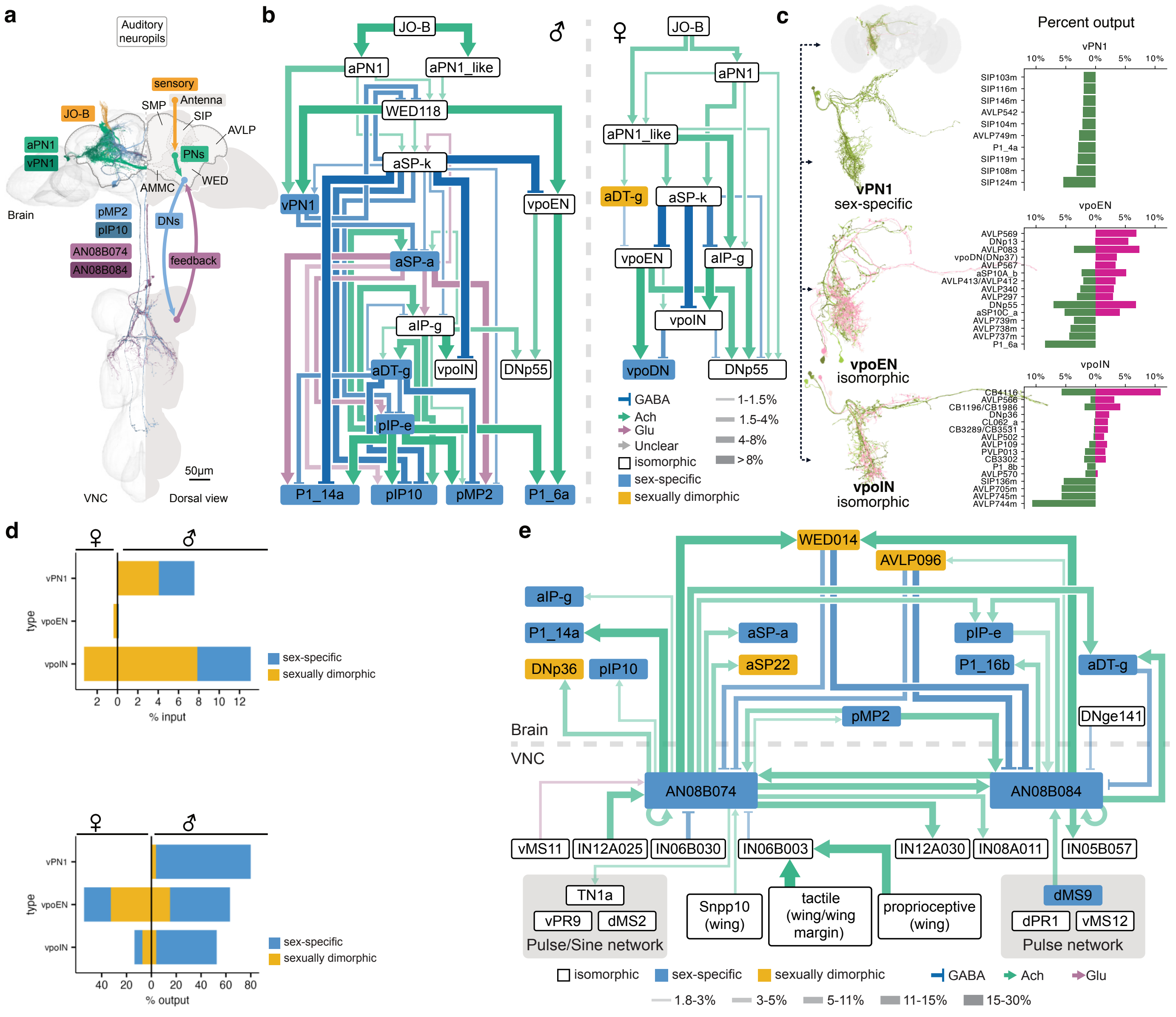

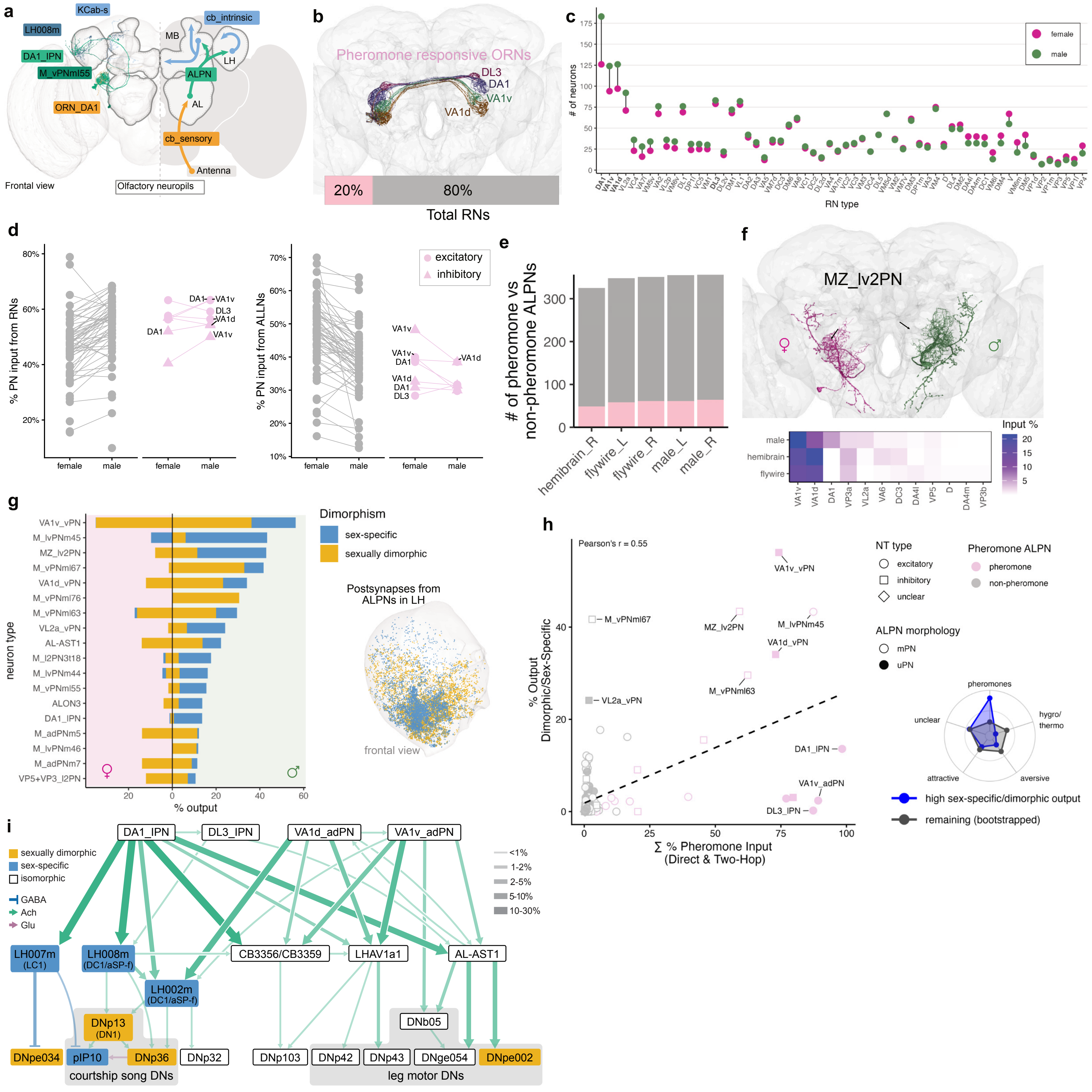

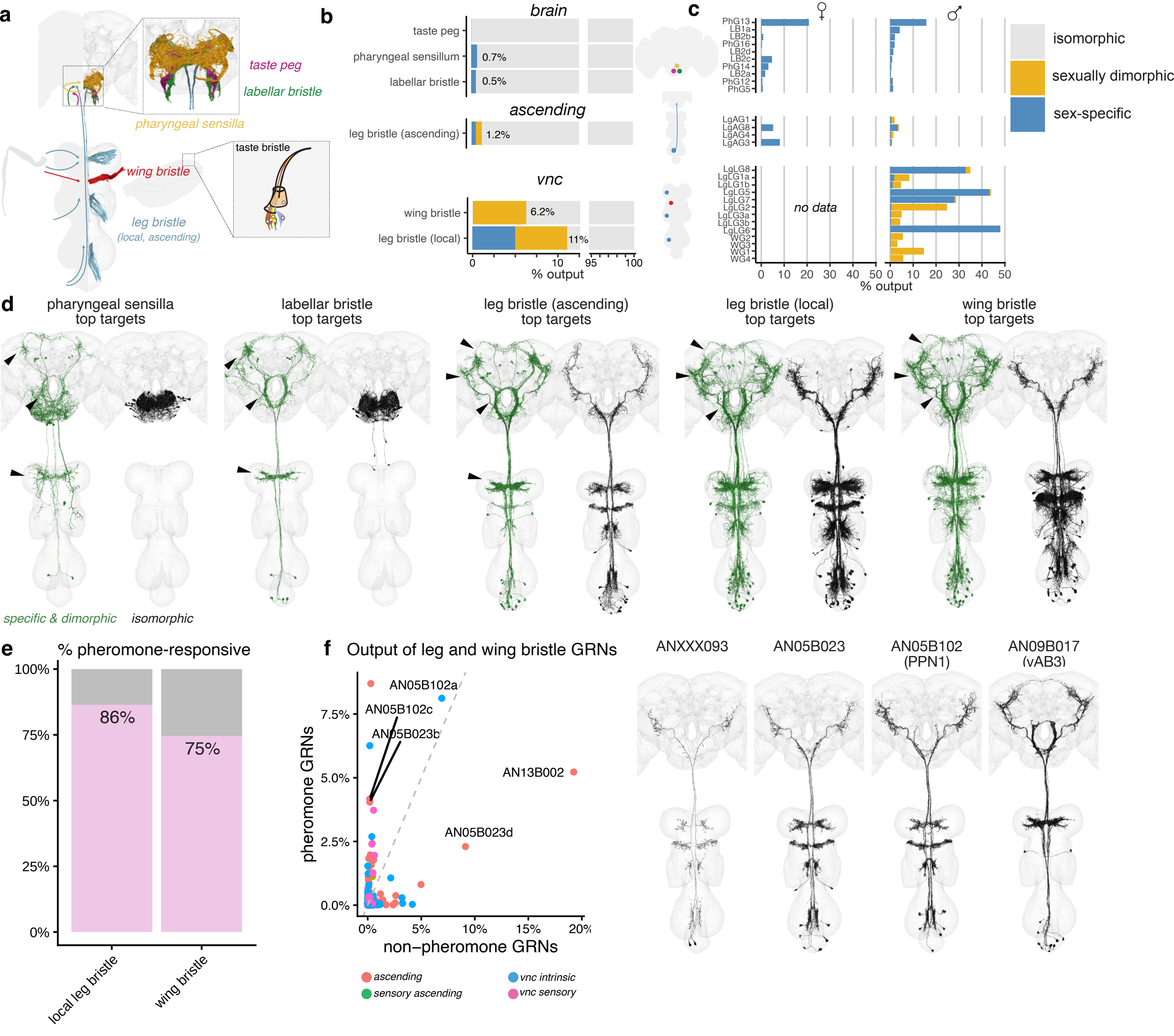

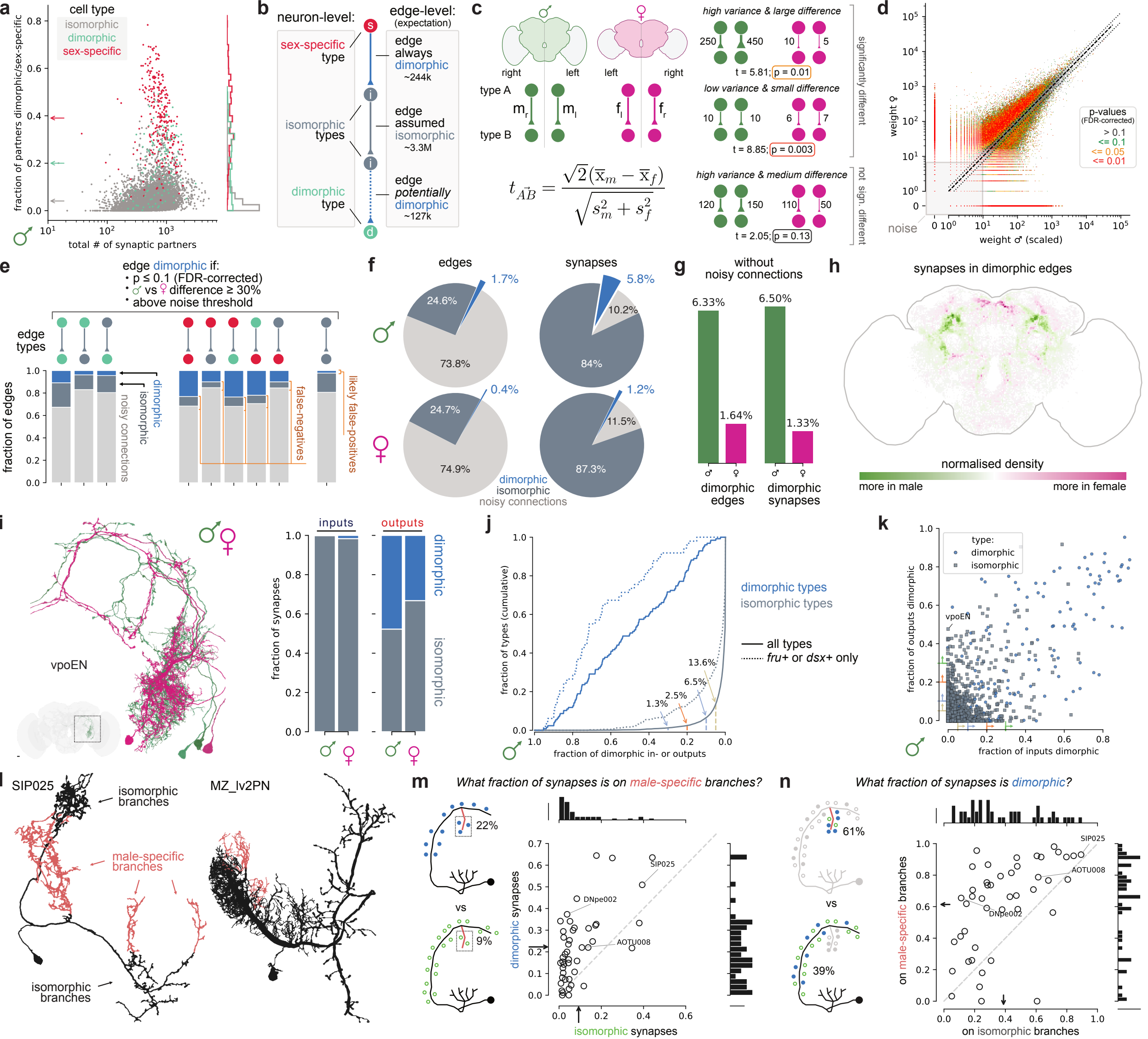

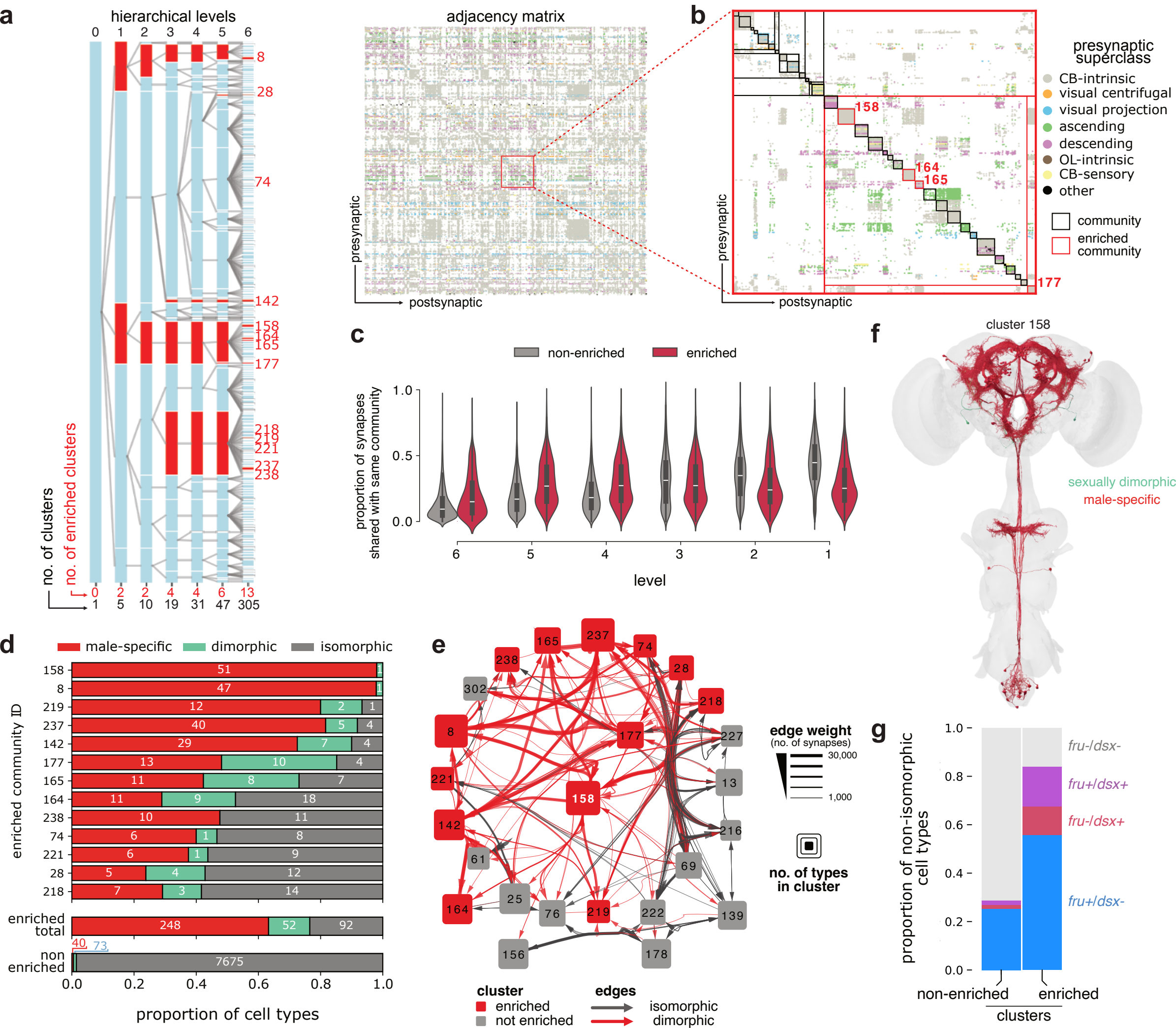
