## Supplementary material for "Sexual dimorphism in the complete connectome of the *Drosophila* male central nervous system": Higher Resolution Supplementary Figures S1-S9

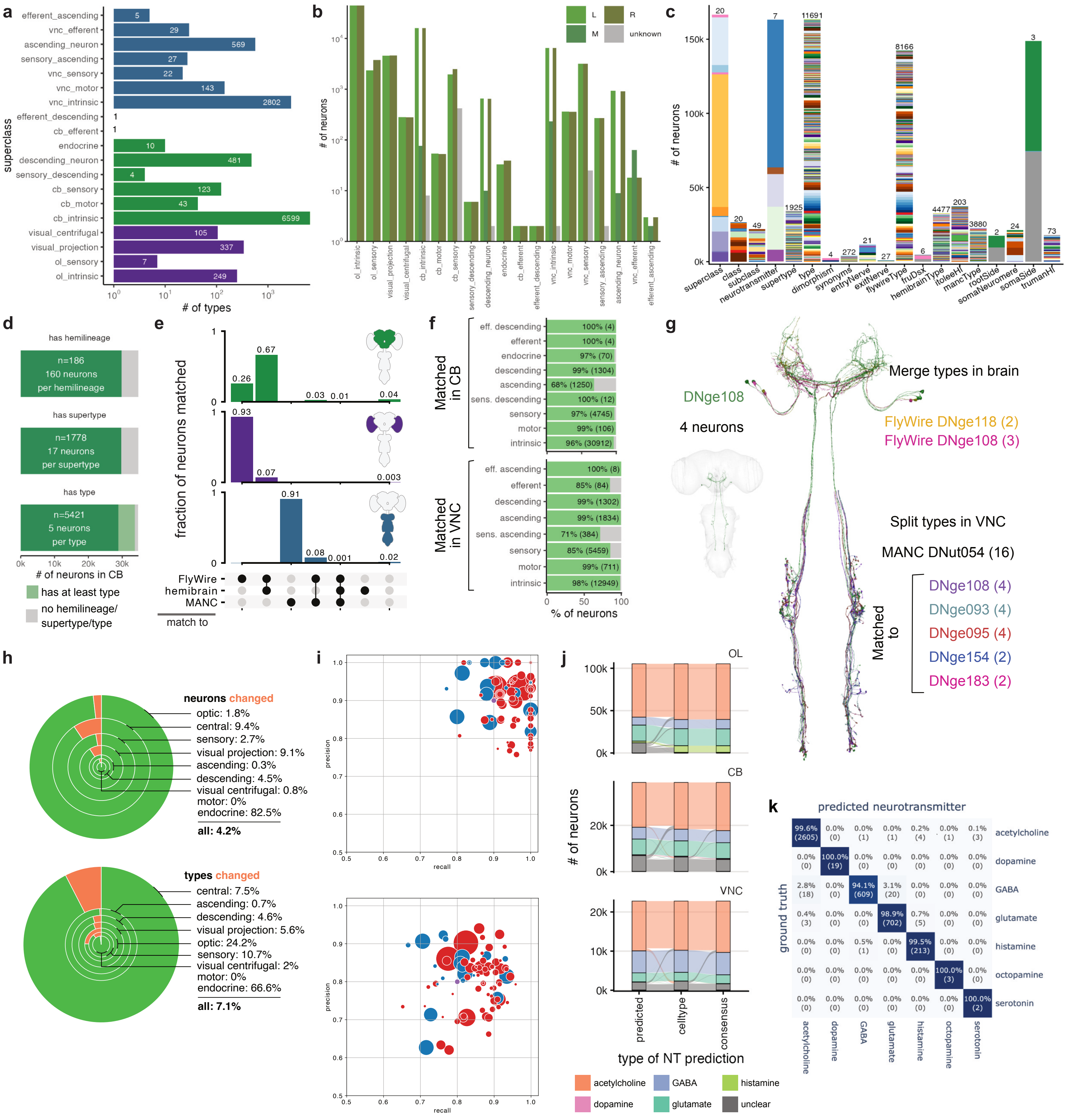

a
efferent_ascending 5
vnc_efferent 29
ascending_neuron 569
sensory_ascending 27
vnc_sensory 22
vnc_motor 143
vnc_intrinsic 2802
efferent_descending 1
cb_efferent 1
endocrine 10
descending_neuron 481
sensory_descending 4
cb_sensory 123
cb_motor 43
cb_intrinsic 6599
visual_centrifugal 105
visual_projection 337
ol_sensory 7
ol_intrinsic 249
superclass
### of types
b
L
R
M
unknown
### of neurons
ol_intrinsic
ol_sensory
visual_projection
visual_centrifugal
cb_intrinsic
cb_motor
cb_sensory
sensory_descending
descending_neuron
endocrine
cb_efferent
efferent_descending
vnc_intrinsic
vnc_motor
vnc_sensory
sensory_ascending
ascending_neuron
vnc_efferent
efferent_ascending
c
### of neurons
150k
100k
50k
0k
20
20
49
7
1925
11691
4
272
21
27
8166
6
4477
203
3880
2
24
3
73
superclass
class
subclass
neurotransmitter
supertype
type
dimorphism
synonyms
entryNerve
exitNerve
flywireType
fruDsx
hemibrainType
itoleeHl
mancType
rootSide
somaNeuromere
somaSide
trumanHl
d
has hemilineage
n=186
160 neurons
per hemilineage
has supertype
n=1778
17 neurons
per supertype
has type
n=5421
5 neurons
per type
0k
10k
20k
30k
### of neurons in CB
has at least type
no hemilineage/
supertype/type
e
fraction of neurons matched
0.26
0.67
0.03
0.01
0.04
0.93
0.07
0.003
0.91
0.08
0.001
0.02
FlyWire
hemibrain
MANC
match to
f
Matched in CB
eff. descending 100% (4)
efferent 100% (4)
endocrine 97% (70)
descending 99% (1304)
ascending 68% (1250)
sens. descending 100% (12)
sensory 97% (4745)
motor 99% (106)
intrinsic 96% (30912)
Matched in VNC
eff. ascending 100% (8)
efferent 85% (84)
descending 99% (1302)
ascending 99% (1834)
sens. ascending 71% (384)
sensory 85% (5459)
motor 99% (711)
intrinsic 98% (12949)
% of neurons
g
DNge108
4 neurons
Merge types in brain
FlyWire DNge118 (2)
FlyWire DNge108 (3)
Split types in VNC
MANC DNut054 (16)
Matched to
DNge108 (4)
DNge093 (4)
DNge095 (4)
DNge154 (2)
DNge183 (2)
h
neurons changed
optic: 1.8%
central: 9.4%
sensory: 2.7%
visual projection: 9.1%
ascending: 0.3%
descending: 4.5%
visual centrifugal: 0.8%
motor: 0%
endocrine: 82.5%
all: 4.2%
types changed
central: 7.5%
ascending: 0.7%
descending: 4.6%
visual projection: 5.6%
optic: 24.2%
sensory: 10.7%
visual centrifugal: 2%
motor: 0%
endocrine: 66.6%
all: 7.1%
i
precision
recall
j
OL
CB
VNC
### of neurons
predicted
celltype
consensus
type of NT prediction
acetylcholine
GABA
histamine
dopamine
glutamate
unclear
k
predicted neurotransmitter
ground truth
99.6% (2605) | 0.0% (0) | 0.0% (1) | 0.0% (1) | 0.2% (4) | 0.0% (1) | 0.1% (3) acetylcholine
0.0% (0) | 100.0% (19) | 0.0% (0) | 0.0% (0) | 0.0% (0) | 0.0% (0) | 0.0% (0) dopamine
2.8% (18) | 0.0% (0) | 94.1% (609) | 3.1% (20) | 0.0% (0) | 0.0% (0) | 0.0% (0) GABA
0.4% (3) | 0.0% (0) | 0.0% (0) | 98.9% (702) | 0.7% (5) | 0.0% (0) | 0.0% (0) glutamate
0.0% (0) | 0.0% (0) | 0.5% (1) | 0.0% (0) | 99.5% (213) | 0.0% (0) | 0.0% (0) histamine
0.0% (0) | 0.0% (0) | 0.0% (0) | 0.0% (0) | 0.0% (0) | 100.0% (3) | 0.0% (0) octopamine
0.0% (0) | 0.0% (0) | 0.0% (0) | 0.0% (0) | 0.0% (0) | 0.0% (0) | 100.0% (2) serotonin
acetylcholine
dopamine
GABA
glutamate
histamine
octopamine
serotonin

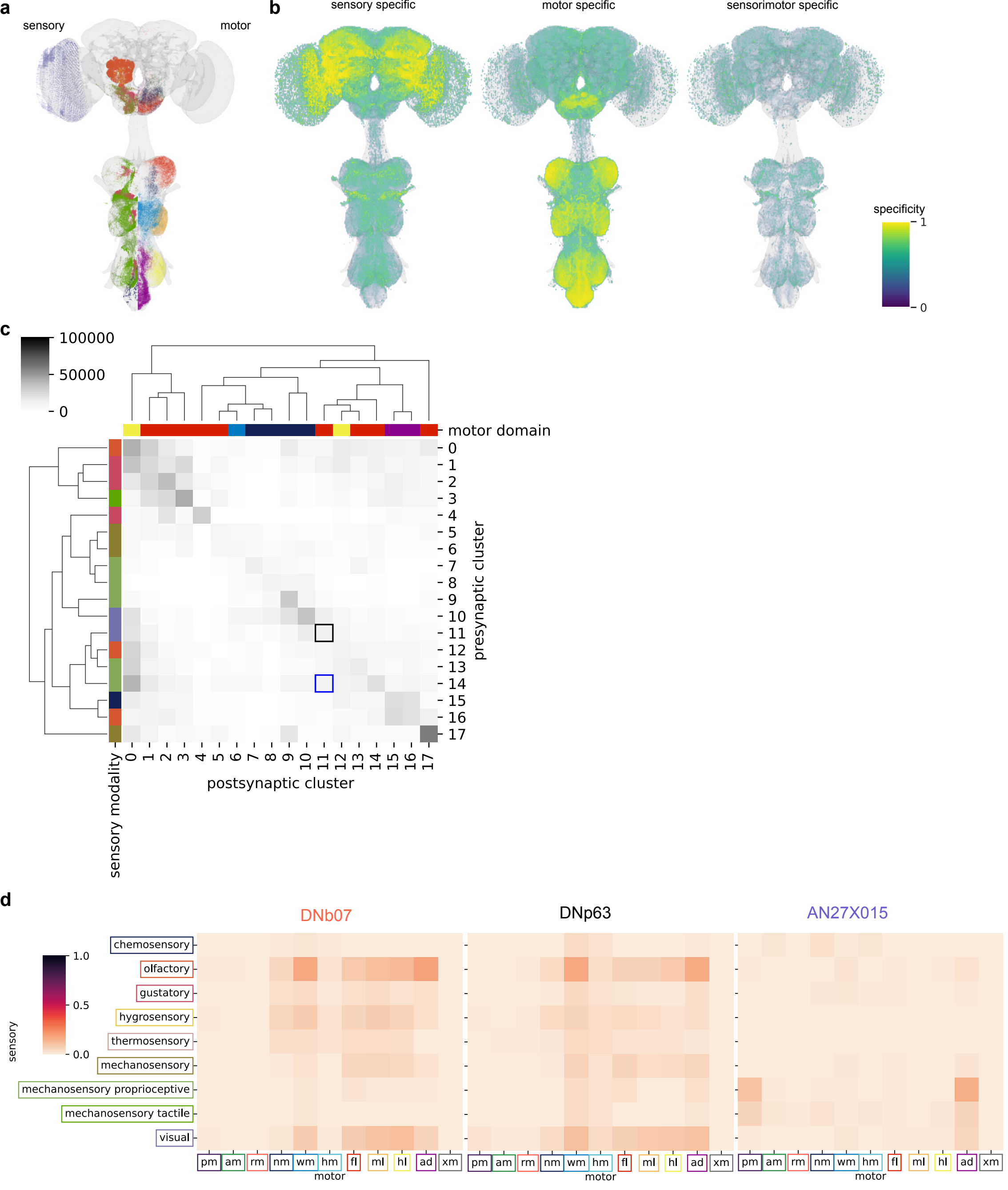
a
sensory
motor
b
sensory specific
motor specific
sensorimotor specific
specificity
1
0
c
100000
50000
0
motor domain
presynaptic cluster
postsynaptic cluster
sensory modality
d
DNb07
DNp63
AN27X015
1.0
0.5
0.0
sensory
chemosensory
olfactory
gustatory
hygrosensory
thermosensory
mechanosensory
mechanosensory proprioceptive
mechanosensory tactile
visual
pm
am
rm
nm
wm
hm
fl
ml
hl
ad
xm
motor

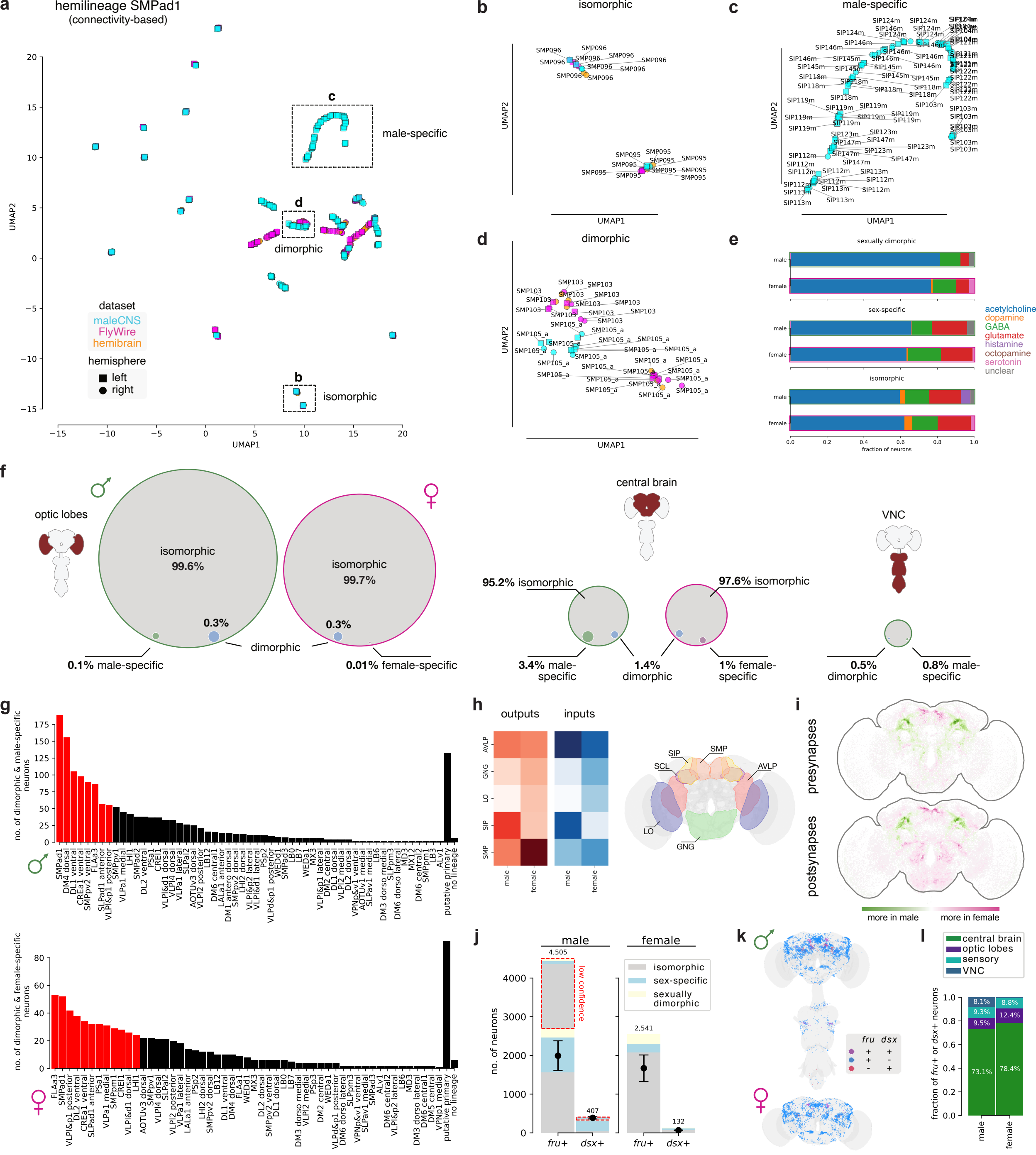

hemilineage SMPad1
(connectivity-based)
male-specific
dimorphic
isomorphic
dataset
maleCNS
FlyWire
hemibrain
hemisphere
left
right
UMAP1
UMAP2
sexually dimorphic
sex-specific
acetylcholine
dopamine
GABA
glutamate
histamine
octopamine
serotonin
unclear
fraction of neurons
optic lobes
central brain
VNC
isomorphic 99.6%
isomorphic 99.7%
0.1% male-specific
0.01% female-specific
95.2% isomorphic
97.6% isomorphic
3.4% male-specific
1.4% dimorphic
1% female-specific
0.5% dimorphic
0.8% male-specific
no. of dimorphic & male-specific neurons
no. of dimorphic & female-specific neurons
outputs
inputs
presynapses
postsynapses
more in male
more in female
no. of neurons
low confidence
sex-specific
sexually dimorphic
central brain
optic lobes
sensory
VNC
fraction of fru+ or dsx+ neurons

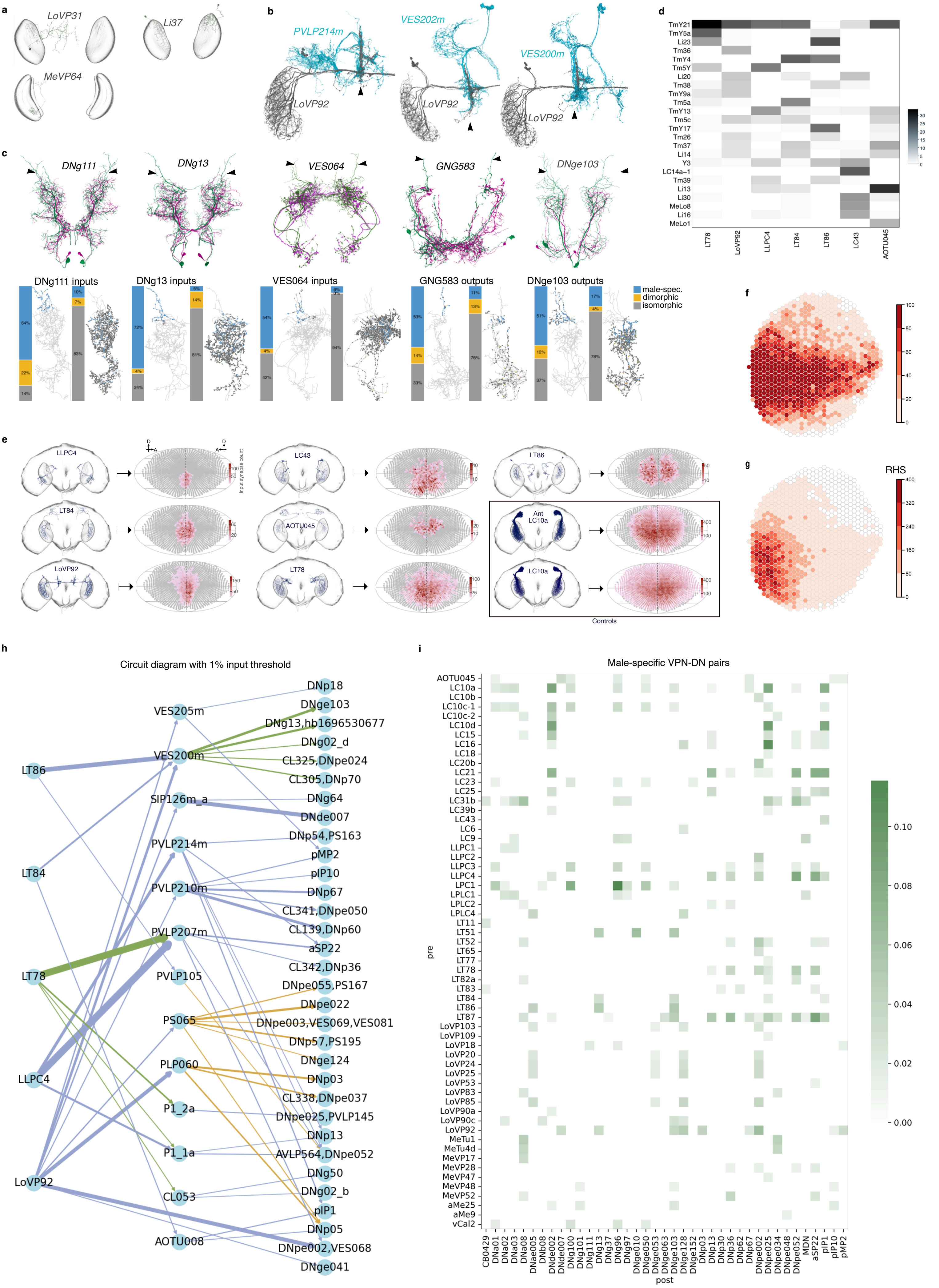
a
LoVP31
Li37
MeVP64
b
PVLP214m
VES202m
VES200m
LoVP92
c
DNg111
DNg13
VES064
GNG583
DNge103
DNg111 inputs
DNg13 inputs
VES064 inputs
GNG583 outputs
DNge103 outputs
male-spec.
dimorphic
isomorphic
d
TmY21
TmY5a
Li23
Tm36
TmY4
Tm5Y
Li20
Tm38
TmY9a
Tm5a
TmY13
Tm5c
TmY17
Tm26
Tm37
Li14
Y3
LC14a−1
Tm39
Li13
Li30
MeLo8
Li16
MeLo1
LT78
LoVP92
LLPC4
LT84
LT86
LC43
AOTU045
e
LLPC4
LC43
LT86
LT84
AOTU045
Ant LC10a
LoVP92
LT78
LC10a
Input synapse count
Controls
f
g
RHS
h
Circuit diagram with 1% input threshold
LT86
LT84
LT78
LLPC4
LoVP92
VES205m
VES200m
SIP126m_a
PVLP214m
PVLP210m
PVLP207m
PVLP105
PS065
PLP060
P1_2a
P1_1a
CL053
AOTU008
DNp18
DNge103
DNg13,hb1696530677
DNg02_d
CL325,DNpe024
CL305,DNp70
DNg64
DNde007
DNp54,PS163
pMP2
pIP10
DNp67
CL341,DNpe050
CL139,DNp60
aSP22
CL342,DNp36
DNpe055,PS167
DNpe022
DNpe003,VES069,VES081
DNp57,PS195
DNge124
DNp03
CL338,DNpe037
DNpe025,PVLP145
DNp13
AVLP564,DNpe052
DNg50
DNg02_b
pIP1
DNp05
DNpe002,VES068
DNge041
i
Male-specific VPN-DN pairs
pre
post
AOTU045
LC10a
LC10b
LC10c-1
LC10c-2
LC10d
LC15
LC16
LC18
LC20b
LC21
LC23
LC25
LC31b
LC39b
LC43
LC6
LC9
LLPC1
LLPC2
LLPC3
LLPC4
LPC1
LPLC1
LPLC2
LPLC4
LT11
LT51
LT52
LT65
LT77
LT78
LT82a
LT83
LT84
LT86
LT87
LoVP103
LoVP109
LoVP18
LoVP20
LoVP24
LoVP25
LoVP53
LoVP83
LoVP85
LoVP90a
LoVP90c
LoVP92
MeTu1
MeTu4d
MeVP17
MeVP28
MeVP47
MeVP48
MeVP52
aMe25
aMe9
vCal2
CB0429
DNa01
DNa02
DNa03
DNa08
DNae005
DNb08
DNde002
DNde007
DNg100
DNg101
DNg111
DNg13
DNg37
DNg96
DNg97
DNge010
DNge050
DNge053
DNge063
DNge103
DNge128
DNge152
DNp03
DNp13
DNp30
DNp36
DNp62
DNp67
DNpe002
DNpe025
DNpe034
DNpe048
DNpe052
MDN
aSP22
pIP1
pIP10
pMP2
0.10
0.08
0.06
0.04
0.02
0.00

**a**

| groups | # cell types R | # cell types L | # cells R | # cells L | # input conn R | # input conn L | # output conn R | # output conn L |
|---|---|---|---|---|---|---|---|---|
| ONIN | 149 | 149 | 15.7K | 14.4K | 8.3M | 7.3M | 15.3M | 15.4M |
| ONCN | 95 | 95 | 32.4K | 32.3K | 13.9M | 11.6M | 27.6M | 28.1M |
| VPN | 352 | 348 | 4.5K | 4.5K | 6.0M | 5.1M | 9.9M | 10.1M |
| VCN | 104 | 104 | 267 | 273 | 776.1K | 757.2K | 2.5M | 2.7M |

**b**

| neuropil | # cell types R | # cell types L | # cells R | # cells L | # input conn R | # input conn L | # output conn R | # output conn L |
|---|---|---|---|---|---|---|---|---|
| LA | 15 | 22 | 6.9K | 4.5K | 395.1K | 272.9K | 529.7K | 295.3K |
| ME | 345 | 366 | 41.9K | 41.8K | 15.5M | 13.3M | 26.7M | 27.0M |
| LO | 422 | 430 | 25.9K | 25.9K | 7.4M | 6.1M | 14.1M | 14.4M |
| LOP | 128 | 125 | 13.1K | 12.7K | 3.7M | 3.0M | 6.0M | 5.8M |
| AME | 60 | 25 | 137 | 83 | 26.8K | 7.2K | 46.9K | 15.1K |

**c**

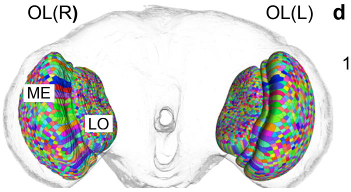


| neuropil | # column rois R | # column rois L |
|---|---|---|
| ME | 902 | 890 |
| LO | 882 | 874 |
| LOP | 860 | 846 |

**d**

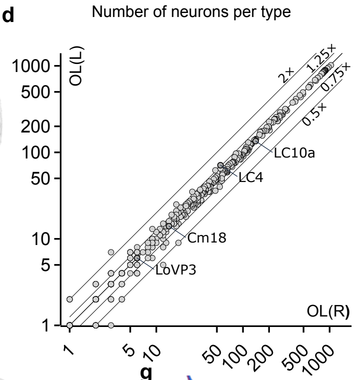


**e**

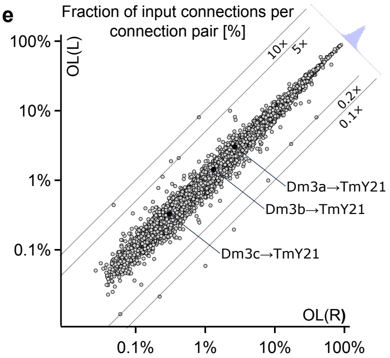


**f**

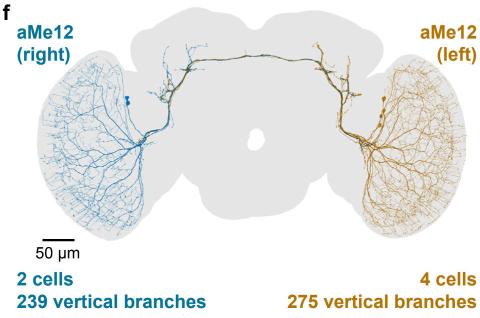


**g**

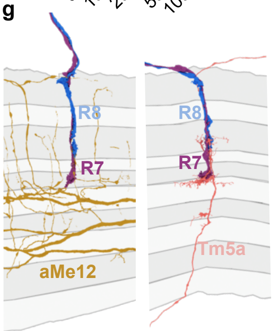


**h**

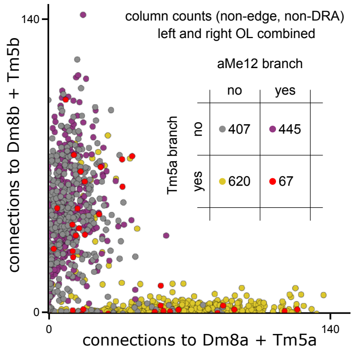


**i**

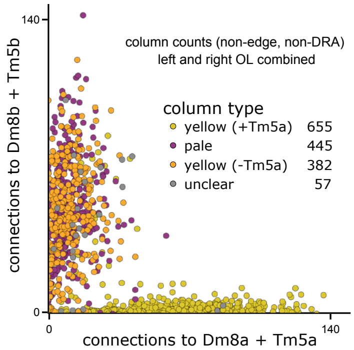


**j**

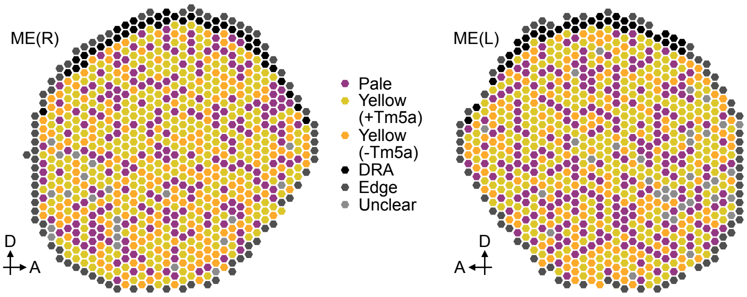

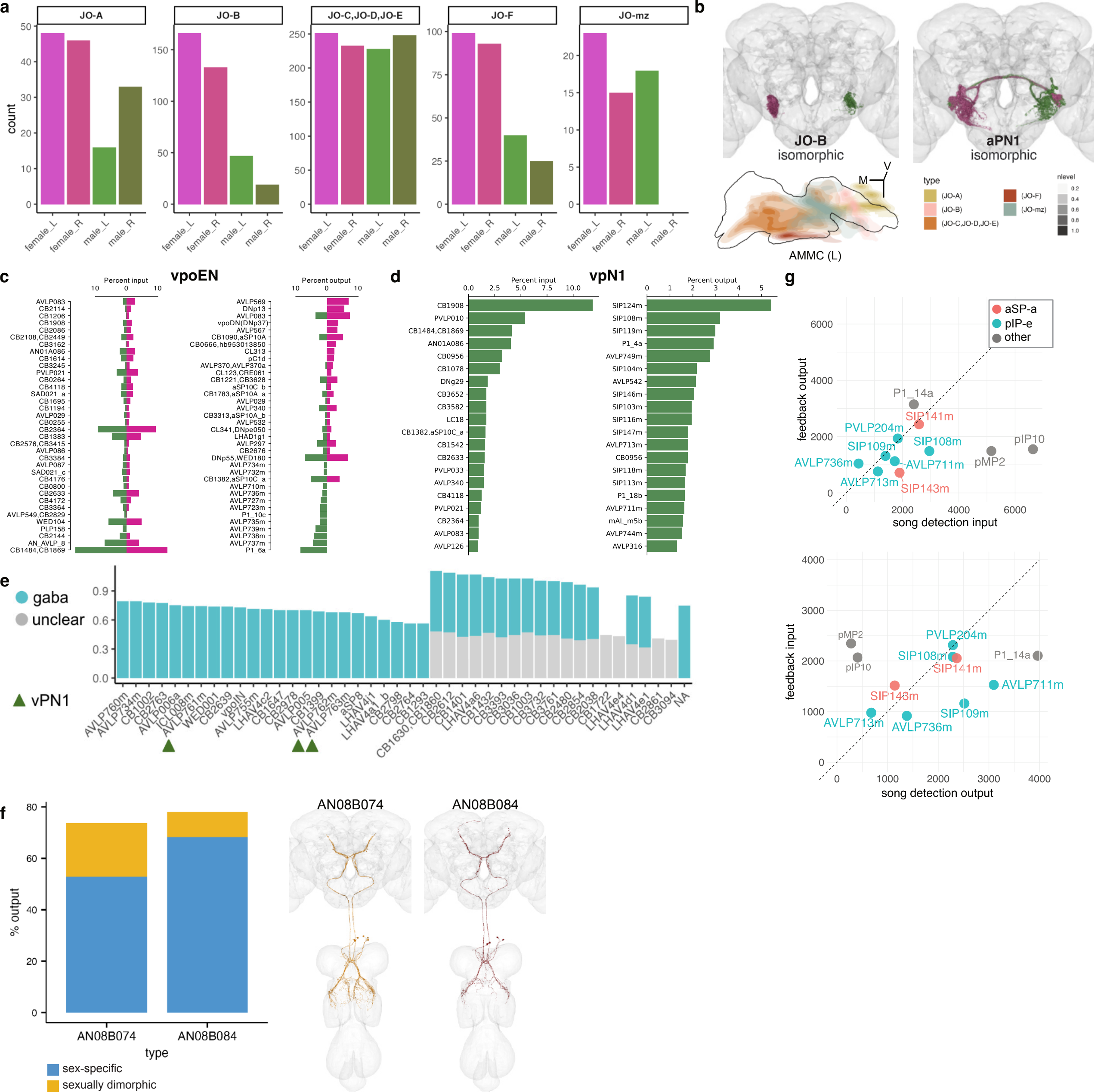

a
JO-A
JO-B
JO-C,JO-D,JO-E
JO-F
JO-mz
count
female_L
female_R
male_L
male_R
b
JO-B
isomorphic
aPN1
isomorphic
AMMC (L)
M
V
type
(JO-A)
(JO-B)
(JO-C,JO-D,JO-E)
(JO-F)
(JO-mz)
nlevel
c
vpoEN
Percent input
Percent output
d
vpN1
Percent input
Percent output
CB1908
PVLP010
CB1484,CB1869
AN01A086
CB0956
CB1078
DNg29
CB3652
CB3582
LC18
CB1382,aSP10C_a
CB1542
CB2633
PVLP033
AVLP340
CB4118
PVLP021
CB2364
AVLP083
AVLP126
SIP124m
SIP108m
SIP119m
P1_4a
AVLP749m
SIP104m
AVLP542
SIP146m
SIP103m
SIP116m
SIP147m
AVLP713m
CB0956
SIP118m
SIP113m
P1_18b
AVLP711m
mAL_m5b
AVLP744m
AVLP316
e
gaba
unclear
vPN1
f
% output
type
AN08B074
AN08B084
sex-specific
sexually dimorphic
g
aSP-a
pIP-e
other
feedback output
song detection input
feedback input
song detection output
P1_14a
SIP141m
PVLP204m
SIP109m
SIP108m
AVLP736m
AVLP711m
pMP2
pIP10
AVLP713m
SIP143m

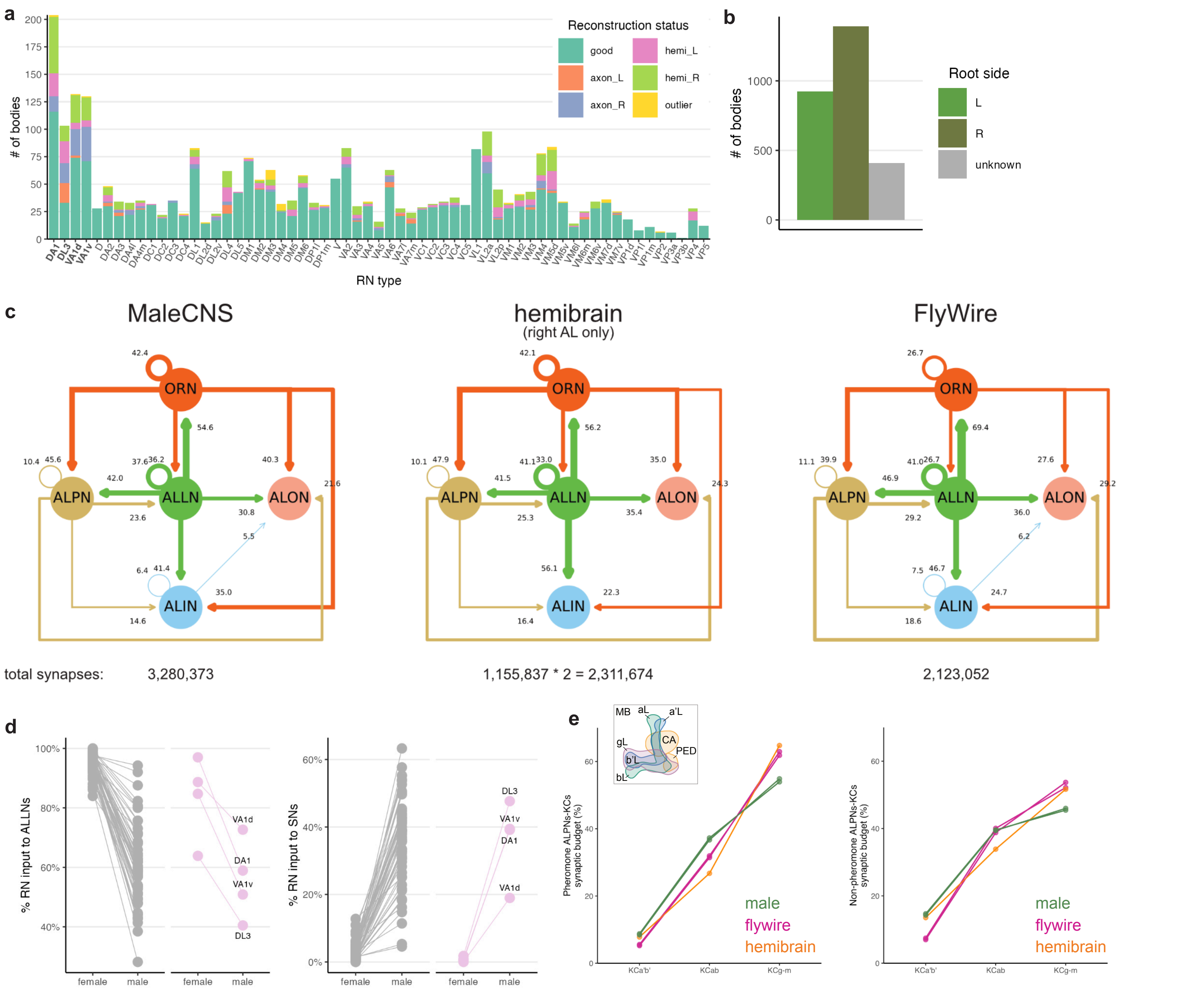

a
Reconstruction status
good
axon_L
axon_R
hemi_L
hemi_R
outlier
### of bodies
RN type
b
Root side
L
R
unknown
### of bodies
c
MaleCNS
hemibrain
(right AL only)
FlyWire
ORN
ALPN
ALLN
ALON
ALIN
total synapses:
3,280,373
1,155,837 * 2 = 2,311,674
2,123,052
d
% RN input to ALLNs
% RN input to SNs
female
male
VA1d
DA1
VA1v
DL3
e
MB
aL
a'L
gL
CA
PED
b'L
bL
Pheromone ALPNs-KCs synaptic budget (%)
Non-pheromone ALPNs-KCs synaptic budget (%)
KCa'b'
KCab
KCg-m
male
flywire
hemibrain

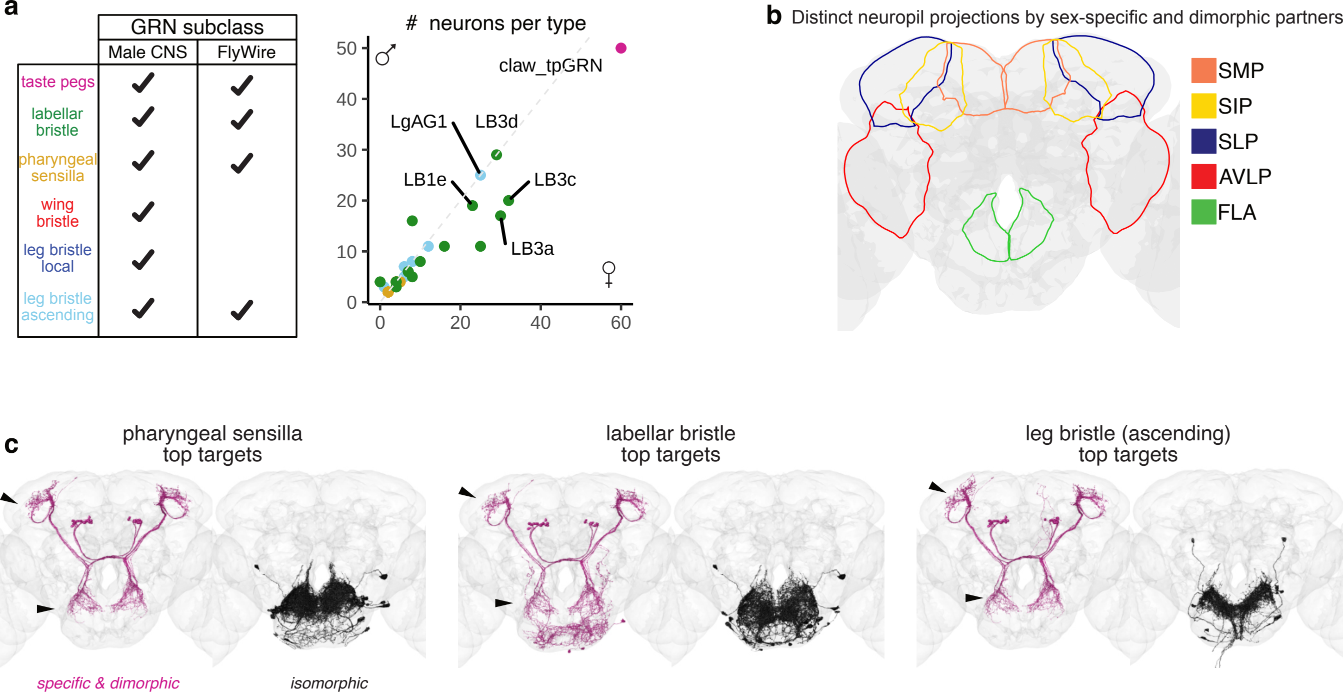


| GRN subclass | Male CNS | FlyWire |
|---|---|---|
| taste pegs | ✓ | ✓ |
| labellar bristle | ✓ | ✓ |
| pharyngeal sensilla | ✓ | ✓ |
| wing bristle | ✓ | |
| leg bristle local | ✓ | |
| leg bristle ascending | ✓ | ✓ |

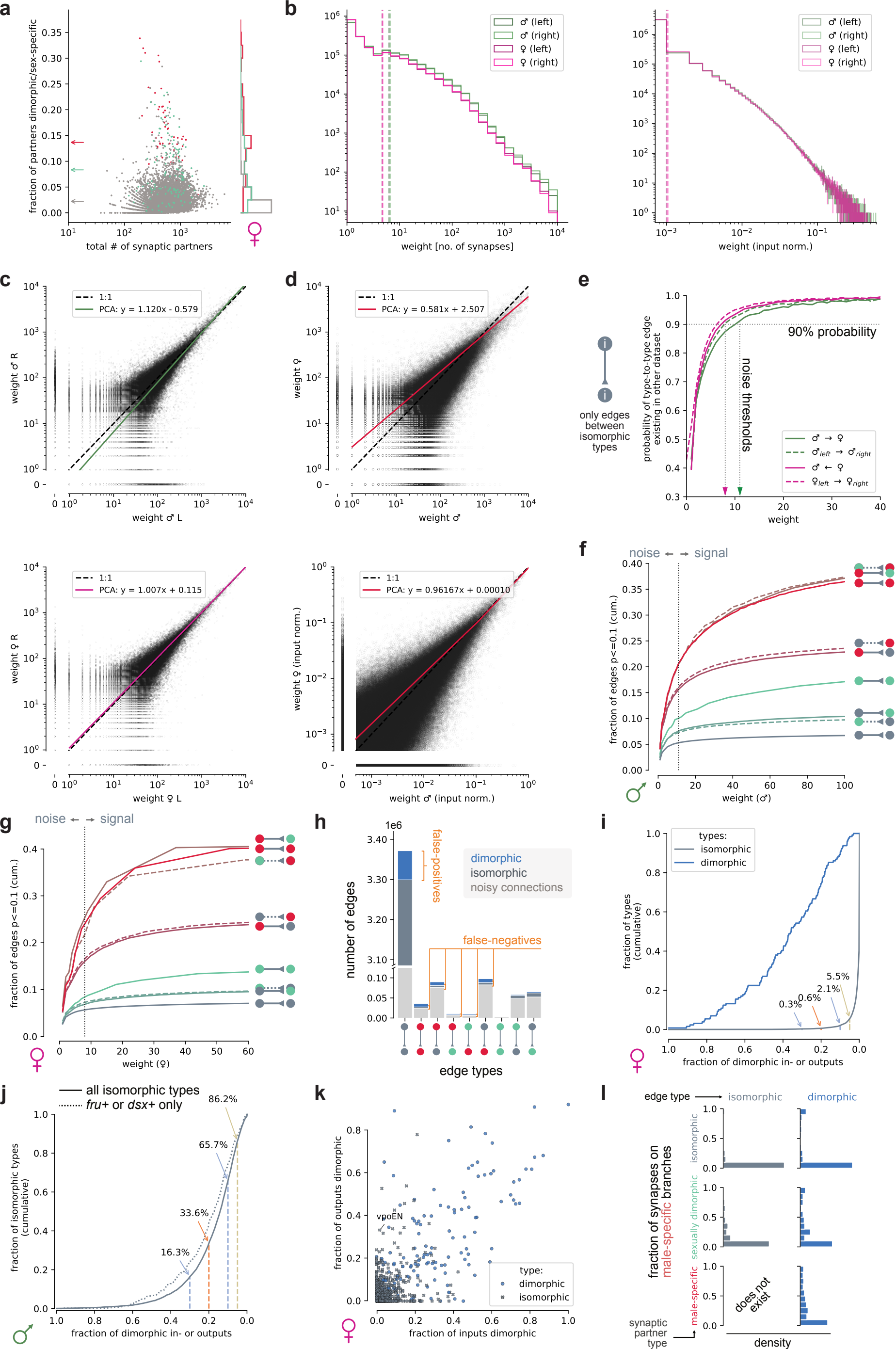
a
fraction of partners dimorphic/sex-specific
total # of synaptic partners
b
♂ (left)
♂ (right)
♀ (left)
♀ (right)
weight [no. of synapses]
weight (input norm.)
c
1:1
PCA: y = 1.120x - 0.579
weight ♂ R
weight ♂ L
PCA: y = 1.007x + 0.115
weight ♀ R
weight ♀ L
d
PCA: y = 0.581x + 2.507
weight ♀
weight ♂
PCA: y = 0.96167x + 0.00010
weight ♀ (input norm.)
weight ♂ (input norm.)
e
only edges between isomorphic types
probability of type-to-type edge existing in other dataset
90% probability
noise thresholds
weight
f
noise ← → signal
fraction of edges p<=0.1 (cum.)
weight (♂)
g
noise ← → signal
fraction of edges p<=0.1 (cum.)
weight (♀)
h
1e6
false-positives
false-negatives
dimorphic
isomorphic
noisy connections
number of edges
edge types
i
types:
isomorphic
dimorphic
fraction of types (cumulative)
5.5%
2.1%
0.3%
0.6%
fraction of dimorphic in- or outputs
j
all isomorphic types
fru+ or dsx+ only
86.2%
65.7%
33.6%
16.3%
fraction of isomorphic types (cumulative)
fraction of dimorphic in- or outputs
k
vpoEN
fraction of outputs dimorphic
fraction of inputs dimorphic
type:
dimorphic
isomorphic
l
edge type
isomorphic
dimorphic
fraction of synapses on male-specific branches
isomorphic
sexually dimorphic
male-specific
does not exist
synaptic partner type
density

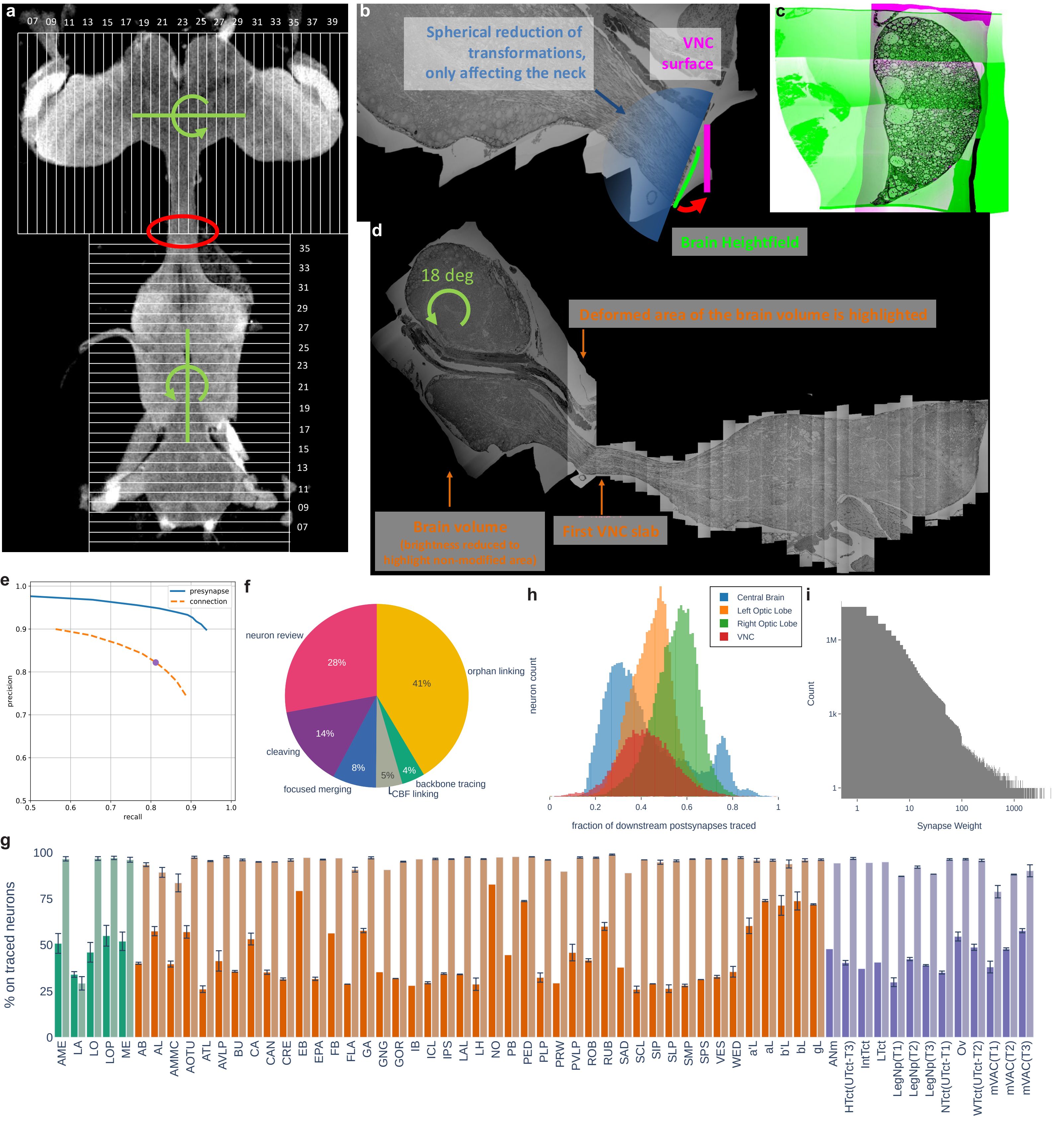

a
07 09 11 13 15 17 19 21 23 25 27 29 31 33 35 37 39
35 33 31 29 27 25 23 21 19 17 15 13 11 09 07
b
Spherical reduction of transformations, only affecting the neck
VNC surface
Brain Heightfield
c
d
18 deg
Deformed area of the brain volume is highlighted
Brain volume
(brightness reduced to highlight non-modified area)
First VNC slab
e
presynapse
connection
precision
recall
f
neuron review
28%
orphan linking
41%
cleaving
14%
focused merging
8%
5%
4%
backbone tracing
CBF linking
h
Central Brain
Left Optic Lobe
Right Optic Lobe
VNC
neuron count
fraction of downstream postsynapses traced
i
Count
Synapse Weight
g
% on traced neurons
AME LA LO LOP ME AB AL AMMC AOTU ATL AVLP BU CA CAN CRE EB EPA FB FLA GA GNG GOR IB ICL IPS LAL LH NO PB PED PLP PRW PVLP ROB RUB SAD SCL SIP SLP SMP SPS VES WED a'L aL b'L bL gL ANm HTct(UTct-T3) IntTct LTct LegNp(T1) LegNp(T2) LegNp(T3) NTct(UTct-T1) Ov WTct(UTct-T2) mVAC(T1) mVAC(T2) mVAC(T3)
